# Discovery of Antiviral Cyclic Peptides Targeting the Main Protease of SARS-CoV-2 *via* mRNA Display

**DOI:** 10.1101/2021.08.23.457419

**Authors:** Jason Johansen-Leete, Sven Ullrich, Sarah E. Fry, Rebecca Frkic, Max J. Bedding, Anupriya Aggarwal, Anneliese S. Ashhurst, Kasuni B. Ekanayake, Mithun C. Mahawaththa, Vishnu M. Sasi, Toby Passioura, Mark Larance, Gottfried Otting, Stuart Turville, Colin J. Jackson, Christoph Nitsche, Richard J. Payne

## Abstract

Antivirals that specifically target SARS-CoV-2 are needed to control the COVID-19 pandemic. The main protease (M^pro^) is essential for SARS-CoV-2 replication and is an attractive target for antiviral development. Here we report the use of the Random nonstandard Peptide Integrated Discovery (RaPID) mRNA display on a chemically cross-linked SARS-CoV-2 M^pro^ dimer, which yielded several high-affinity thioether-linked cyclic peptide inhibitors of the protease. Structural analysis of M^pro^ complexed with a selenoether analogue of the highest-affinity peptide revealed key binding interactions, including glutamine and leucine residues in sites S_1_ and S_2_, respectively, and a binding epitope straddling both protein chains in the physiological dimer. Several of these M^pro^ peptide inhibitors possessed antiviral activity against SARS-CoV-2 *in vitro* with EC_50_ values in the low micromolar range. These cyclic peptides serve as a foundation for the development of much needed antivirals that specifically target SARS-CoV-2.

## Introduction

The COVID-19 pandemic, caused by infection with severe acute respiratory syndrome coronavirus 2 (SARS-CoV-2) has caused widespread morbidity and mortality as well as devastation to the global economy since the disease was first reported in late 2019 in Wuhan, China^1^. At the time of writing there has been more than 200 million confirmed cases and 4.3 million deaths worldwide as a result of COVID-19^2^. There has been significant effort from the global research community to develop effective vaccines for COVID-19; this has been enormously successful, with adenoviral vectored vaccines, protein vaccines and mRNA vaccines now in widespread use across the world. Whilst vaccines will enable protective immunity in most, there will be populations where vaccine-based immunity may fail, and these individuals will be vulnerable to SARS-CoV-2 infection in the future. It is furthermore unclear what changes will appear in the virus in contemporary SARS-CoV-2 viral variants (highlighted by the recent emergence of the delta variant B.1.617.2)^3^, and how those variants will navigate both convalescent and vaccine immune responses. Given that this is the third coronavirus that has crossed via zoonoses, antiviral development against SARS-CoV-2, and future coronaviruses with pandemic potential, are desperately needed in addition to prophylactic vaccines.

While there have been significant efforts toward the discovery of effective antivirals for SARS-CoV-2, the vast majority of molecules that have completed (or are currently being assessed in) clinical trials were originally developed for other infectious and inflammatory disease indications and are being repurposed for COVID-19. For example, remdesivir, originally trialled for Ebola, is currently the only antiviral drug to be approved by the U.S. Food and Drug Administration (FDA) for the treatment of COVID-19. While the molecule has been shown to possess some activity during early infection, it has shown limited to no efficacy in a number of trials^4,5^, as well as in infections in patients hospitalized with COVID-19^6^. Other repurposed antiviral drugs that have entered trials include the HIV combination therapy lopinavir-ritonavir^7,8^, type I interferon treatments^9,10^, and the antimalarial hydroxychloroquine^11–13^; however, these have not demonstrated improvement in disease progression over standard care. In fact, it has recently been suggested that many repurposing efforts may be compromised by experimental artefacts reflecting the physicochemical properties of certain drugs rather than specific target-based activities^14^. To date, the most effective therapeutic intervention for improving COVID-19 patient outcomes in a hospital setting is the use of the corticosteroid dexamethasone, which reduces inflammation-mediated lung injury associated with SARS-CoV-2 infection in patients with elevated levels of C-reactive protein^15 16^. Based on the above, there is an urgent need for the discovery of effective antivirals for COVID-19, ideally with mechanisms of action that specifically target proteins critical in the SARS-CoV-2 lifecycle.

Infection of human cells by SARS-CoV-2 is initiated by interaction between the receptor binding domain of the trimeric viral spike protein (S) with the host cell-surface receptor angiotensin converting enzyme 2 (ACE2) (Figure 1). Following receptor binding of the virus, the spike protein is activated by cleavage between the S1 and S2 domains leading to host cell entry via two distinct pathways: 1) an endocytic pathway through endosomal–lysosomal compartments with spike cleavage facilitated by lysosomal cathepsins, or 2) a cell surface pathway following activation by a serine protease such as TMPRSS2 ^17–19^. Following proteolysis, the N-terminus of the cleaved S2 domain is embedded into the cell membrane and leads to fusion of the membranes of the virus and the host cell, followed by transfer of viral RNA into the cytoplasm ^20^.

**Fig. 1.**
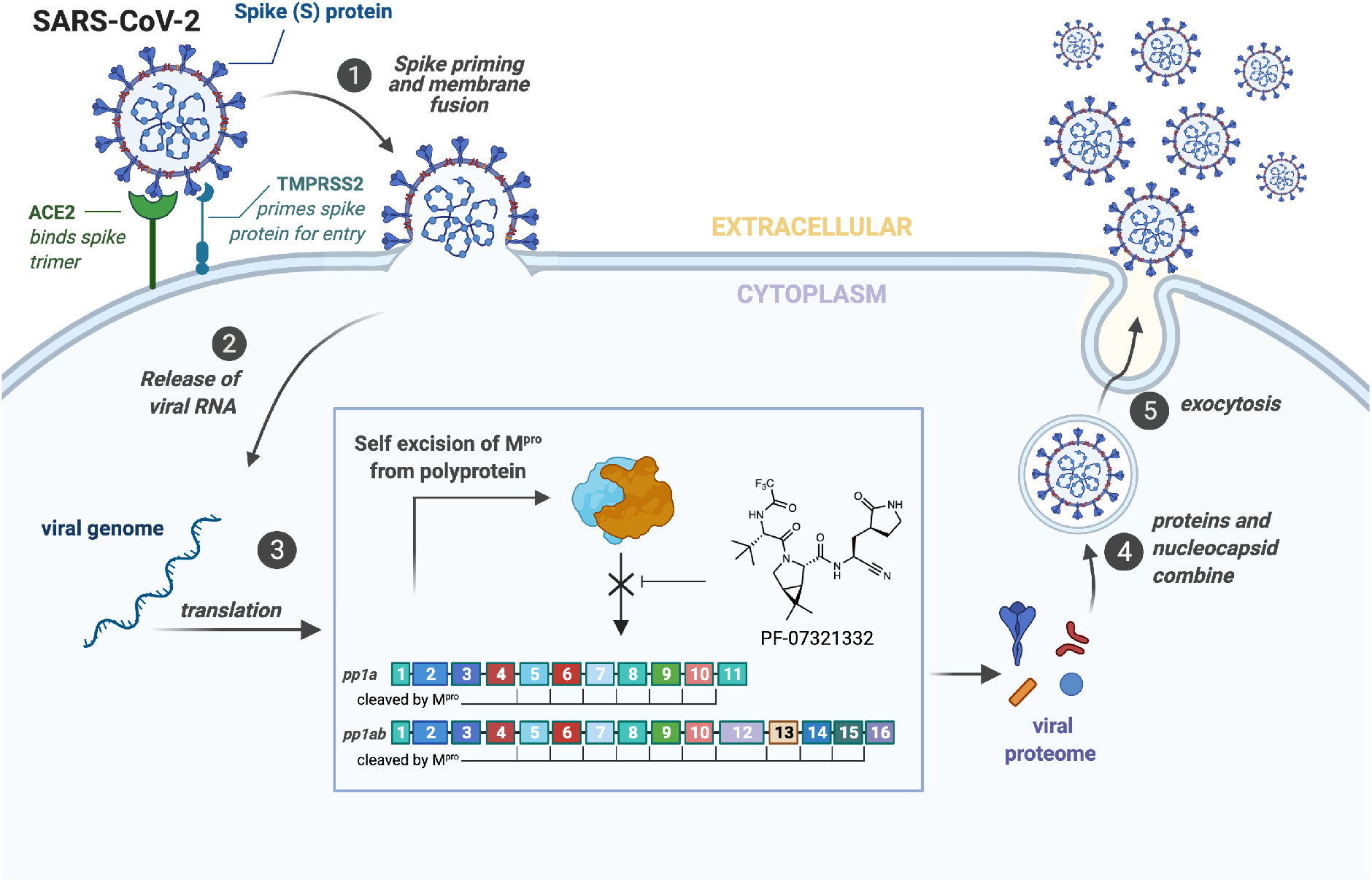
**Mechanism of SARS-CoV-2 entry into host cells and replication** (cell-surface entry pathway mediated by TMPRSS2 shown).The proteolytic activity of SARS-CoV-2 M^pro^ on the two polyproteins pp1a and pp1ab is shown in the box, including the structure of the covalent peptidomimetic M^pro^ inhibitor PF-07321332 developed by Pfizer.

Viral gene expression within the host cell results in the translation of two overlapping polyproteins, pp1a and pp1ab. Embedded within these polyproteins are sixteen non-structural proteins critical for viral replication, the majority of which form the viral replication and transcription complex (RTC) including the viral RNA-dependent RNA polymerase (RdRp, nsp12) and helicase (nsp13)^20^. These proteins become functional only after proteolytic release by two viral proteases. The first of these is a domain of nsp3 called the papain-like protease (PL^pro^) which cleaves the pp1a and pp1ab at three sites, releasing nsp1, nsp2 and nsp3^20,21^. The second is the SARS-CoV-2 main protease (M^pro^), also called nsp5 or the chymotrypsin-like protease (3CL^pro^), which cleaves pp1a and pp1ab at a minimum of 11 distinct cleavage sites to release nsp4-16 (Figure 1)^20^. Interestingly, M^pro^ has also been found to aid in immune evasion by inhibiting type I IFN production, contributing to the impaired type I IFN response that has become a hallmark of severe SARS-CoV-2 infection, with persistent viral load and poor patient outcomes^22–25^. M^pro^ forms a catalytically active homodimer which cleaves with high specificity at Leu-Gln↓Xaa (where ↓ represents the cleavage site and Xaa can be Ser, Ala or Asn)^26–28^. Such sequence specificity has not been observed for any human proteases and therefore peptide or peptidomimetic based inhibitors are predicted to inhibit SARS-CoV-2 M^pro^ with high selectivity and with minimal off-target effects in humans^20,29^.

The key role of M^pro^ for the replication and viability of SARS-CoV-2 has naturally led to the search for novel inhibitors of the protease. Perhaps the most promising of these are the peptidomimetic compounds developed by Pfizer, inspired by PF-00835231 (IC_50_ = 4-8 nM against SARS-CoV-2 M^pro^) that was originally developed against SARS-CoV-1 M^pro^, which possesses high homology to the SARS-CoV-2 protease^30–32^. Specifically, a phosphate prodrug of this inhibitor (PF07304814) has recently completed a phase 1b trial (clinical trials identifier NCT04535167). A second-generation orally available peptidomimetic M^pro^ inhibitor developed by Pfizer (PF-07321332, Figure 1) has also recently entered phase 2 clinical trials for treatment of COVID-19 (clinical trials identifier NCT04960202)^33^. Both molecules possess a γ-lactam mimic of the glutamine (Gln) residue found at the P1 position in physiological cleavage sites and also inhibit SARS-CoV-2 M^pro^ via a covalent mechanism through electrophilic warheads embedded within the inhibitors ^30^. Specifically, PF-00835231 possesses a hydroxymethyl ketone moiety, while PF-07321332 contains a nitrile warhead, both of which react with the catalytic cysteine (Cys145) to inactivate the protease. In addition to these clinical candidates, a number of other peptidomimetic inhibitors of SARS-CoV-2 M^pro^ are currently under preclinical investigation^34–37^, including repurposed drugs such as boceprevir, a serine protease inhibitor approved in 2011 for the treatment of hepatitis C^35,38,39^, and the feline anticoronaviral drug GC376 ^35,38^.

Macrocyclic peptides are attractive chemotypes for medicinal chemistry efforts due to their ability to bind targets with high affinity and selectivity, whilst exhibiting greater proteolytic stability and membrane permeability than their linear counterparts ^40–42^. In this work we describe several potent cyclic peptide inhibitors of the SARS-CoV-2 M^pro^, identified through the use of the <u>Ra</u>ndom nonstandard <u>P</u>eptide <u>I</u>ntegrated <u>D</u>iscovery (RaPID) technology, which couples mRNA display with flexizyme-mediated genetic code reprogramming^42–44^. Importantly, we also report a crystal structure of the SARS-CoV-2 M^pro^ dimer bound to our most potent cyclic peptide inhibitor that highlights the residues important for binding both at the catalytic site and across the dimer interface. Finally, we demonstrate that three of the cyclic peptides identified exhibited antiviral activity against SARS-CoV-2 *in vitro*, with an additional peptide gaining antiviral activity upon conjugation to a cell penetrating peptide.

## Results

### Selection against chemically cross-linked SARS-CoV-2 M^pro^ homodimer

In order to identify cyclic peptide inhibitors of SARS-CoV-2 M^pro^, we sought to utilize RaPID, which allows the screening of >10^12^ cyclic peptides for affinity against a protein target of interest immobilized on magnetic beads. However, functional SARS-CoV-2 M^pro^ is a homodimer with relatively weak affinity between the monomers (K_D_ = 2.5 μM), and we hypothesized that the protein may exist in a predominantly monomeric (i.e. inactive) form when immobilized on magnetic beads27. Consistent with this, we found that the catalytic activity (measured by fluorescent substrate cleavage) of C-terminally His6-tagged M^pro^ was significantly diminished (ca. 30%) when immobilized on His Pull-down Dynabeads™ compared to that of the wild-type protein in solution (**Supplementary Fig. 1a**); this indicated that a significant proportion of the immobilized M^pro^ was unable to form an active homodimer. We therefore investigated the use of a chemical cross-linking strategy to covalently lock SARS-CoV-2 M^pro^ in the homodimeric state to ensure that RaPID resulted in selection of ligands to the catalytically active form of the protease. The presence of proximal lysine residues near the dimer interface in the crystal structure of SARS-CoV-2 M^pro^ (PDB: 6Y2E) prompted us to investigate a number of amine-reactive bifunctional cross-linkers of differing lengths for this purpose27. Efficient cross-linking was obtained by disuccinimidyl glutarate (DSG) (**Figure 2a and 2b**), with subsequent mapping by LC-MS/MS revealing intermolecular cross-links between lysine residues on the different monomer chains (K97-K97*, K97-K90*). We also observed cross-links between K12 and K97, which are positioned in close proximity on different monomers of the dimer, as well as within the same monomer unit; however, in this case we could not differentiate between intermolecular or intramolecular cross-links by mass spectrometry (**Supplementary Fig. 1b**). Importantly, cross-linked M^pro^ exhibited catalytic activity comparable to that of wild-type M^pro^ (in solution), following immobilization on magnetic beads (**Supplementary Fig. 1a**). Based on these data, we moved on to RaPID selections against the cross-linked SARS-CoV-2 M^pro^ with a view to discovering novel cyclic peptide inhibitors.

**Fig. 2.**
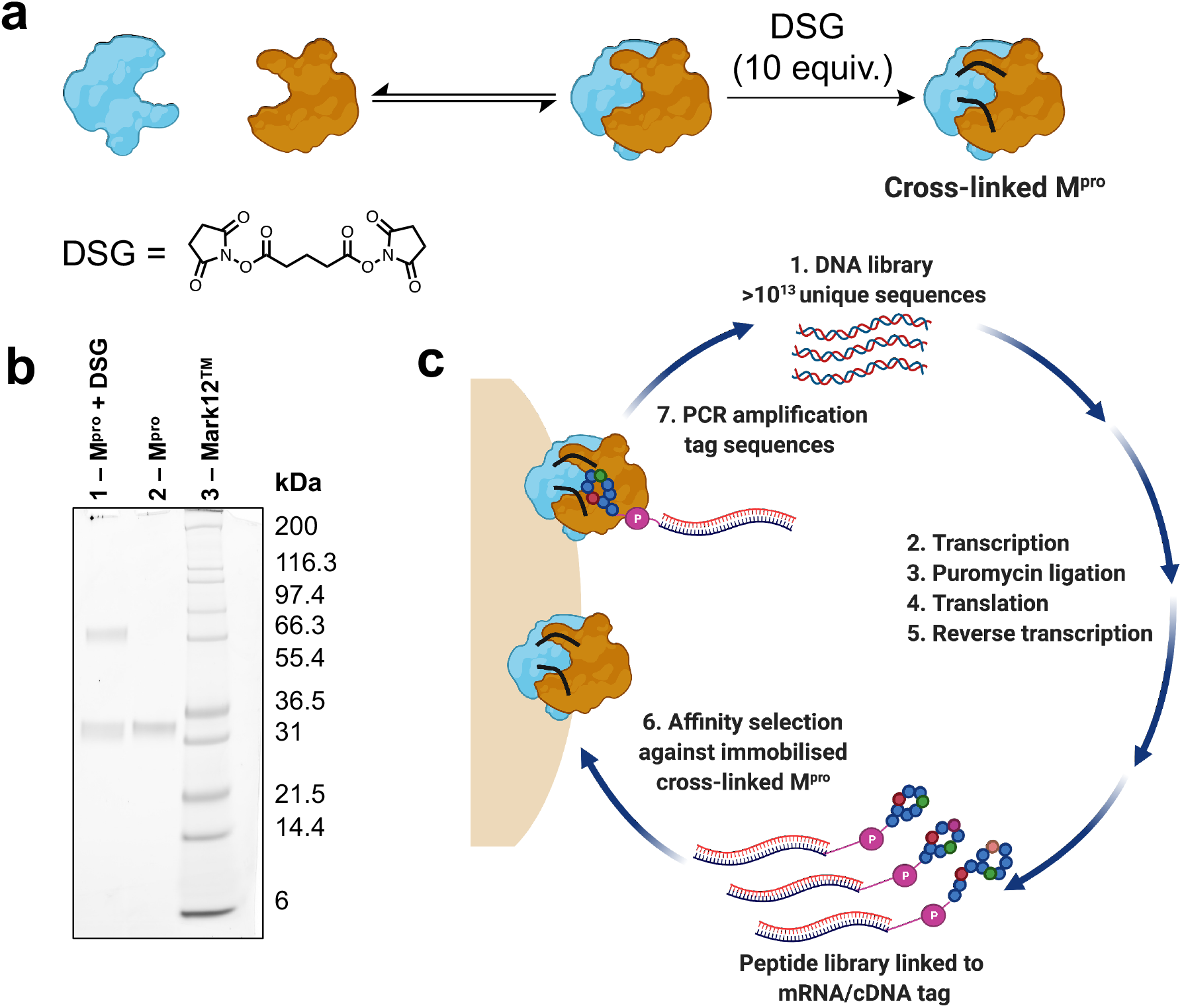
**a** Scheme for cross-linking of SARS-CoV-2 M^pro^ with disuccinimidyl glutarate (DSG). **b** SDS-PAGE gel of cross-linking reaction of M^pro^. Lane 1: Reaction of SARS-CoV-2 M^pro^ (25 μM) with DSG (10 equivalents relative to M^pro^ monomer) for 1 h at 37°C in aqueous buffer (20 mM HEPES, 100 mM NaCl, pH 7.6). Lane 2: SARS-CoV-2 M^pro^. Lane 3: Mark12™ ladder. **c** Workflow for the discovery of macrocyclic peptide ligands of cross-linked SARS-CoV-2 M^pro^ using RaPID mRNA display.

To select for peptide binders of SARS-CoV-2 M^pro^ we performed parallel display using the RaPID mRNA display technology, employing peptide libraries initiated with either *N*-chloroacetyl-L-tyrosine or *N*-chloroacetyl-D-tyrosine to induce thioether macrocyclization through reaction with the side chain of a downstream cysteine residue. Tyr was chosen as the initiating amino acid due to its high translation efficiency in place of *N*-formyl-Met^45,46^, while the use of D-Tyr in addition to L-Tyr expanded the accessible chemical space of the library. For each selection, a semi-randomized DNA library of >10^13^ unique sequences were transcribed into mRNA followed by ligation to a puromycin linker (**Figure 2c**). Translation of the mRNA-puromycin hybrids *in vitro*, followed by reverse transcription, yielded a library of peptide-mRNA:cDNA conjugates that were counter-selected against His Pull-down Dynabeads™ (to remove bead-binding peptides) before an affinity selection step, in which the libraries were panned against bead-immobilized cross-linked SARS-CoV-2 M^pro^. PCR of the bead-bound fraction yielded an enriched DNA library that was used as the starting point for the subsequent rounds of selection (**Figure 2c**). Following nine iterative rounds of this process, next-generation sequencing of the final DNA library identified several peptide sequences predicted to bind to SARS-CoV-2 M^pro^ based on high relative abundance (**Supplementary Fig. 2**).

### Synthesis and *in vitro* evaluation of cyclic peptide inhibitors of SARS-CoV-2 M^pro^

We selected eight peptides from the enriched libraries for synthesis (five L-Tyr initiated and three D-Tyr initiated) based on their abundance in the final DNA library and diversity in sequence (**Figure 3a** and **3b**). Peptides **1**–**8** were synthesized *via* Fmoc-strategy solid-phase peptide synthesis on Rink amide resin with the N-terminal L- or D-Tyr residue derivatized with chloroacetic acid to facilitate thioether cyclization with the thiol of a downstream cysteine residue. Deprotection and cleavage from resin followed by cyclization provided the target thioether-linked cyclic peptides **1**–**8** following purification by reverse-phase HPLC. The synthetic cyclic peptides were next evaluated for inhibitory activity against SARS-CoV-2 M^pro^ using a previously reported fluorescence resonance energy transfer (FRET)-based assay^27,47^. Gratifyingly, we observed potent inhibition of proteolytic activity for six of the eight peptides in the series, with IC_**50**_ values for peptides **1**–**6** ranging from 0.070 – 12.7 μM (**Figure 3c**). Interestingly, despite being enriched in the selection, lariat peptide **7** and head to tail cyclic peptide **8** did not show appreciable inhibition of the protease (IC_50_ > 50 μM). The most potent SARS-CoV-2 M^pro^ inhibitor in the series was cyclic peptide **1**, which exhibited an IC_50_ of 70 ± 18 nM and a *K*_i_ of 14 ± 3 nM against the protease. Given the superior inhibitory potency of this molecule over the other cyclic peptides that emerged from the RaPID screen (>15-fold), this molecule was selected for further structure-activity investigations.

**Fig. 3.**
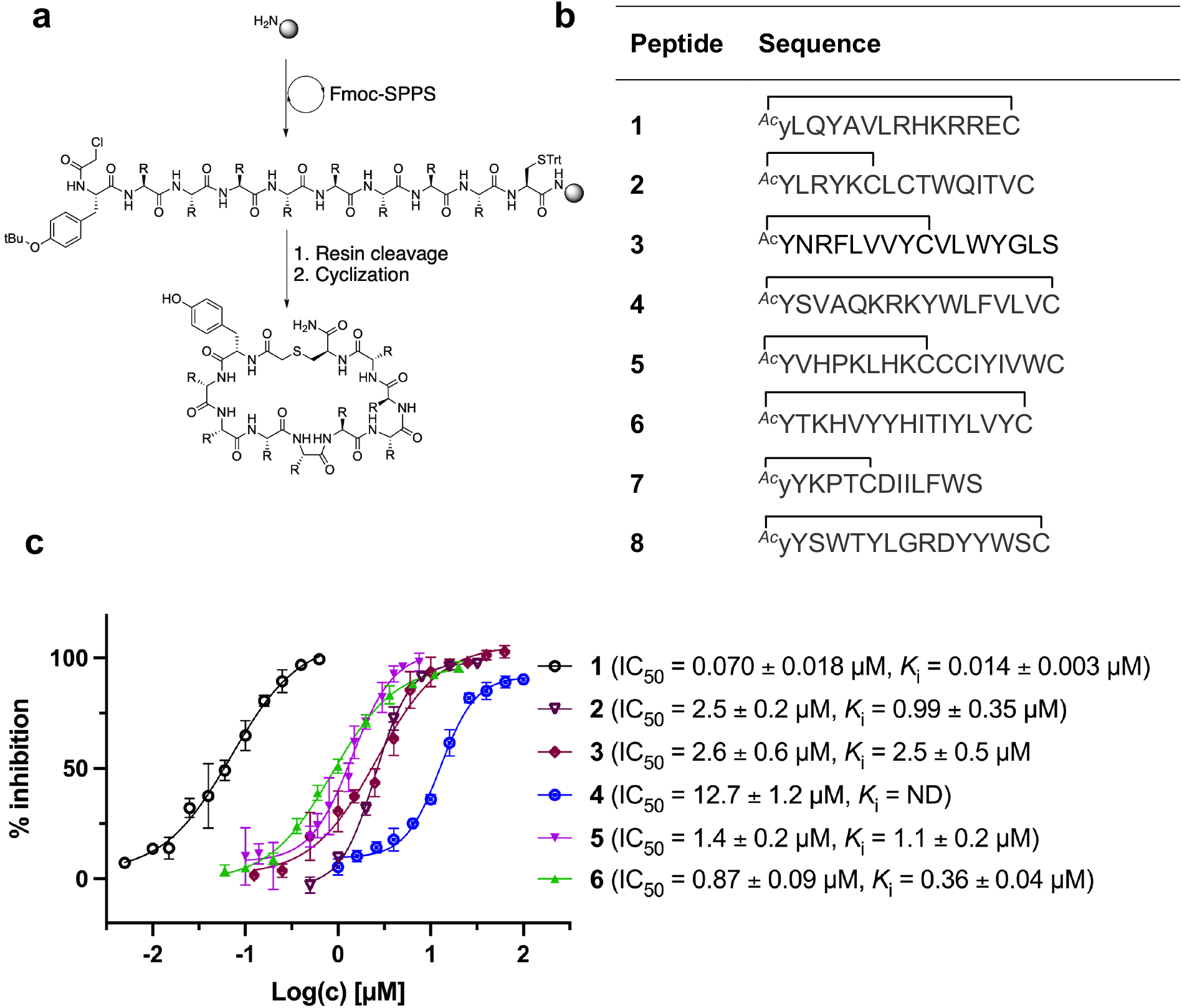
**a** Synthesis of cyclic peptides **1**–**8** *via* Fmoc-SPPS (see Supplementary methods for full synthetic details). **b** Sequences of peptides **1**-**8** in one letter amino acid code, thioether cyclization is represented as a black line. **c** *In vitro* inhibition data of SARS-CoV-2 M^pro^ for peptides **1**–**6** with associated IC_50_ and *K*_i_ values ±SEM. NB: cyclic peptides **7** and **8** showed no inhibition of SARS-CoV-2 M^pro^ at 50 μM, ND = not determined.

### Probing the binding interaction of lead cyclic peptide 1

We next performed experiments to probe the key interactions of lead cyclic peptide inhibitor **1** with SARS-CoV-2 M^pro^. Given the critical role dimerization plays in M^pro^ catalytic activity, we first investigated the effect of **1** on the monomer-dimer equilibrium of M^pro^ by multi-angle laser light scattering with size exclusion chromatography (SEC-MALLS). Interestingly, the addition of two molar equivalents of peptide **1** to a solution of wild-type M^pro^ (present in an approximately 1:1 ratio of monomer:dimer) resulted in a shift in the dimer-monomer equilibrium to afford a solution of exclusively homodimeric protease (**Figure 4a**). This data indicates that cyclic peptide **1** stabilizes the homodimeric form of SARS-CoV-2 M^pro^ and suggests that peptide **1** may bind to the active site present in the dimeric, but not monomeric forms of the protein. To corroborate this data, we used 3D NMR spectroscopy (specifically TROSY-HNCO spectra) to analyze SARS-CoV-2 M^pro^ after titration with peptide **1** (**Figure 4b**). This revealed shifts of backbone NMR resonances of residues near the active site, consistent with binding of peptide **1** at this location. However, shifts were also observed for residues located far from the active site (**Supplementary Fig. 3**), suggesting that the protein responds to the binding of **1** with global allosteric changes. In contrast, titration of the obligate monomeric mutant R298A of SARS-CoV-2 M^pro 48^ with peptide **1** led to no shifts in NMR resonances (**Supplementary Fig. 3**, see **Supplementary Tables 1** and **2** and **Supplementary Fig. 4** for full assignment of monomeric M^pro^ mutant). Notably, R298, which is mutated in the obligate monomer, is buried within the dimer interface in the structure of the wild-type protease and therefore the lack of binding of **1** to the monomeric protein is unlikely to be owing to a loss of interactions with this amino acid residue. Taken together, these SEC-MALLS and NMR data therefore suggest that peptide **1** selectively binds to and inhibits the active homodimer of SARS-CoV-2 M^pro^ but does not bind to the inactive monomer.

**Fig. 4.**
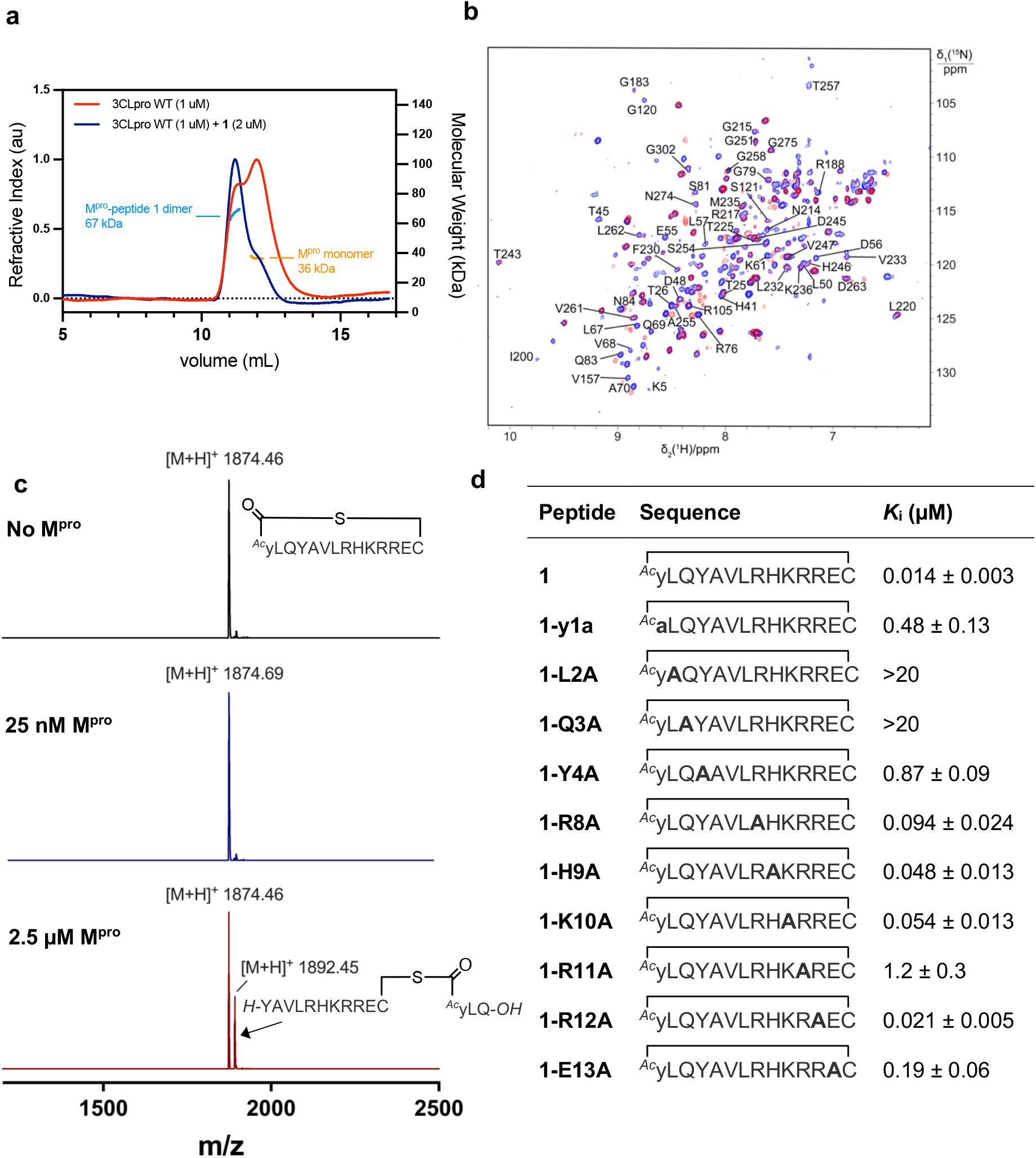
**a SEC-MALLS of SARS-CoV-2 M^pro^ with and without peptide 1.** SARS-CoV-2 M^pro^ (red line) exists in equilibrium between monomeric and homodimeric forms giving rise to two peaks (in ca. 1:1.25 ratio of monomer to dimer) in the size-exclusion chromatogram at a concentration of 1 μM in aqueous buffer (50 mM Tris-HCl pH 8.0, 150 mM NaCl). MALLS analysis indicates an approximate molecular weight of 36 kDa for the monomer (calculated molecular weight = 33.8 kDa). After addition of two molar equivalents of **1**, M^pro^ converges predominantly to a homodimer (blue line) with a MALLS reading of 66 kDa (calculated MW of SARS-CoV-2 M^pro^ dimer = 67.6 kDa, MW of peptide **1**= 1874 Da) indicative of formation of the SARS-CoV-2 M^pro^ homodimer upon binding **1**. **b Cyclic peptide inhibitor 1 binds to the dimeric form of SARS-CoV-2 M^pro^**. Overlay of projections onto the ^15^N-^1^H plane of 3D TROSY-HNCO spectra of 0.3 mM solutions of ^15^N/^13^C/^2^H-labelled wild-type M^pro^. Blue and red contour lines show the spectra recorded in the absence and presence of equimolar inhibitor **1**, respectively. Assignments are shown for peaks that shift or disappear in response to the inhibitor. **c Monitoring of the cleavage of cyclic peptide inhibitor 1 with SARS-CoV-2 M^pro^**. MALDI-TOF mass spectrum of cyclic peptide **1** (top spectrum). Negligible cleavage of **1** was observed following incubation with SARS-CoV-2 M^pro^ under standard assay conditions (25 nM SARS-CoV-2 M^pro^, 5 μM **1**, 20 mM Tris-HCl pH 7.6, 100 mM NaCl, 1 mM DTT, 1 mM EDTA) for 1 h at 37 °C (middle spectrum). Slow cleavage of **1** was observed (ca. ~30% after 1 h) in the presence of a high concentration of SARS-CoV-2 M^pro^ to 2.5 μM (bottom spectrum). **d Inhibitory activity of alanine mutants of lead cyclic peptide 1**. Sequences and associated inhibitory constants for peptide **1** analogues, whereby all polar residues within the randomized region of **1** were each systematically mutated to alanine and their inhibitory activity against SARS-CoV-2 M^pro^ assessed (Cys14 was not mutated as this residue is required for cyclization, Ala substitutions are shown in bold).

Cyclic peptide **1** includes the cleavage recognition motif of M^pro^ found in the natural viral protein substrates, namely a Gln in the P_1_ position and a Leu in the P_2_ position^27^. It was therefore hypothesized that this Leu-Gln motif embedded within **1** mimics the natural substrate and binds to the catalytic site of the protease. Importantly, this proposal is supported by the purely competitive inhibition mode observed for **1** (**Supplementary Fig. 5**). However, this also raises the possibility that the protease may be able to cleave **1** next to the Gln-Leu recognition sequence. To test this, we incubated **1** with wild type SARS-CoV-2 M^pro^ and assessed cleavage by MALDI-TOF-MS (**Figure 4c**). Peptide **1** exhibited notable resistance to proteolysis by M^pro^ with negligible cleavage of peptide **1** observed under standard assay conditions (25 nM M^pro^) after 1 h. However, incubation of **1** with a high concentration of M^pro^ (2.5 μM) resulted in slow cleavage of peptide **1**, with 30% peptide cleavage observed after 1 h. Analysis of the cleavage reaction by LC-MS/MS confirmed that proteolysis had indeed occurred between Gln3 and Tyr4 (**Supplementary Fig. 6a and 6b**). We also synthesized an authentic standard of the resulting cleavage product, which was verified to be identical to M^pro^-cleaved **1** by LC-MS/MS (**Supplementary Fig. 6b**). Finally, we assessed whether the linear peptide product resulting from M^pro^ cleavage of **1** possessed inhibitory activity against SARS-CoV-2 M^pro^. Interestingly, the peptide exhibited an IC_50_ of 23.2 ± 5 μM, ~330-fold higher than the IC_50_ of 70 nM for **1** (**Supplementary Fig. 6c**). This two-orders of magnitude loss in activity upon linearization suggests that the conformation of cyclic peptide **1** is pre-organised for optimal interaction with the protease.

In order to assess the importance of each residue in **1** for M^pro^ inhibitory activity, we systematically replaced all residues in the peptide with alanine (except alkyl side chain-containing amino acids Ala5, Val6 and Leu7) and determined the inhibitory activity of the resulting mutants against M^pro^ (**Figure 4d, Supplementary Fig. 7**). Consistent with the known recognition sequence for SARS-CoV-2 M^pro^ and supported by the mass spectrometry results described above, mutation of either Leu2 or Gln3 to Ala (that would be predicted to bind in the S2 and S1 recognition sites, respectively) led to more than two orders of magnitude reduction in inhibitory activity. Remarkably, mutation of Tyr4 (which would be predicted at P_1_’) also led to a significant loss in inhibitor activity (IC_50_ = 1.9 μM); while this is consistent with the established importance of aromaticity in P_1_’, small residues such as alanine are often found in substrates at this position, and the dramatic reduction in inhibitory potency was therefore unexpected ^27,28^. Interestingly, mutation of Arg11, which is distal from the most prominent recognition residues of the cyclic peptide, also led to a significant reduction in inhibitory activity (IC_50_ = 3.4 μM) suggesting that this residue makes important interactions with the protease and/or serves a crucial role in the adoption of the active conformation of the cyclic peptide. In contrast, mutation of Arg8, His9, Lys10, Arg12 or Glu13 resulted in equipotent activity or only a modest reduction in inhibitory activity (IC_50_ values = 90–390 nM).

### Co-crystallization of peptide 1 and a selenoether analogue with SARS-CoV-2 M^pro^

In order to further interrogate the binding mode of inhibitor **1**, we used X-ray diffraction to solve the co-crystal structure of the SARS-CoV-2 M^pro^-**1** complex to 3.4 Å. The limited resolution of the structure hindered the interpretation of the electron density within the active site. We therefore also solved the structure of a synthetic selenoether analogue of **1** (**Se-1**) in complex with SARS-CoV-2 M^pro^, in which the cysteine residue involved in thioether peptide macrocyclization was replaced with selenocysteine^49^; we rationalized that the greater electron density of Se in this analogue would aid its placement in the active site. Importantly, **Se-1** displayed identical inhibitory activity as **1** (*K*_i_ of **Se-1** = 17 nM vs 14 nM for 1), suggesting its interactions with the protein are also very similar to parent inhibitor **1** (**Supplementary Fig. 8**). A crystal of this SARS-CoV-2 M^pro^-**Se-1** complex diffracted to 2.19 Å resolution in the same space group and with similar cell dimensions as the SARS-CoV-2 M^pro^-**1** complex, thus providing a detailed view of the interactions between the inhibitor and the protease (**Supplementary Table 3**). The crystal dimensions and packing are unique and not observed in any of the hundreds of deposited SARS-CoV-2 M^pro^ structures, with a notably long *b* axis. There were four protein subunits within the asymmetric unit comprising two physiological dimers (**Figure 5a**).

**Fig. 5.**
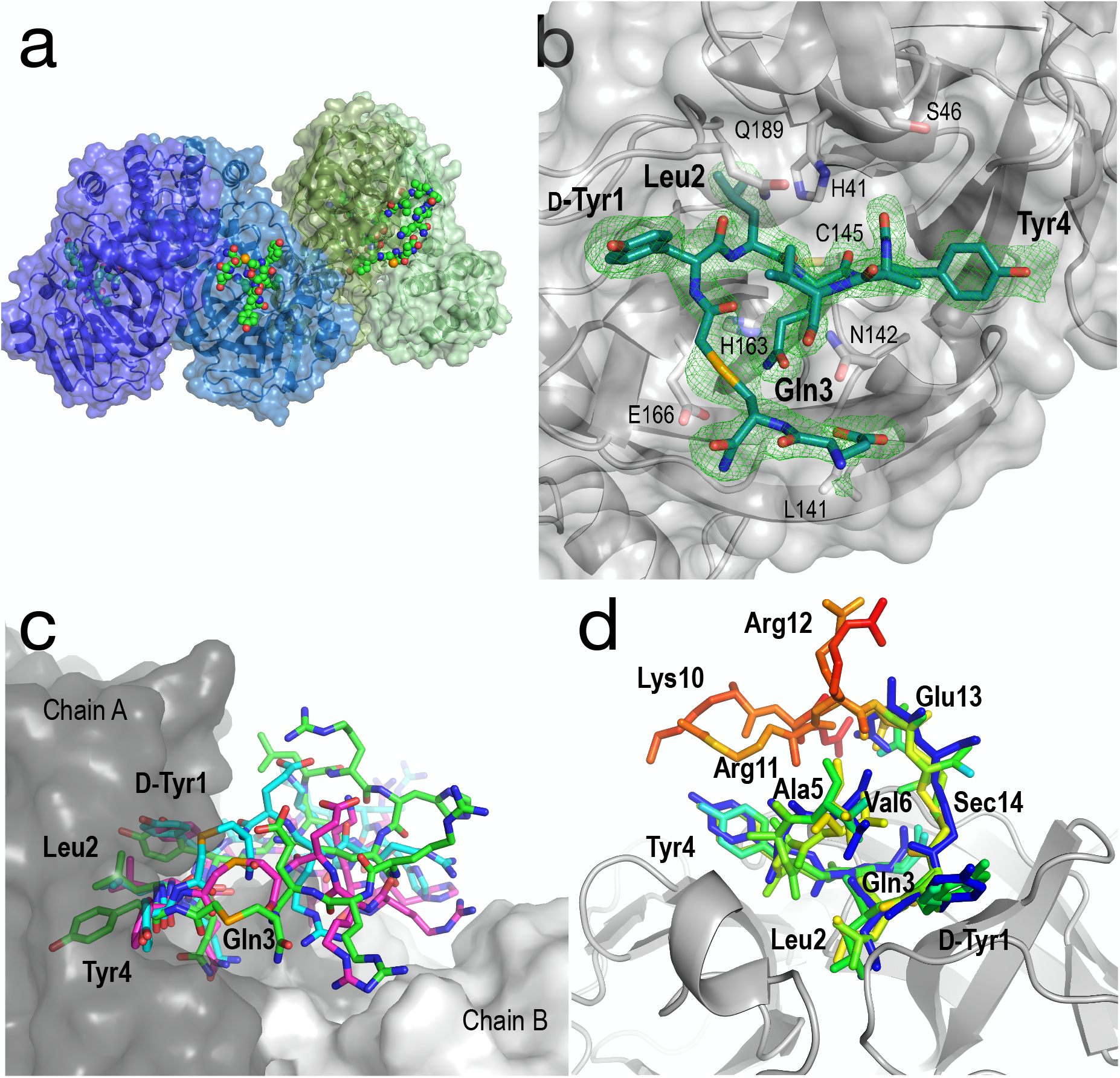
**Structural analysis of the SARS-CoV-2 M^pro^-Se-1 complex (PDB ID: 7RNW). a** The asymmetric unit of the SARS-CoV-2 M^pro^-**Se-1** crystal structure, containing two physiological dimers shown as blue and green. The **Se-1** peptide is shown in a sphere representation. **b** Zoom of the active site with electron density (Polder omit map, contoured to 3.5σ) showing bound peptide (residues 7–12 disordered). Key active site residues of SARS-CoV-2 M^pro^ are shown as sticks, and key positions of the **Se-1** peptide (residues 1-4) are labelled in bold text. **c** Representative conformations (300 ns) from each of the triplicate simulations suggest that Tyr1, Leu2, Gln3 and Tyr4 are stably bound in the active site of SARS-CoV-2 M^pro^, while the remainder of the peptide is more mobile and makes transient interactions across the dimer interface. **d Se-1** peptide bound to chains A, C and D coloured by B-factor, showing the residues 1-4 are stable and bound within the S3, S2, S1 and S-1 subsites, respectively, whereas the polar half of the peptide (Arg8-Glu13) is either too disordered to accurately model, or is modelled with high B-factors.

Although the peptide is not fully resolved in the structure, 9-13 residues were observable in various chains with good electron density; the selenoether linkage and the first 5 residues (D-Tyr1, Leu2, Gln3, Tyr4, Ala5) were very stable, with the remainder of the peptide (7-12) appearing somewhat disordered (**Figure 5b**). The peptide displayed a consensus binding pose across three subunits of the crystal structure. The fourth subunit displayed an alternative binding pose in which Leu2 and Gln3 were present in identical positions, but the flanking D-Tyr1 and Tyr4 residues adopted alternative conformations (**Supplementary Fig. 9a and 9b**). Closer inspection revealed a non-Pro *cis-*peptide bond between Tyr4 and Ala3 in chains A, C, D, which results in a tight kink in the helix, while in chain B this bond remains in the *trans*-configuration (**Supplementary Fig. 9c and 9d**). The Gln3 of the peptide occupies the canonical S_1_ subsite of the protease (following the numbering by Lee *et al*.) ^50^ in every peptide:protein complex, facilitated by interactions with His163, Glu166, and Asn142 (**Figure 5b**). Likewise, Leu2 is always bound in the S_2_ subsite comprised of His41, His164 and Gln189. D-Tyr1 is in the solvent-exposed S_3_ site bordered by Gln189, Ala191 and Pro16, while the peptide twists at the selenoether/D-Tyr1 linkage, turning away from the canonical S_4_ site to be positioned adjacent to Glu166. Tyr4 is therefore positioned in the S_1_’ (P_1_’) position in ¾ chains, fully consistent with the slow proteolysis between Gln3 and Tyr4 observed by mass spectrometry (**Figure 4c**). In chain B, the peptide backbone and Tyr4 have swapped positions (**Supplementary Fig. 9a and 9b**).

To obtain a better understanding of the interaction of the full peptide with the protein in the absence of crystal contacts, we modelled the missing residues as accurately as possible with the available density and used this structure as the starting point for triplicate 333 ns molecular dynamics (MD) simulations ^51^ (~1 μs total simulation time; **Figure 5c**, **Supplementary Fig. 10**). The results were consistent with the crystal structure, i.e. residues 1-4 were all relatively stable within their respective binding pockets within the substrate cleft of SARS-CoV-2 M^pro^, with the mobility of the peptide increasing on either side of these residues (**Figure 5d**). Notably, the simulations showed regular transient interactions between the peptide and the second chain of the physiological dimer.

The structure and MD simulations also provide a molecular explanation for the inhibitory activity of the alanine mutants. Specifically, the structure shows that the central interactions are formed by Leu2 and Gln3, consistent with the large reductions in activity when these positions are mutated to Ala, while D-Tyr1 and Tyr4 also form significant interactions with the protease on either side of these residues. Other positions (8, 9, 10, 12 and 13) that were observed to have little influence on inhibitory activity are either disordered or solvent-exposed. MD simulations suggest that Arg11, the mutation of which to Ala had a significant effect on inhibition (**Figure 4d**), makes contacts across the dimer interface (**Supplementary Fig. 9e**), interacting with the side chain and main chain carbonyl of Q256*. This is consistent with electron density showing Arg11 within hydrogen bonding distance of the neighbouring chain B in one dimer (**Supplementary Fig. 9f)**. These observations are consistent with the SEC-MALLS and NMR data that shows the peptide binds the dimeric form of SARS-CoV-2 M^pro^ exclusively for its mode of inhibition (**Figure 4a and 4b**).

### Antiviral activity of cyclic peptide SARS-CoV-2 M^pro^ inhibitors

Given the promising inhibitory activity of cyclic peptides **1–6** against SARS-CoV-2 M^pro^, we next assessed the antiviral activity of the four most potent peptides against SARS-CoV-2 *in vitro*. Specifically, we used ACE-2 and TMPRSS2 overexpressing HEK293T cells infected with SARS-CoV-2 to assess each peptide. The latter cell line is hyper-permissive to SARS-CoV-2 infection with extensive viral syncytia forming after 18 hours post viral infection. Dose-dependent sigmoidal inhibition curves can be generated through enumeration of remaining single cell nuclei using a live nuclear dye and a standard high content fluorescent microscope. In this setting, we observed a dose-dependent protection for three of the four peptides, with EC_50_ values ranging from 11.8 – 33.1 μM. However, peptide **1**, the most active compound in the enzyme inhibition assay, was inactive up to a concentration of 50 μM, most likely due to low cell permeability owing to the large number of polar and charged residues in this molecule. To facilitate cell entry of **1**, we therefore conjugated its C-terminus to penetratin, a 16 amino acid cell-penetrating peptide (CPP) derived from the *Drosophila* Antennapedia homeodomain ^52^ (to afford **pen-1**). It should be noted that during the selection process the C-terminus of the peptides is conjugated to a large mRNA:cDNA tag, and therefore addition of a CPP to the C-terminus of the peptide (to afford **pen-1**) was deemed unlikely to affect binding and inhibition of M^pro^. Pleasingly, **pen-1** had equivalent *in vitro* inhibitory activity against SARS-CoV-2 M^pro^ to peptide **1** (*K*_i_ = 9 nM for **pen-1** vs 14 nM for **1**, **Supplementary Fig. 11**) indicating that addition of a CPP to the C-terminus of the peptide did not affect binding and inhibition of the protease. We also prepared a penetratin conjugate of cyclic peptide **6**, the second most active peptide in the enzyme inhibition assay (*K*_i_ = 360 nM). While parent cyclic peptide **6** showed antiviral activity in the cell-based assay (EC_50_ of 33.1 μM), we anticipated that the conjugation to penetratin might further improve potency. We also assessed the cytotoxicity of **1**, **pen-1**, **6** and **pen-6** on HEK293-ACE2-TMPRSS2 cells, with none of the compounds affecting cell viability at a concentration of 50 μM (**Supplementary Fig. 12**). Pleasingly, we found that **pen-1** and **pen-6** both exhibited significantly improved antiviral activity, with EC_50_ values of 15.2 μM and 6.6 μM, respectively (**Figure 6**). As expected, the marked improvement in antiviral activity of peptide **1** by conjugation to the penetratin CPP correlated with enhanced cellular uptake, whereby LC-MS/MS analysis showed a 5.5-fold increase in levels of **pen-1** compared to **1** in cell lysates (**Supplementary Fig. 13**).

**Fig. 6.**
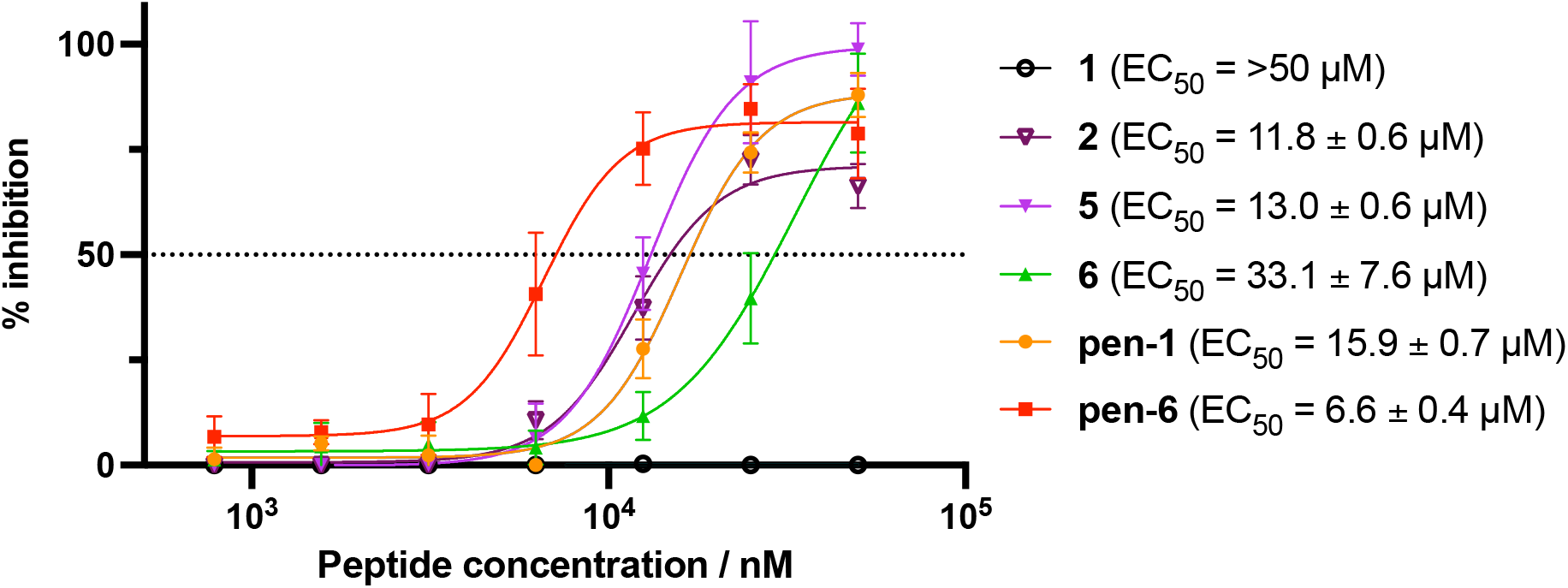
**Antiviral activity of peptides 1, 2, 5, 6 and penetratin conjugates of 1 and 6 (pen-1 and pen-6).** HEK293-ACE2-TMPRSS2 cells were incubated with varying concentrations of cyclic peptide M^pro^ inhibitors and infected with SARS-CoV-2. Inhibition curves and 50% effective concentrations (EC_50_) were determined by non-linear regression analysis using GraphPad Prism. Data are the means ± SD of experiments performed in quadruplicate.

## Discussion

In summary, we discovered macrocyclic peptide inhibitors of SARS-CoV-2 M^pro^ using RaPID mRNA display technology. A protein cross-linking strategy was employed to generate a covalently locked catalytically active dimer, facilitating display selection against solid-supported SARS-CoV-2 M^pro^. This enabled the discovery of potent inhibitors of M^pro^, including the 14-residue cyclic peptide 1with a *K*_i_ of 14 nM that represents one of the most potent inhibitors reported against the protease to date. SEC-MALLS and NMR spectroscopic studies revealed that this molecule inhibits SARS-CoV-2 M^pro^ by exclusively binding to the catalytic protease dimer, highlighting the importance of controlling the nature of the immobilized protein construct for display selections; in this case by covalently cross-linking the M^pro^ homodimer. Co-crystal structures of M^pro^ and **1** and the selenocysteine derivative **Sec-1** solved by X-ray crystallography revealed the canonical interaction between a Gln residue and subsite S_1_, flanked by hydrophobic residues. This half of the peptide was relatively stable, while the hydrophilic face (8–12) was relatively mobile, interacting with solvent and across the dimer interface to facilitate an exclusive dimer binding mode for the inhibitor. The strain imposed by cyclization prevents the peptide from adopting a fully relaxed conformation and likely explains the very slow turnover, despite the inhibitor possessing canonical residues that bind to the key recognition subsites of the protease. Thus, cyclization of the peptide has essentially converted a peptide substrate into a highly potent SARS-CoV-2 M^pro^ inhibitor, highlighting a key benefit of the cyclic peptides that emerge from RaPID screens in medicinal chemistry studies. Several of the cyclic peptides also exhibited antiviral activity against SARS-CoV-2 *in vitro* and enhancing cellular uptake of cyclic peptides by conjugation to the CPP penetratin led to dramatic improvements in antiviral activity. This was especially pronounced for lead molecule **1**, which was ineffective without the covalent CPP tag but exhibited an EC_50_ of 15.9 μM following fusion to penetratin. Taken together, these compounds now serve as bona fide starting points for the rational design of peptide- or peptidomimetic-based antivirals for COVID-19 that target SARS-CoV-2 M^pro^. Moreover, the crosslinked protease selection strategy developed as part of this work is expected to inform medicinal chemistry campaigns targeting other dimeric coronaviral main proteases. Future work in our laboratories will involve lead optimization of the discovered peptides for improved activity and cell permeability both by rational design and high-throughput affinity maturation^53,54^ with a view to developing molecules with potent antiviral activity *in vivo*.

## Methods

### Plasmid construction

The gene of the SARS-CoV-2 main protease (M^pro^)^27^ was cloned in between the *Nde*I and *Xho*I sites of the T7 vector pET-47b (+). The construct contains the M^pro^ self-cleavage-site (SAVLQ↓SGFRK; arrow indicating the cleavage site) at the N-terminus. At the C-terminus, the construct contains a modified PreScission cleavage site (SGVTFQ↓GP) connected to a His_6_-tag. All plasmid constructions and mutagenesis were conducted with cloning and QuikChange protocols using mutant T4 DNA polymerase.

### Protein expression

Wild-type M^pro^ was expressed in BL21 DE3 cells transformed with the desired plasmid. Protein expression was conducted in a Labfors 5 bioreactor (INFORS HT, Switzerland). After induction, the culture was grown at 18 °C overnight for protein expression. Cells were harvested by centrifugation at 5,000 g for 15 minutes and lysed by passing twice through a Emulsiflex-C5 homogenizer (Avestin, Canada). The lysate was centrifuged at 13,000 *g* for 60 minutes and the filtered supernatant was loaded onto a 5 mL Ni-NTA column (GE Healthcare, USA) equilibrated with binding buffer (50 mM Tris-HCl pH 7.5, 300 mM NaCl, 5 % glycerol). The protein was eluted with elution buffer (binding buffer containing, in addition, 300 mM imidazole) and the fractions were analyzed by 12% SDS-PAGE. PreScission cleavage and TEV cleavage were conducted in binding buffer in the presence of 1 mM DTT with a protein-to-protease ratio of 100:1. Following cleavage of the His6-tag, the buffer was exchanged to 20 mM HEPES-KOH pH 7.0, 150 mM NaCl, 1 mM DTT, 1 mM EDTA. All samples were analyzed by mass spectrometry using an Orbitrap Fusion Tribrid mass spectrometer (Thermo Scientific, USA) coupled with an UltiMate S4 3000 UHPLC (Thermo Scientific, USA).

### Protein cross-linking

C-terminally His-tagged SARS-CoV-2 M^pro^ (25 μM) was incubated with disuccinimidyl glutarate (H6/D6, Creative Molecules Inc.) (250 μM) in aqueous buffer (20 mM HEPES pH 7.6, 100 mM NaCl) at 37 °C for 1 h before the reaction was quenched by dilution of the protein to 10 μM (monomer concentration) by addition of aqueous buffer containing 20 mM Tris-HCl pH 7.6, 100 mM NaCl. Cross-linking efficiency was analyzed by 12% SDS-PAGE and stained with Sypro™ Ruby.

### Proteomics to determine crosslink sites on SARS-CoV-2 M^pro^

Crosslinked proteins were digested with trypsin and peptides desalted as described previously^55^. Peptide mixtures were analyzed by LC-MS/MS using data dependent acquisition with HCD fragmentation on a Thermo Orbitrap Eclipse mass spectrometer. Crosslinked peptides were identified with a 2% false-discovery rate using the Byonic search engine (Protein Metrics) and a custom database containing the 3CL protein sequence and common proteomics contaminants (e.g. trypsin, albumin, keratins, etc.). DSG crosslinks and hydrolysis products were allowed for the 3CL protein only. Carbamidomethylation was specified as a fixed modification on C, while oxidation of methionine, deamidation of N/Q and pyro-Glu for N-terminal Q/E were variable modifications. Plots of crosslinked residues were generated using UCSF Chimera 1.15rc with the 3CL dimer structure (PDB: 6Y2E).

### RaPID mRNA display selection

Library preparation and display selection were performed as described previously with slight modifications ^44,56–58^. Briefly, an mRNA library comprising an AUG start codon, 4–15 NNS (N = A, C, G or T; S = C or G) codons, a TGC cysteine codon and a 3’ region encoding a linking sequence (Gly-Asn-Leu-Ile) with mRNA lengths pooled proportional to theoretical diversity was prepared as described previously. The mRNA library was then ligated to a puromycin-linked oligonucleotide using T4 ligase. The puromycin-linked mRNA was then translated using the PURExpress ΔRF kit (New England Biolabs) with the addition of release factors 2 and 3 per the manufacturer’s instructions along with a “Solution A” containing 19 amino acids (-Met) (prepared as previously described) and either tRNA^fMet^_Ini_-*N*-chloroacetyl-L-Tyr or tRNA^fMet^_Ini_-*N*-chloroacetyl-D-Tyr (25 μM) to initiate translation and facilitate peptide macrocyclization. Translation was performed at 100 μL scale for the first round and 2.5 μL scale for subsequent rounds. Following translation, ribosomal denaturation was performed by addition of EDTA before reverse transcription with RNase H- reverse transcriptase. After exchange into selection buffer (50 mM Tris-HCl pH 8.0, 150 mM NaCl, 0.1% Tween-20) through G-25 resin the peptide-mRNA:cDNA libraries were incubated with cross-linked linked SARS-CoV-2 M^pro^ (200 nM) immobilized on Dynabeads™ His-tag Isolation & Pulldown (Life Technologies) at 4 °C for 30 min after which the beads were washed with ice-cold selection buffer (3 x 100 μL). The beads were then suspended in aqueous 0.1 vol.% Triton™ X-100 (100 μL) and heated to 95 °C for 5 min to recover binding peptides. The recovered cDNA was amplified by PCR, purified by ethanol precipitation and transcribed with T7 RNA polymerase to yield an enriched mRNA library to enter the following round of selection. Subsequent rounds were performed in the same manner with the addition of six negative selection steps against beads alone to remove bead binding sequences. Sequencing of recovered cDNA from rounds seven through nine was conducted using an iSeq high-throughput sequencer (Illumina).

### Cyclic peptide synthesis

For materials and methods, full synthetic details and characterization (HPLC, ESI-MS) of all peptides, see Supplementary Methods and Supplementary Figures 14-19.

### SARS-CoV-2 M^pro^ activity assay

The M^pro^ inhibition assay protocol was adapted and modified from ^27,47^ and was carried out with a reaction volume of 100 μL in 96-well-plates (black, polypropylene, U-bottom; Greiner Bio-One, Austria) using an aqueous buffer composed of 20 mM Tris-HCl pH 7.3, 100 mM NaCl, 1 mM EDTA, 1 mM DTT. All measurements were performed in triplicate. The compounds were incubated with recombinant SARS-CoV-2 M^pro^ for 10 min at 37 °C. The enzymatic reaction was initiated by addition of the FRET substrate (DABCYL)-KTSAVLQ↓SGFRKM-E(EDANS)-NH_2_ (Mimotopes, Australia). The final concentrations for IC_50_ determination amounted to 25 nM M^pro^ and 25 μM substrate, with inhibitor concentrations ranging from nanomolar to micromolar. The final concentrations for *K*_i_ determination amounted to 12.5 nM enzyme and 10, 20, 35 and 50 μM substrate, with a control and three inhibitor concentrations ranging from nanomolar to micromolar. The fluorescence signal was monitored by a fluorophotometer (Infinite 200 PRO M Plex; Tecan, Switzerland) for 5 min at 490 nm, using an excitation wavelength of 340 nm. Initial velocities were derived from the linear range of the enzymatic reaction. For IC_50_ determination, 100% enzymatic activity was defined as the initial velocity of control triplicates containing no inhibitor and the percentage of inhibition was calculated in relation to 100% enzymatic activity. An EDANS standard curve generated as described by Ma *et al.^35^* was used to convert fluorescence intensity to the amount of cleaved substrate (calibration curve). Data sets were analyzed with Prism 9.2 (Graph Pad Software, USA) to generate dose-response curves and calculate IC_50_ values, as well as to generate Michaelis-Menten curves and calculate *K*_i_ values assuming the appropriate inhibition mode.

### Crystallography

To obtain a crystallographic complex of SARS-CoV-2 M^pro^ and **1**, the peptide was dissolved in DMSO to a stock concentration of 50 mM. Protein, prepared as described above, was diluted to 4 mg/mL in TBS (50 mM Tris-HCl pH 7.5, 300 mM NaCl) and incubated with 2.5-fold molar excess (~300 μM) **1** for 2 hours on ice to saturate the binding sites of the protease. The complex was briefly buffer exchanged to remove unbound peptide using an Amicon Ultra 0.5 mL centrifugal filter with a 10 kDa MWCO to a final protein concentration of 4 mg/mL. This complex was used for high-throughput crystallography trials at 18 °C using drop sizes of 0.5 μL protein and 0.5 μL reservoir, which yielded a hit in the Index sparse matrix screen (Hampton Research). This hit was optimized but did not yield crystals of suitable diffraction quality; however, the crystals were able to be crushed and used to seed crystals with better morphology. Serial seeding was performed using a protein concentration of 1 mg/mL for several rounds until crystals of suitable diffracting quality were obtained, which were formed at 18 °C in a drop size of 1 μL reservoir, 1.5 μL protein, and 0.5 μL seed stock against a reservoir solution of 25% PEG 3350, 0.1 M Bis-Tris pH 6.5, 0.3 M NaCl. Crystals were flash frozen without cryoprotecting and diffraction data were collected at 100 K using the MX2 beamline at the Australian Synchrotron. As the diffraction was not adequate to unambiguously model the inhibitor into the crystal structure, we solved the co-crystal structure of the SARS-CoV-2 M^pro^-**Se-1** complex. The complex was prepared the same as with **1** but with overnight incubation at 4 °C. The complex was diluted to a protein concentration of 1 mg/mL and crystallized at 18 °C in a drop size of 1 μL reservoir, 1.5 μL protein and 0.5 μL seed stock against a reservoir solution of 22% PEG 3350, 0.1 M Bis-Tris pH 6.0, 0.3 M NaCl. Crystals were seeded using crystal seeds of the SARS-CoV-2 M^pro^-**1** crystals. Crystals formed as thin plates after ~4–5 days and were flash frozen without cryoprotection.

Diffraction data were collected at 100 K using the MX2 beamline at the Australian Synchrotron^59^. Reflections collected were indexed and integrated using XDS^60^ and scaled in Aimless (CCP4)^61^. The phase problem was overcome by molecular replacement in Phaser MR (CCP4)^62^, using PDB ID 7JKV as the search model. The structure was refined by iterative rounds of rebuilding in Coot ^63^, and twin refinement in Refmac^64,65^. The crystal was twinned P21 (-h, -k, (k+l)); initially autoindexing as C221). Data collection and refinement statistics are given in **Supplementary Table 3**. The final structure was deposited in the Protein Data Bank (PDB: 7RNW).

### SARS-CoV-2 Infectivity Assays

HEK293-ACE2-TMPRSS2 cells stably expressing human ACE2 and TMPRSS2 were generated as previously described^66^. A high content fluorescence microscopy approach was used to assess the ability of the cyclic peptide M^pro^ inhibitors to protect cells from SARS-CoV-2 induced cytopathic effects in permissive cells. In brief, the engineered cell line succumbs to viral cytopathic effects after 6 to 18 hours post infection. Cytopathic effects can be enumerated, as cells and their nuclei collapse into large syncytia after 18 hours of viral culture. The remaining cells outside of the syncytia increase in a dose dependent manner the lower the viral titers are and/or if a viral inhibitor is introduced within the culture.

For testing the cyclic peptides, compounds were initially diluted in cell culture medium (DMEM-5% FCS) to make 4x working stock solutions and then serially diluted further in the above media to achieve a 2-fold dilution series. On the day of the assay, HEK293-ACE2-TMPRSS2 cells were trypsinized, stained with Nucblue in suspension and then seeded at 16,000 cells in a volume of 40 μL of DMEM-5% FCS per well in a 384-well plate (Corning #CLS3985). Diluted compounds (20 μL) were added to the cells and the plates containing cells and compounds incubated for 1 hour at 37 °C, 5% CO_2_. 20 μL of virus solution at 8×10^3^ TCID50/mL ^67^ was then added to the wells and plates were incubated for a further 24 hours at 37 °C, 5% CO_2_. Stained cells were then imaged using the InCell 2500 (Cytiva) high throughput microscope, with a 10× 0.45 NA CFI Plan Apo Lambda air objective. Acquired nuclei were counted using InCarta high-content image analysis software (Cytiva) to give a quantitative measure of CPE. Virus inhibition/neutralization was calculated as %N= (D-(1-Q))x100/D, where; “Q” is the value of nuclei in test well divided by the average number of nuclei in untreated uninfected controls, and “D”=1-Q for the wells infected with virus but untreated with inhibitors. Thus, the average nuclear counts for the infected and uninfected cell controls get defined as 0% and 100% neutralization respectively. To account for cell death due to drug toxicity, cells treated with a given compound alone and without virus were included in each assay. The % neutralization for each compound concentration in infected wells was normalized to % neutralization in wells with equivalent amount of compound but without the virus to yield the final neutralization values for each condition. Inhibition curves and 50% (EC_50_) effective concentrations were determined by non-linear regression analysis using GraphPad Prism software (version 9.1.2, GraphPad software, USA).

### Cytotoxicity and targeted proteomics on HEK293-ACE2-TMPRSS2 cell line

HEK293-ACE2-TMPRSS2 cells were maintained in Dulbecco’s Modified Eagle Medium (DMEM) with 4.5 g/L D-Glucose, L-Glutamine and 110 mg/L sodium pyruvate (Gibco), supplemented with 10% foetal calf serum (Hyclone). Cells were sub-cultured between 70-90% confluency and tested regularly to ensure free of mycoplasma contamination. Cytotoxicity of **1**, **pen-1**, **6** and **pen-6**, was determined by an Alamar Blue HS (Invitrogen) cell viability assay as per manufacturer’s instructions. Briefly, 5×10^4^ cells were seeded into wells of a 96-well flat bottom culture plate (Corning) and once adhered, compounds or vehicle control (DMSO) were added at varying concentrations and incubated for 24 hours (37 °C, 5% CO_2_). Alamar Blue HS cell viability reagent (Invitrogen) was added and the cells further incubated for 2-3 hours. Relative fluorescent units (RFU) were determined per well at ex/em 560/590 nm (Tecan Infinite M1000 pro plate reader). Increasing RFU is proportional to cell viability.

For targeted proteomics, cells in 6-well plates at 70% confluency were treated with vehicle control (DMSO) or inhibitors at 10 μM. At various time points (performed in triplicate), the cells were washed 3 times with 2 mL DPBS (Gibco), scraped and lysed in 4% SDC buffer (4% sodium deoxycholate, 0.1 M Tris-HCl pH 8.0) then immediately heated to 95 °C for 10 mins before freezing at −30 °C prior to processing for targeted proteomics^68^. Cell lysates were digested to peptides as described previously. Peptides were analyzed by LC-MS/MS using a data-independent acquisition method as described previously.^69^ Extracted ion chromatograms for fragment ions derived from tryptic peptides were plotted using Xcalibur Qual Browser (Thermo Scientific). The area under the curve for each peak was used for quantification.

## Data availability

The X-Ray co-crystal structure of SARS-CoV-2 M^pro^-**Se-1** has been deposited to the PDB (PDB ID: 7RNW). All other data supporting the findings reported in this study are available in the article and its Supplementary Information.

## Acknowledgements

We acknowledge the ARC Centre of Excellence for Innovations in Peptide and Protein Science (CE200100012 to G.O., C.J.J. and R.J.P.) for funding and the John A. Lamberton Research Scholarship (to J.J-L) and Research Training Program from the Australian government (to J.J-L., S.E.F. and M.J.B) for PhD scholarships. C.N. acknowledges ARC funding (DE190100015, DP200100348). This research was undertaken in part using cyclic peptide display screening facilities in Sydney Analytical, and the MX2 beamline at the Australian Synchrotron, part of ANSTO, and made use of the Australian Cancer Research Foundation (ACRF) detector. Next generation sequencing was conducted through the Ramaciotti Centre for Genomics. M.L. is a Cancer Institute New South Wales Future Research Leader Fellow. We thank SydneyMS for providing the instrumentation used in this study. We also acknowledge Karishma Patel for her assistance in the use of SEC-MALLS.

## Author contributions

J.J-L., G.O., S.T., C.J.J. C.N. and R.J.P conceived the project. J.J-L. performed protein cross-linking and SEC-MALLS. J.J-L. and T.P. performed RaPID selection experiments. J.J-L. S.E.F. and M.J.B performed peptide synthesis. S.U. performed all M^pro^ inhibition assays. K.B.E and M.C.E. produced wild-type and mutant variants of SARS-CoV-2 M^pro^ and performed NMR assignments. R.F. performed M^pro^ crystallographic studies. V.M.S. performed molecular dynamics simulations. A.A. performed all SARS-CoV-2 inhibition assays. A.S.A assessed compounds for cytotoxicity and conducted cellular uptake experiments. M.L. performed proteomics experiments. J.J-L., G.O., S.T., C.J.J. C.N. and R.J.P wrote the paper with contributions from all authors.

