## Supplementary Figures and Materials for "Discovery of Antiviral Cyclic Peptides Targeting the Main Protease of SARS-CoV-2 *via* mRNA Display"

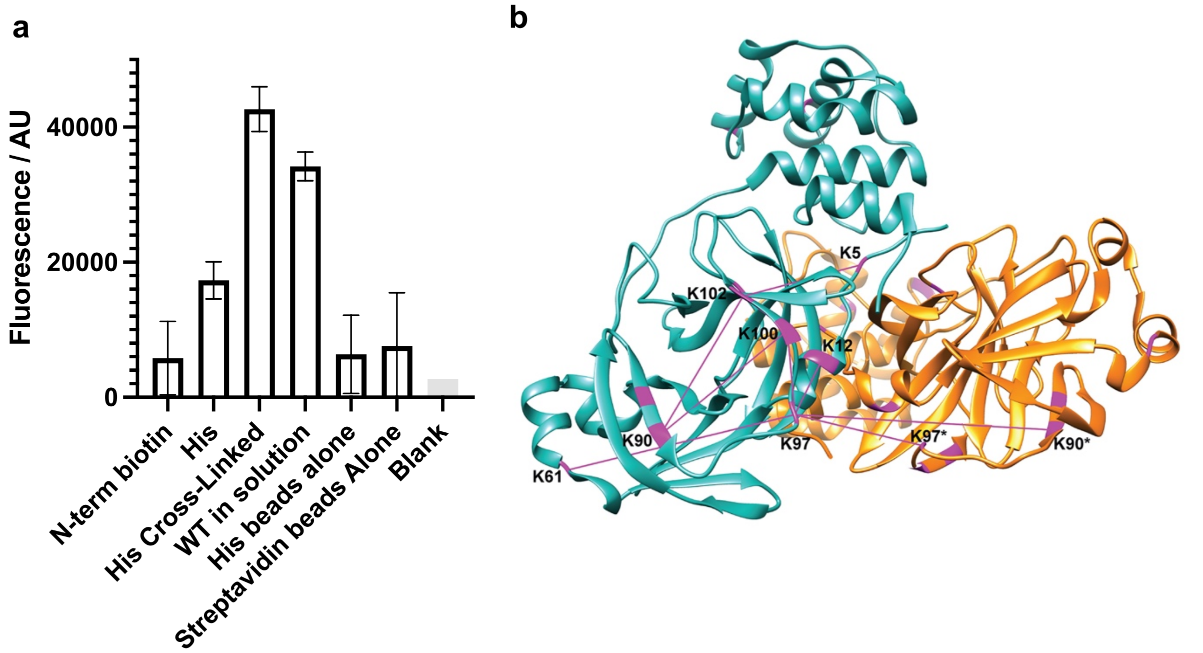

**Supplementary Figure 1. a** Catalytic activity of different SARS-CoV-2 M$^{pro}$ constructs including: N-terminally biotinylated M$^{pro}$ immobilized to Streptavidin Dynabeads$^{TM}$ (N-term biotin), C-terminally His-tagged M$^{pro}$ immobilized onto $Co^{2+}$-NTA Dynabeads$^{TM}$ (His), DSG cross-linked and C-terminally His-tagged M$^{pro}$ immobilized onto $Co^{2+}$-NTA Dynabeads$^{TM}$ (His Cross-Linked), wild-type M$^{pro}$ in solution (WT in solution) All experiments were performed with 25 nM M$^{pro}$ and 20 µM FRET substrate (DABCYL)-KTSAVLQ↓SGFRKM-E(EDANS)-$NH_2$ (Mimotopes, Australia) with incubation at 37 °C for 15 min. **b** Identification of DSG crosslinking sites in SARS-CoV-2 M$^{pro}$ was performed after trypsin digestion by mass spectrometry using the Byonic search engine (Protein Metrics). Crosslinks <30 Å are displayed on the dimeric structure of SARS-CoV-2 M$^{pro}$ (PDB: 6Y2E) as purple lines connecting lysine residues (all shown in purple on ribbon) using UCSF Chimera.[1] All intramolecular crosslinks are shown only on one monomer (cyan) with residue numbers indicated. Intermolecular crosslinks are shown with linked lysines in the second monomer (orange) indicated by an asterisk.

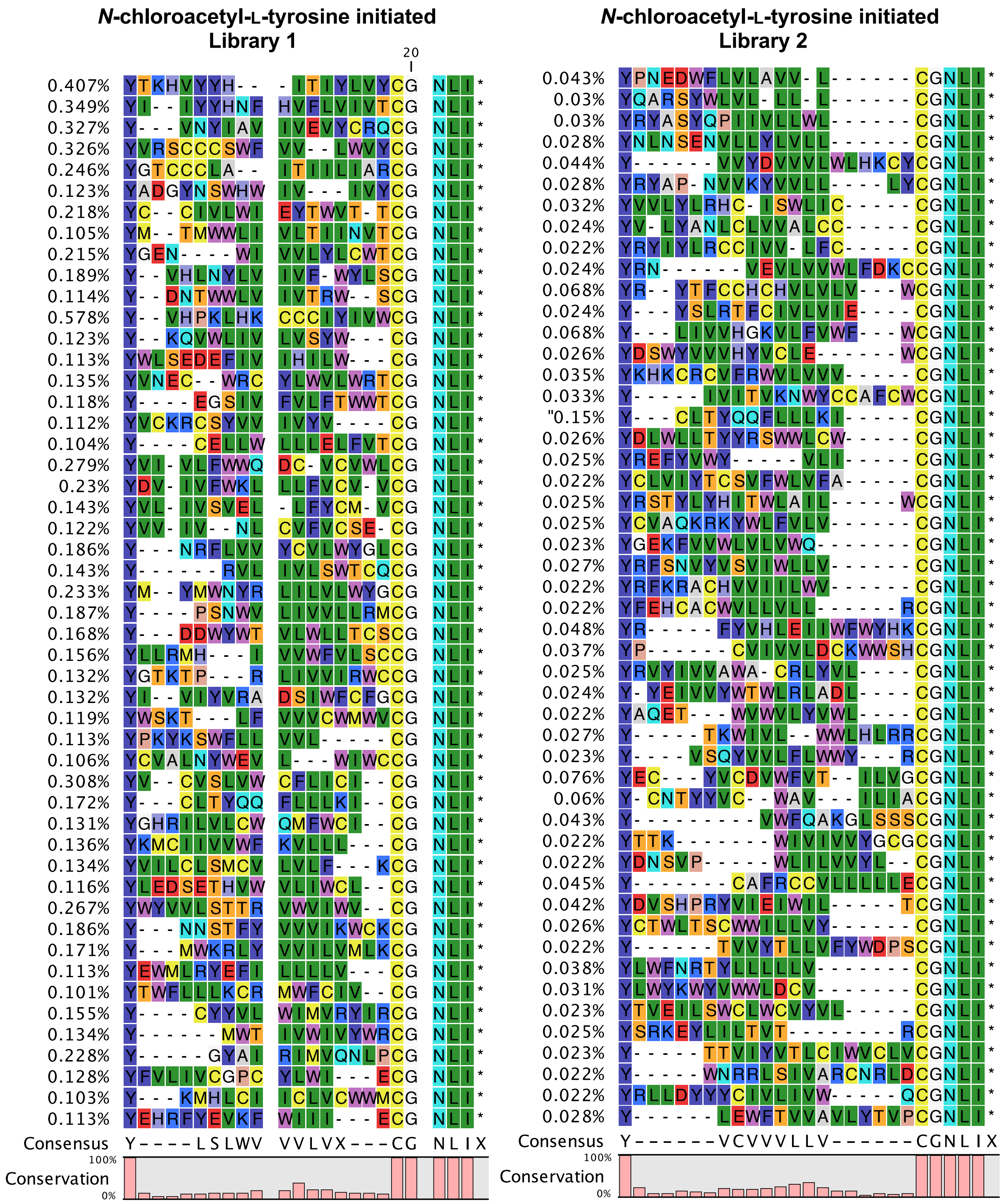
N-chloroacetyl-L-tyrosine initiated
Library 1
20
0.407% YTKHVYYH-- -ITIYLVYCG NLI*
0.349% YI--IYYHNF HVFLVIVTCG NLI*
0.327% Y---VNYIAV IVEVYCRQCG NLI*
0.326% YVRSCCCSWF VV--LWVYCG NLI*
0.246% YGTCCCLA-- ITIILIARCG NLI*
0.123% YADGYNSWHW IV---IVYCG NLI*
0.218% YC--CIVLWI EYTWVT-TCG NLI*
0.105% YM--TMWWLI VLTIINVTCG NLI*
0.215% YGEN----WI VVLYLCWTCG NLI*
0.189% Y--VHLNYLV IVF-WYLSCG NLI*
0.114% Y--DNTWWLV IVTRW--SCG NLI*
0.578% Y--VHPKLHK CCCIYIVWCG NLI*
0.123% Y--KQVWLIV LVSYW---CG NLI*
0.113% YWLSEDEFIV IHILW---CG NLI*
0.135% YVNEC--WRC YLWVLWRTCG NLI*
0.118% Y----EGSIV FVLFTWWTCG NLI*
0.112% YVCKRCSYVV IVYV----CG NLI*
0.104% Y----CELLW LLLELFVTCG NLI*
0.279% YVI-VLFWWQ DC-VCVWLCG NLI*
0.23% YDV-IVFWKL LLFVCV-VCG NLI*
0.143% YVL-IVSVEL -LFYCM-VCG NLI*
0.122% YVV-IV--NL CVFVCSE-CG NLI*
0.186% Y---NRFLVV YCVLWYGLCG NLI*
0.143% Y------RVL IVLSWTCQCG NLI*
0.233% YM--YMWNYR LILVLWYGCG NLI*
0.187% Y----PSNWV LIVVLLRMCG NLI*
0.168% Y---DDWYWT VLWLLTCSCG NLI*
0.156% YLLRMH---I VVWFVLSCCG NLI*
0.132% YGTKTP---R LIVVIRWCCG NLI*
0.132% YI--VIYVRA DSIWFCFGCG NLI*
0.119% YWSKT---LF VVVCWMWVCG NLI*
0.113% YPKYKSWFLL VVL-----CG NLI*
0.106% YCVALNYWEV L---WIWCCG NLI*
0.308% YV--CVSLVW CFLICI--CG NLI*
0.172% Y---CLTYQQ FLLLKI--CG NLI*
0.131% YGHRILVLCW QMFWCI--CG NLI*
0.136% YKMCIIVVWF KVLLL---CG NLI*
0.134% YVILCLSMCV LVLF---KCG NLI*
0.116% YLEDSETHVW VLIWCL--CG NLI*
0.267% YWYVVLSTTR VWVIWV--CG NLI*
0.186% Y---NNSTFY VVVIKWCKCG NLI*
0.171% Y---MWKRLY VVILVMLKCG NLI*
0.113% YEWMLRYEFI LLLLV---CG NLI*
0.101% YTWFLLLKCR MWFCIV--CG NLI*
0.155% Y----CYYVL WIMVRYIRCG NLI*
0.134% Y------MWT IVWIVYWRCG NLI*
0.228% Y-----GYAI RIMVQNLPCG NLI*
0.128% YFVLIVCGPC YLWI---ECG NLI*
0.103% Y---KMHLCI ICLVCWWMCG NLI*
0.113% YEHRFYEVKF WIII---ECG NLI*
Consensus Y----LSLWV VVLVX---CG NLIX
100%
Conservation
0%
N-chloroacetyl-L-tyrosine initiated
Library 2
0.043% YPNEDWFLVLAVV-L------CGNLI*
0.03% YQARSYWLVL-LL-L------CGNLI*
0.03% YRYASYQPIIVLLWL------CGNLI*
0.028% YNLNSENVLLYLVLL------CGNLI*
0.044% Y------VVYDVVVLWLHKCYCGNLI*
0.028% YRYAP-NVVKYVVLL----LYCGNLI*
0.032% YVVLYLRHC-ISWLIC-----CGNLI*
0.024% YV-LYANLCLVVALCC-----CGNLI*
0.022% YRYIYLRCCIVV-LFC-----CGNLI*
0.024% YRN------VEVLVVWLFDKCCGNLI*
0.068% YR--YTFCCHCHVLVLV---WCGNLI*
0.024% Y---YSLRTFCIVLVIE----CGNLI*
0.068% Y---LIVVHGKVLFVWF---WCGNLI*
0.026% YDSWYVVVHYVCLE------WCGNLI*
0.035% YKHKCRCVFRWVLVVV-----CGNLI*
0.033% Y-----IVITVKNWYCCAFCWCGNLI*
"0.15% Y---CLTYQQFLLLKI-----CGNLI*
0.026% YDLWLLTYYRSWWLCW-----CGNLI*
0.025% YREFYVWY-----VLI-----CGNLI*
0.022% YCLVIYTCSVFWLVFA-----CGNLI*
0.025% YRSTYLYHITWLAIL-----WCGNLI*
0.025% YCVAQKRKYWLFVLV------CGNLI*
0.023% YGEKFVVVWLVLVWQ------CGNLI*
0.027% YRFSNVYVSVIWLLV------CGNLI*
0.022% YRFKRACHVVIILWV------CGNLI*
0.022% YFEHCACWVLLVLL------RCGNLI*
0.048% YR-----FYVHLEIIWFWYHKCGNLI*
0.037% YP------CVIVVLDCKWWSHCGNLI*
0.025% YRVYIVVAWA-CRLYVL----CGNLI*
0.024% Y-YEIVVYWTWLRLADL----CGNLI*
0.022% YAQET---WVWVLYVWL----CGNLI*
0.027% Y-----TKWIVL--WWLHLRRCGNLI*
0.023% Y----VSQYVVLFLWWY---RCGNLI*
0.076% YEC---YVCDVWFVT--ILVGCGNLI*
0.06% Y-CNTYYVC--WAV---ILIACGNLI*
0.043% Y---------VWFQAKGLSSSCGNLI*
0.022% YTTK-------WIVIVVYGCGCGNLI*
0.022% YDNSVP-----WLILVVYL--CGNLI*
0.045% Y-------CAFRCCVLLLLLECGNLI*
0.042% YDVSHPRYVIEIWIL-----TCGNLI*
0.026% YCTWLTSCWWILLVY------CGNLI*
0.022% Y------TVVYTLLVFYWDPSCGNLI*
0.038% YLWFNRTYLLLLLV-------CGNLI*
0.031% YLWYKWYVWWLDCV-------CGNLI*
0.023% YTVEILSWCLWCVYVL-----CGNLI*
0.025% YSRKEYLILTVT--------RCGNLI*
0.023% Y-----TTVIYVTLCIWVCLVCGNLI*
0.022% Y-----WNRRLSIVARCNRLDCGNLI*
0.022% YRLLDYYYCIVLIVW-----QCGNLI*
0.028% Y------LEWFTVVAVLYTVPCGNLI*
Consensus Y------VCVVVLLV------CGNLIX
100%
Conservation
0%

***N*-chloroacetyl-D-tyrosine initiated**

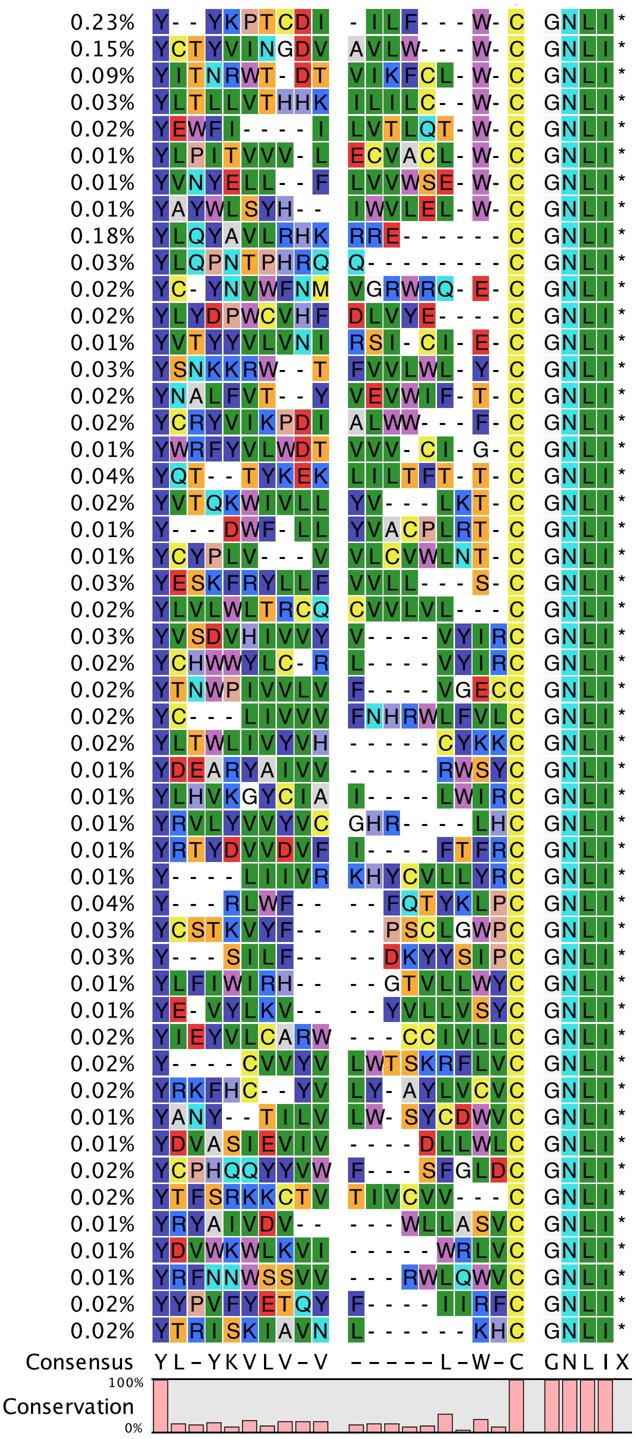

**Supplementary Figure 2. Top 50 sequences for each selection based on percentage enrichment in the final library after round 9.**

**a**

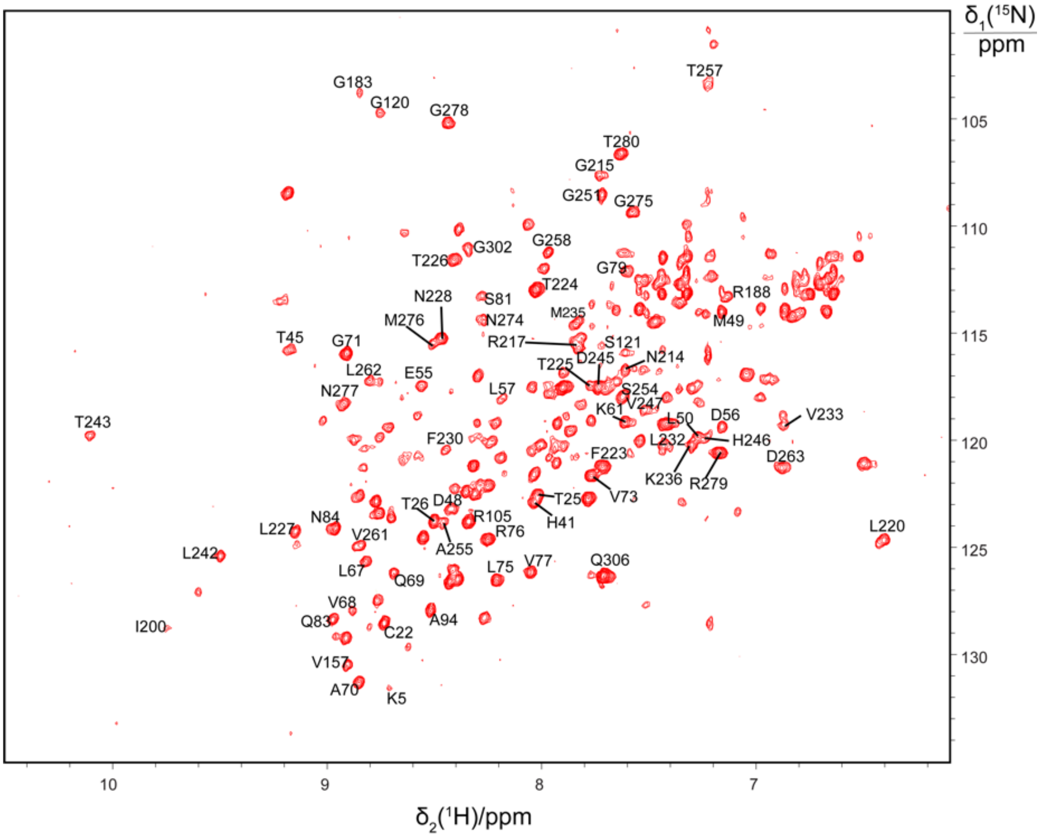

δ1(15N)/ppm
δ2(1H)/ppm
105
110
115
120
125
130
10
9
8
7
G183
G120
G278
T257
T280
G215
G251
G275
G258
G302
T226
G79
T224
N228
S81
N274
M276
M235
R188
M49
T45
G71
R217
S121
T225
D245
N214
L262
E55
N277
L57
S254
V247
V233
T243
K61
L50
D56
H246
L232
D263
F230
F223
K236
R279
V73
T26
D48
T25
R105
H41
L227
N84
R76
L220
V261
A255
L242
L67
Q69
L75
V77
Q306
V68
Q83
A94
I200
C22
V157
A70
K5

**b**

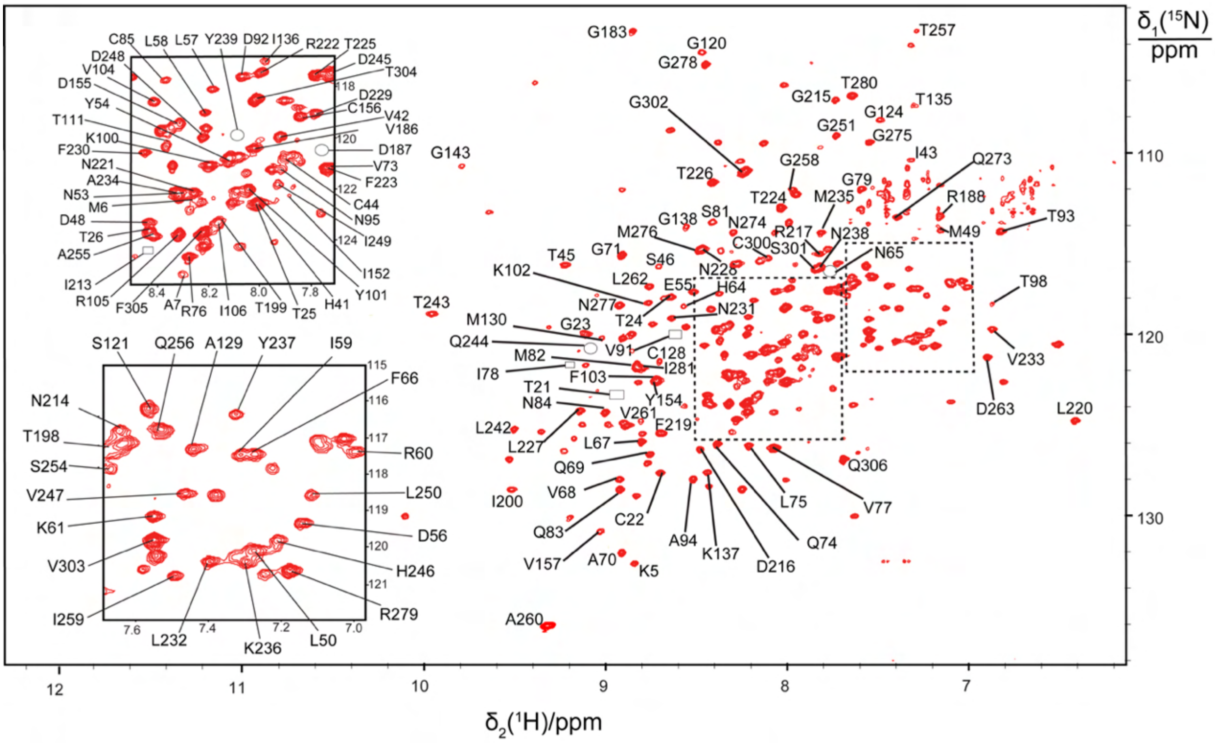

δ1(15N)/ppm
δ2(1H)/ppm
12
11
10
9
8
7
110
120
130
G183
G120
G278
T257
T280
G215
T135
G124
G251
G302
G275
I43
G143
G258
Q273
T226
G79
T224
M235
R188
G138
S81
N274
R217
N238
M49
T93
M276
C300
S301
N65
T45
G71
S46
N228
K102
L262
H64
T98
E55
N277
T243
N231
T24
M130
G23
Q244
V91
C128
V233
M82
I281
I78
F103
D263
T21
Y154
L220
N84
V261
F219
L242
L227
L67
Q69
Q306
V68
I200
L75
V77
C22
Q83
A94
K137
Q74
V157
A70
K5
D216
A260
C85
L58
L57
Y239
D92
I136
R222
T225
D248
D245
V104
T304
D155
D229
Y54
C156
V42
T111
V186
K100
D187
F230
N221
V73
A234
F223
N53
C44
M6
N95
D48
I249
T26
A255
I152
I213
Y101
R105
F305
A7
R76
I106
T199
T25
H41
S121
Q256
A129
Y237
I59
N214
F66
T198
S254
R60
V247
L250
K61
D56
V303
H246
I259
L232
K236
L50
R279

**c**

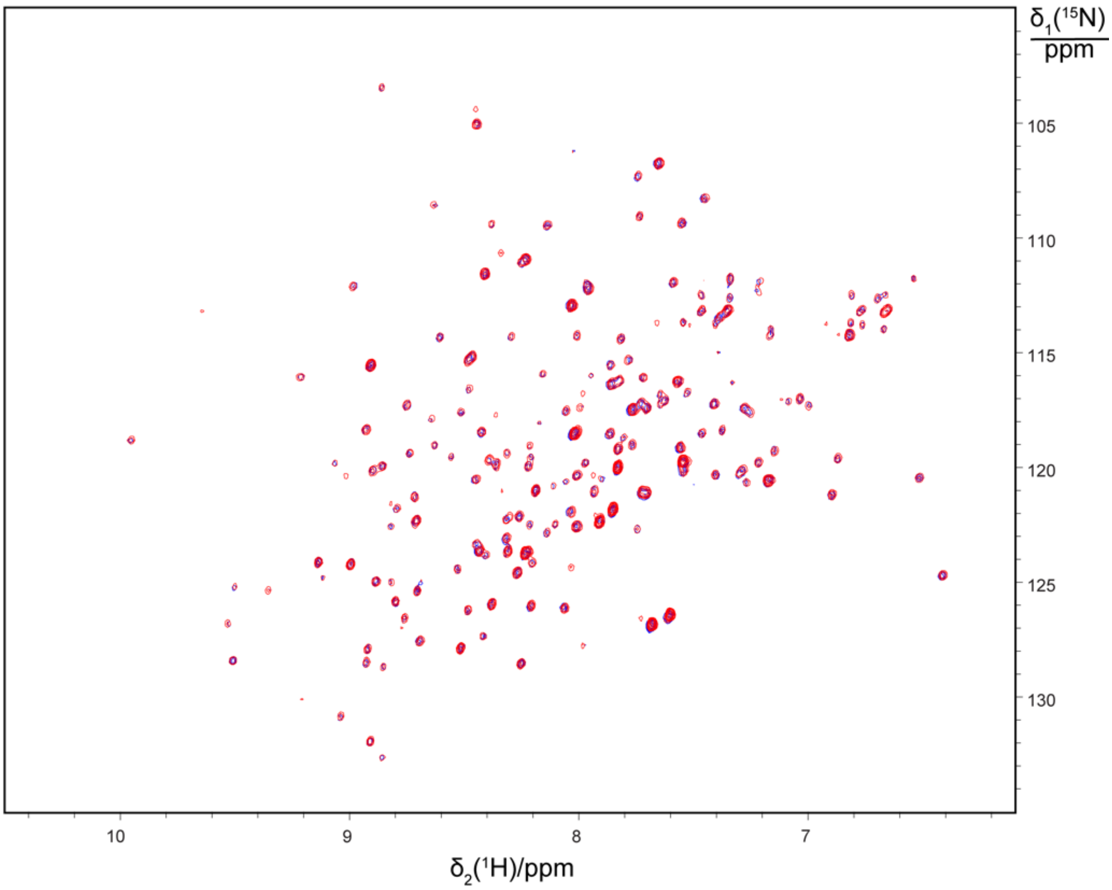

**d**

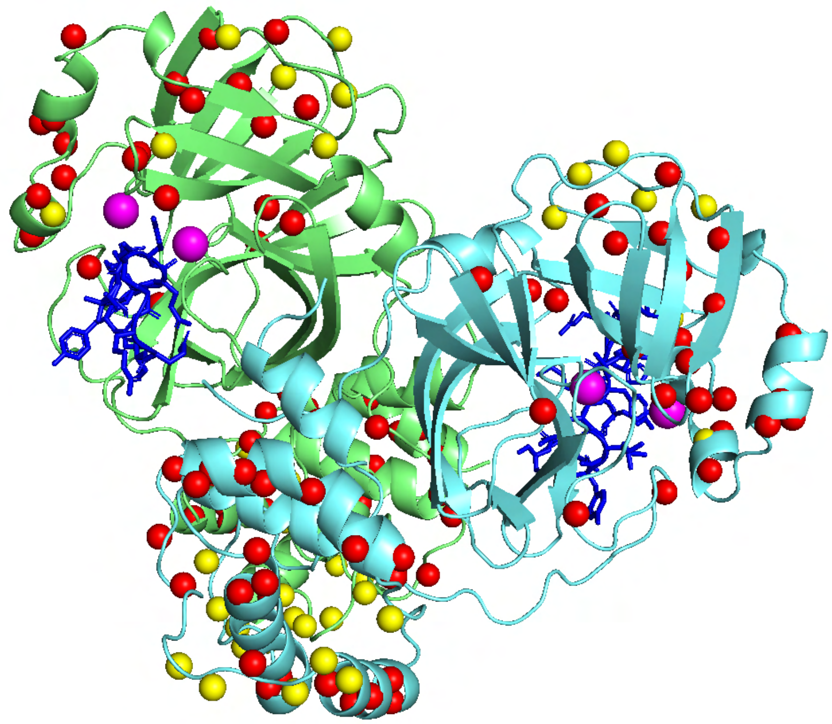

**Supplementary Figure 3. a Projection onto the $^{15}N$-$^{1}H$ plane of 3D TROSY-HNCO spectra recorded of wild-type $M^{pro}$.** Assignments are shown for peaks that could be assigned by comparison with the assignments made in the monomeric $M^{pro}$ R298A mutant. **b [$^{15}N$,$^{1}H$]-TROSY spectrum of a 0.3 mM solution of uniformly $^{15}N$-labelled $M^{pro}$ R298A recorded at 25 °C ($t_{1max}$ = 82 ms, $t_{2max}$ = 142 ms, total recording time 2.2 h).** The peaks are labelled with the residue type and amino acid sequence number of the assignments made. Some cross-peaks were observed and assigned in 3D NMR spectra or in [$^{15}N$,$^{1}H$]-HSQC spectra of selectively $^{15}N$-labelled samples, but not observed in the TROSY spectrum shown here. Their positions are marked by symbols identifying the location and the spectrum where they were observed: ○ peak found in TROSY-HNCA or TROSY-HNCO spectrum; □ peak found in [$^{15}N$,$^{1}H$]-HSQC spectrum of selectively $^{15}N$-labelled samples. **c Projection onto the $^{15}N$-$^{1}H$ plane of 3D TROSY-HNCO spectra recorded of 0.3 mM solutions of $^{15}N$/$^{13}C$/$^{2}H$-labelled $M^{pro}$ R298A.** Two projections are superimposed, recorded of samples without (blue contours) and with equimolar inhibitor (red contours). The spectrum recorded in the presence of inhibitor superimposes almost perfectly the spectrum without inhibitor, indicating that the inhibitor fails to bind and cause any spectral changes. **d** Crystal structure (PDB ID: 7RNW) of the dimer of wild-type $M^{pro}$ showing the location of backbone amide protons which changed in the NMR spectra upon titration with the inhibitor. Red spheres indicate amides with significant changes in chemical shift or intensity in the presence of inhibitor. Yellow spheres indicate amides remaining unchanged in the presence of inhibitor. The locations of the active site residues His41 and Cys145 are highlighted in magenta and the inhibitor **Se-1** in blue.

**Supplementary Table 1.** Number of specific resonance assignments made in [$^{15}N$,$^{1}H$]-HSQC spectra by site-directed mutagenesis.

| Residue | Mutated to | Number of mutants | Peaks assigned |
|---|---|---|---|
| glycine | alanine | 19 | 8 |
| threonine | serine | 11 | 8 |
| isoleucine | valine | 6 | 4 |
| valine | isoleucine | 9 | 5 |
| leucine | alanine | 11 | 6 |
| lysine | arginine | 7 | 3 |
| cysteine | serine | 5 | 3 |
| methionine | alanine | 5 | 2 |
| serine | alanine | 6 | 1 |
| alanine | glycine | 4 | 0 |

**Supplementary Table 2.** Compilation of [$^{15}$N,$^{1}$H]-HSQC spectra recorded of selectively $^{15}$N-labelled samples of M$^{pro}$ R298A and mutants enabling specific resonance assignments.[a]

| Panel of Supplementary Figure 4 | Residue type | Sequence number | Total recording time / h |
|---|---|---|---|
| 01 | glycine | 23 | 1.3 |
| 02 | glycine | 71 | 1.3 |
| 03 | glycine | 79 | 1.3 |
| 04 | glycine | 124 | 1.3 |
| 05 | glycine | 138 | 1.3 |
| 06 | glycine | 183 | 2.7 |
| 07 | glycine | 251 | 1.3 |
| 08 | glycine | 278 | 2.7 |
| 09 | threonine | 21 | 1.3 |
| 10 | threonine | 24 | 1.3 |
| 11 | threonine | 25 | 1.3 |
| 12 | threonine | 45 | 1.3 |
| 13 | threonine | 93 | 1.3 |
| 14 | threonine | 98 | 2.7 |
| 15 | threonine | 111 | 1.3 |
| 16 | threonine | 199 | 1.3 |
| 17 | isoleucine | 59 | 1.3 |
| 18 | isoleucine | 78 | 1.3 |
| 19 | isoleucine | 106 | 1.3 |
| 20 | isoleucine | 200 | 1.3 |
| 21 | valine | 68 | 1.3 |
| 22 | valine | 73 | 1.3 |
| 23 | valine | 77 | 1.3 |
| 24 | valine | 91 | 1.3 |
| 25 | valine | 104 | 1.3 |
| 26 | leucine | 50 | 10.6 |
| 27 | leucine | 58 | 13.3 |
| 28 | leucine | 67 | 1.3 |
| 29 | leucine | 75 | 1.3 |
| 30 | leucine | 220 | 1.3 |
| 31 | leucine | 232 | 1.3 |
| 32 | lysine | 100 | 7.3 |
| 33 | lysine | 102 | 2.7 |
| 34 | lysine | 137 | 7.3 |
| 35 | cysteine | 85 | 5.3 |
| 36 | cysteine | 128 | 1.3 |
| 37 | cysteine | 156 | 1.3 |
| 38 | methionine | 6 | 1.3 |
| 39 | methionine | 276 | 2.7 |
| 40 | serine | 46 | 1.3 |

[a] The table columns refer to the number of the spectrum, the residue type labelled with $^{15}$N, the amino acid sequence number of the assignment made and the total recording

time of the spectrum. The individual spectra are shown in Supplementary Figure 4 below.

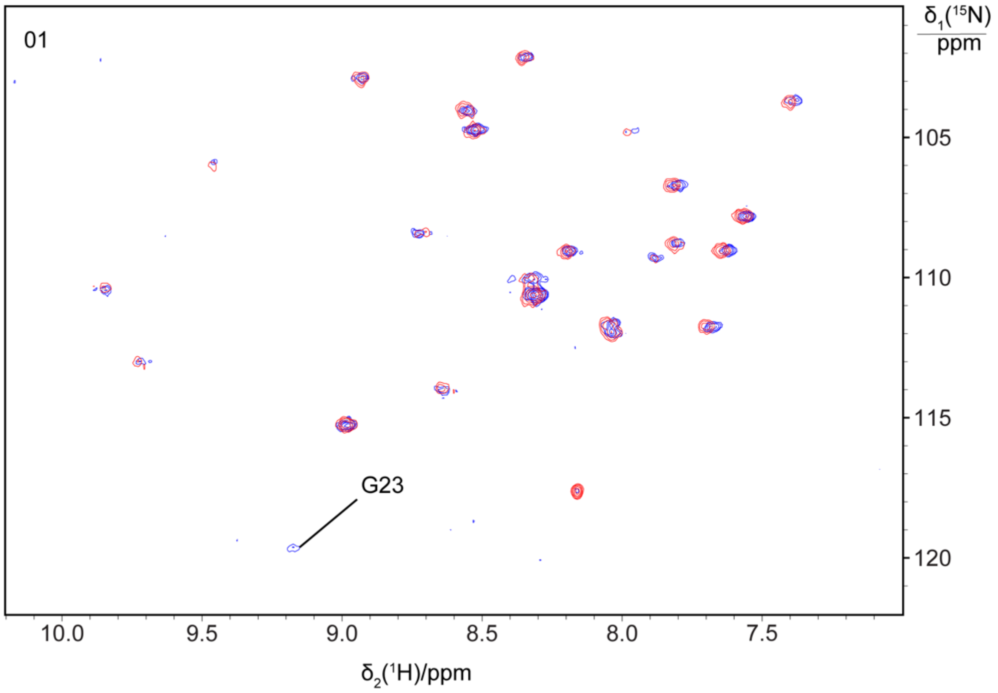

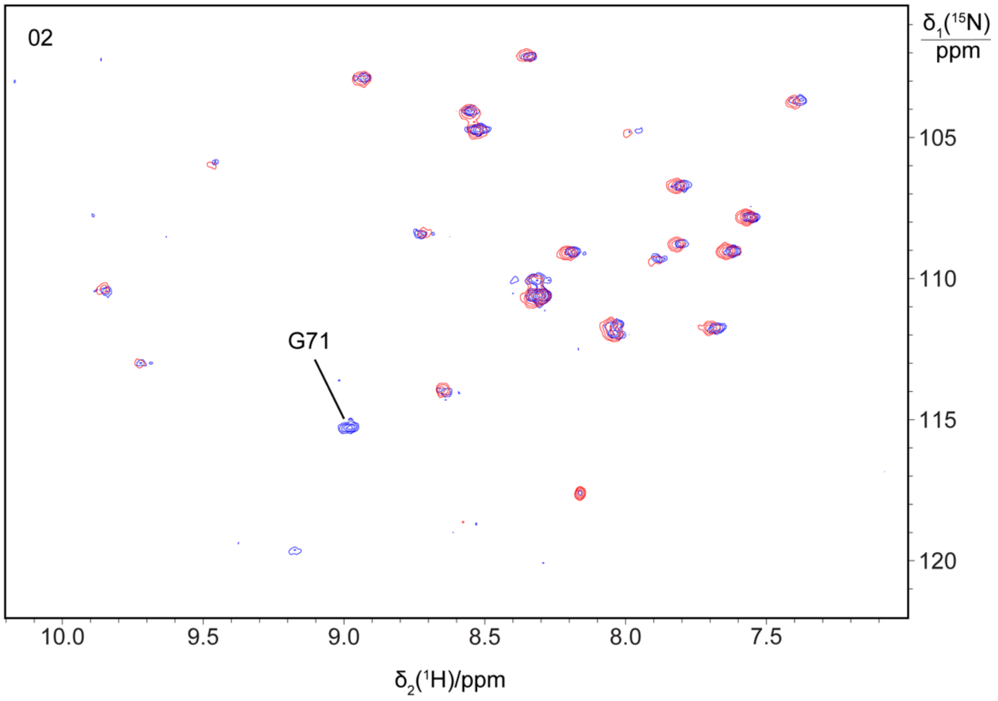

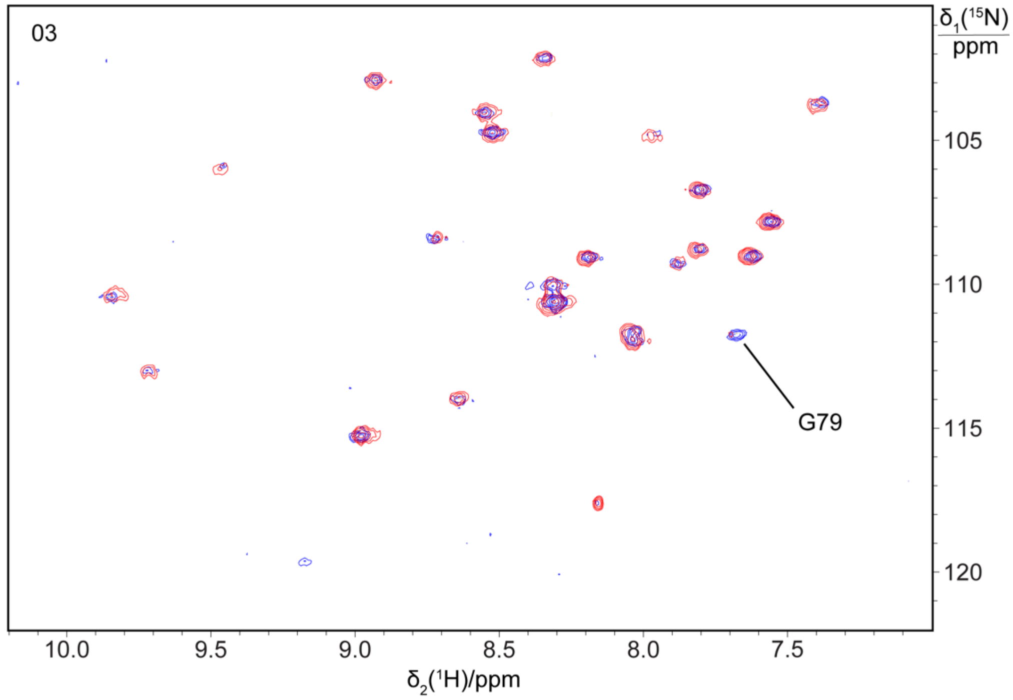
03
G79
δ1(15N)/ppm
105
110
115
120
10.0
9.5
9.0
8.5
8.0
7.5
δ2(1H)/ppm

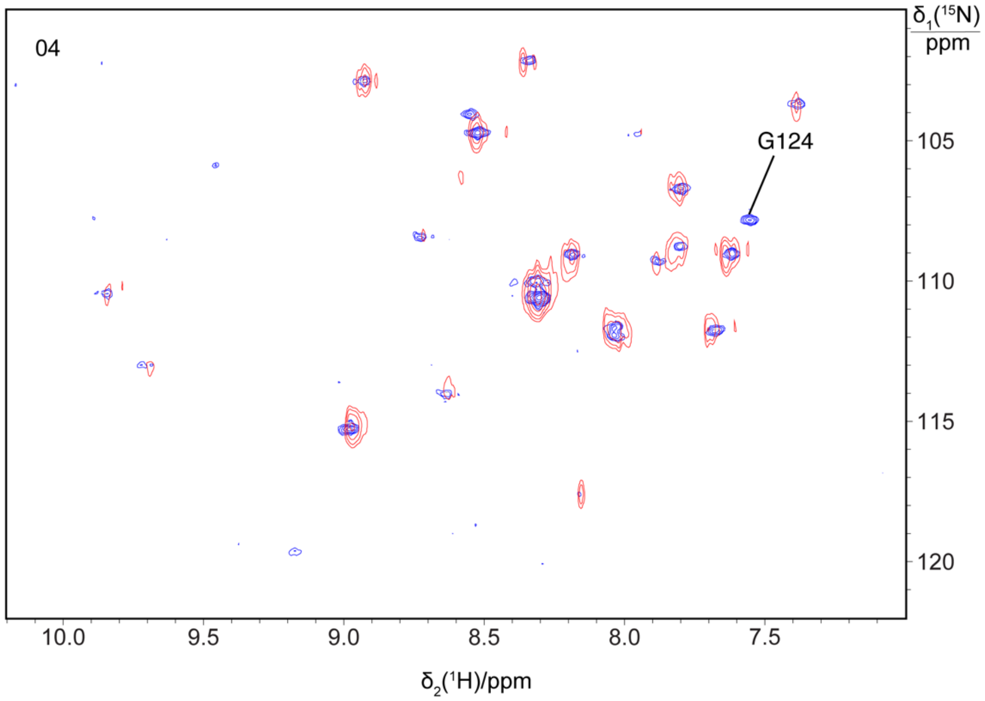
04
G124
δ1(15N)/ppm
105
110
115
120
10.0
9.5
9.0
8.5
8.0
7.5
δ2(1H)/ppm

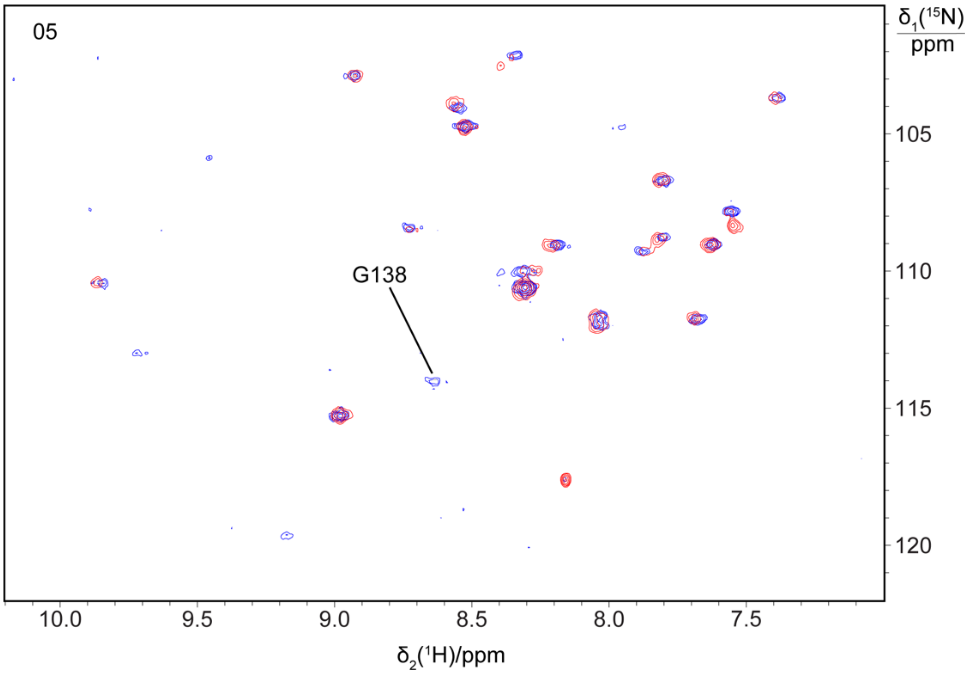
05
G138
δ1(15N)
ppm
105
110
115
120
10.0
9.5
9.0
8.5
8.0
7.5
δ2(1H)/ppm

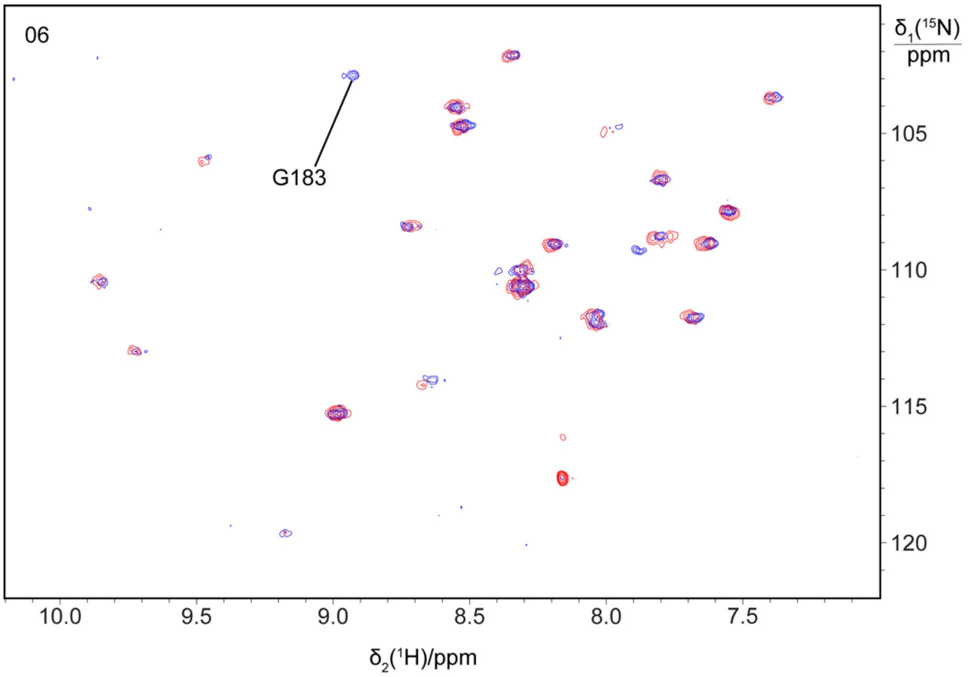
06
G183
δ1(15N)
ppm
105
110
115
120
10.0
9.5
9.0
8.5
8.0
7.5
δ2(1H)/ppm

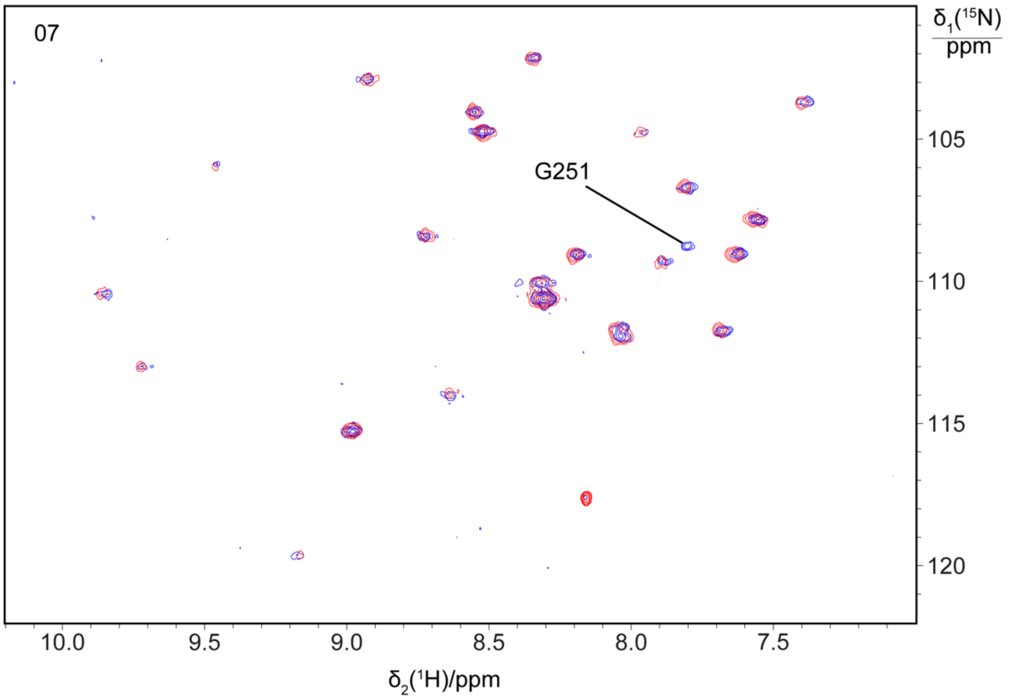
07
G251
δ1(15N)
ppm
105
110
115
120
10.0
9.5
9.0
8.5
8.0
7.5
δ2(1H)/ppm

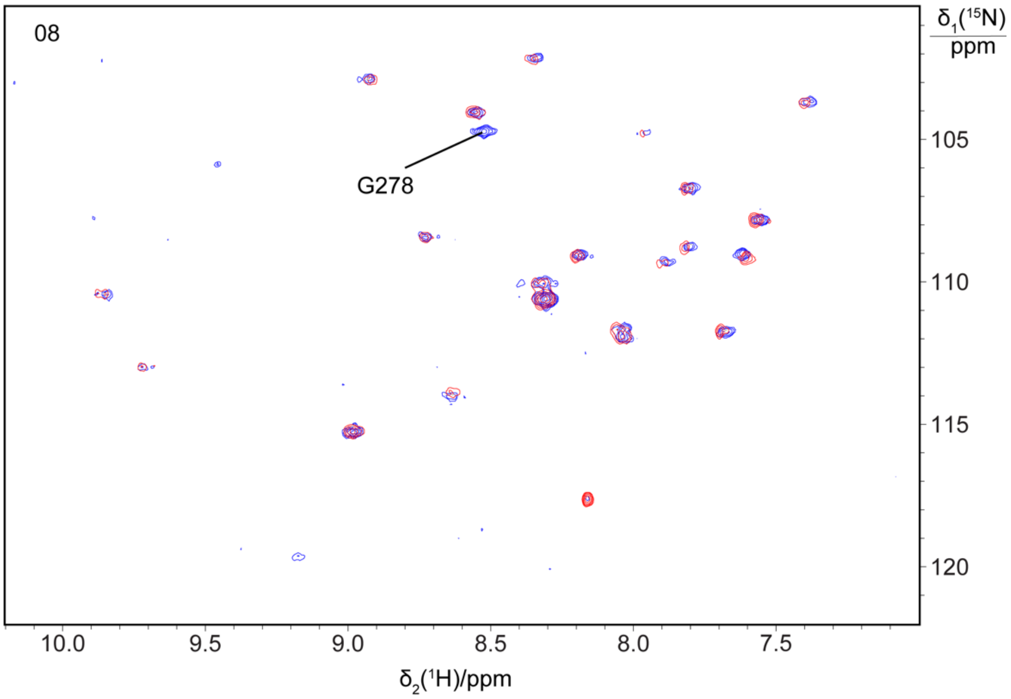
08
G278
δ1(15N)
ppm
105
110
115
120
10.0
9.5
9.0
8.5
8.0
7.5
δ2(1H)/ppm

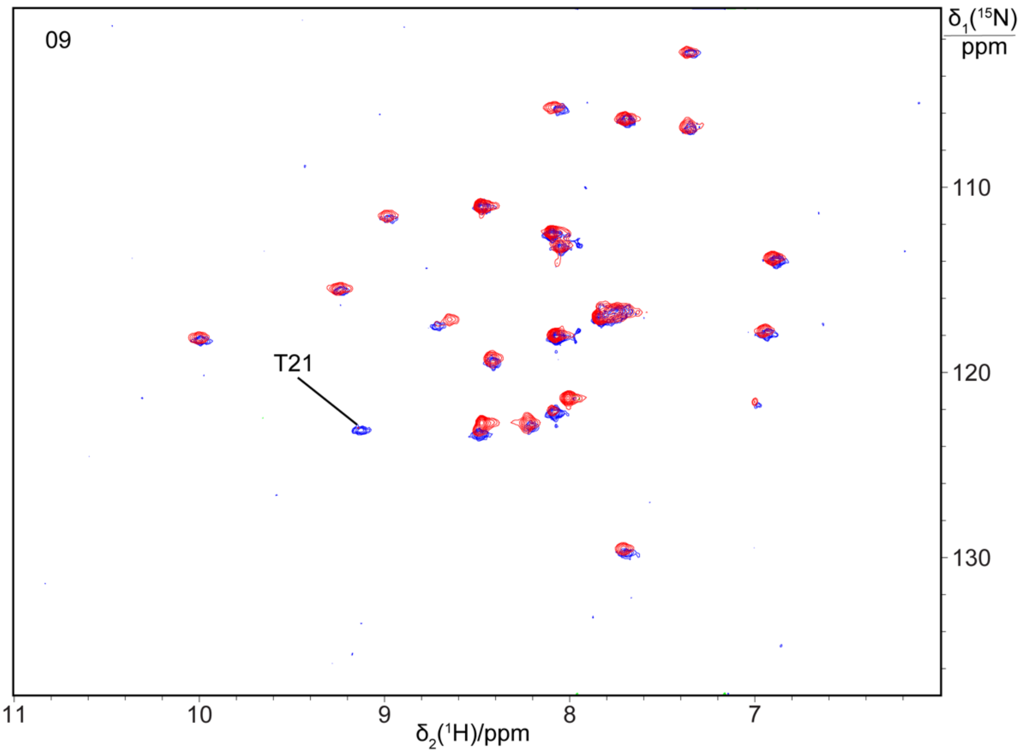
09
δ1(15N)
ppm
110
120
130
T21
11
10
9
8
7
δ2(1H)/ppm

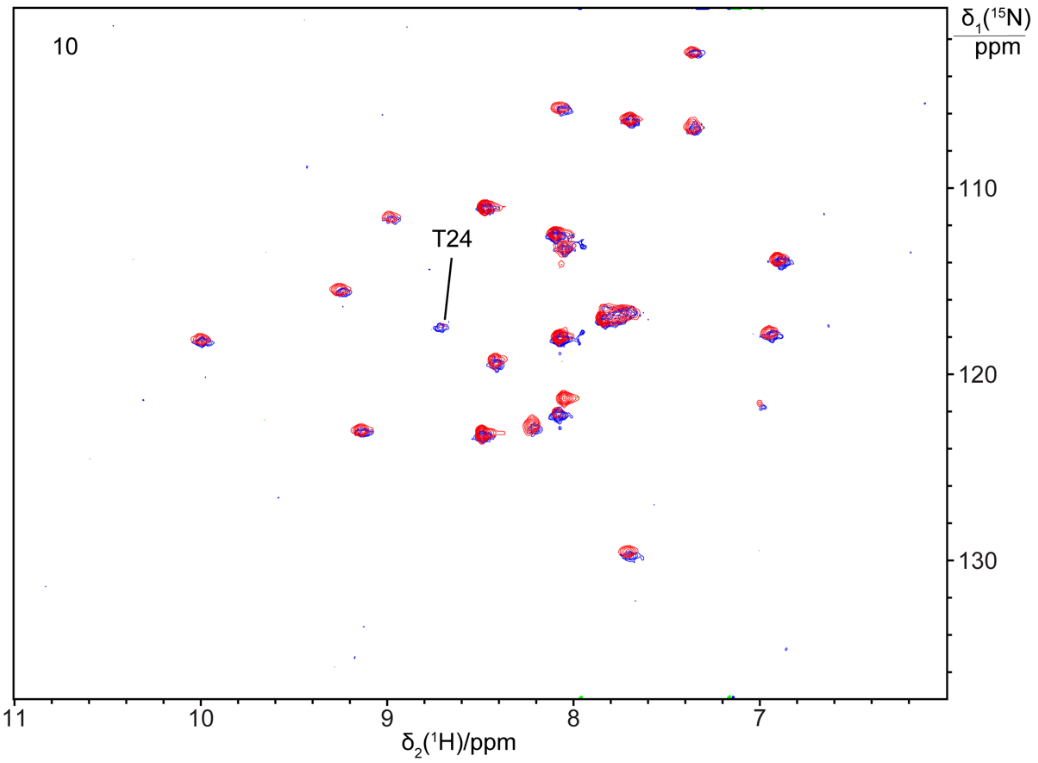
10
δ1(15N)
ppm
110
120
130
T24
11
10
9
8
7
δ2(1H)/ppm

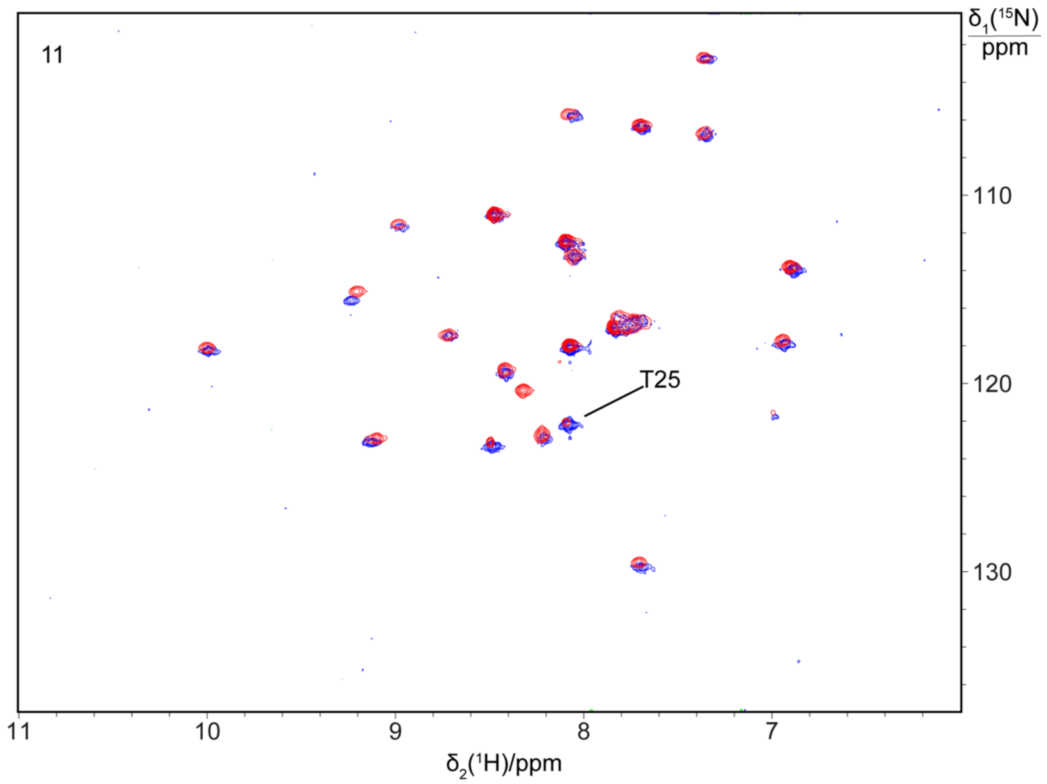
11
T25
δ1(15N)/ppm
110
120
130
11
10
9
8
7
δ2(1H)/ppm

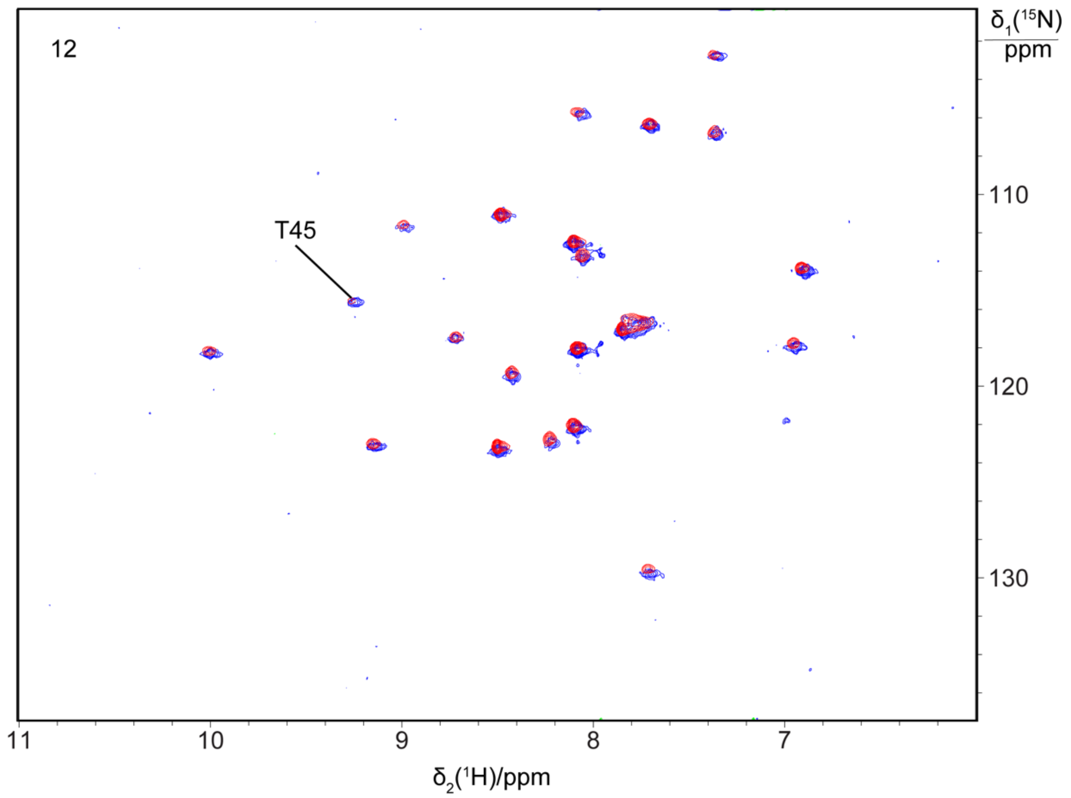
12
T45
δ1(15N)/ppm
110
120
130
11
10
9
8
7
δ2(1H)/ppm

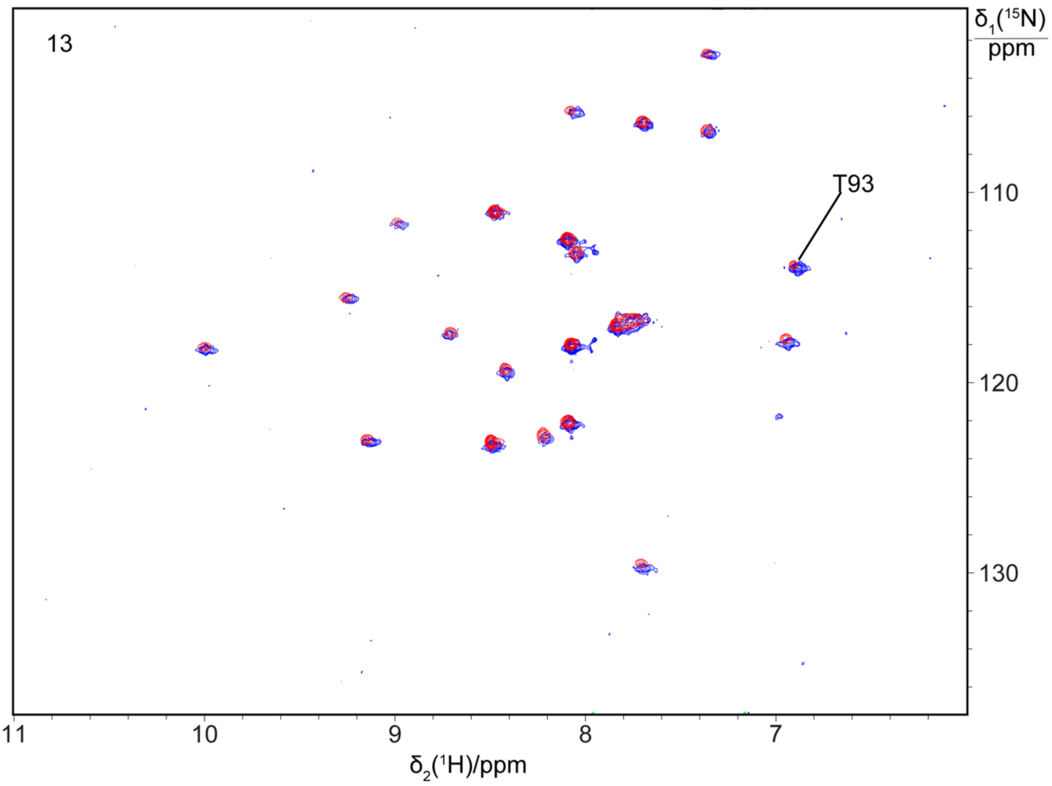
13
T93
δ1(15N)/ppm
110
120
130
11
10
9
8
7
δ2(1H)/ppm

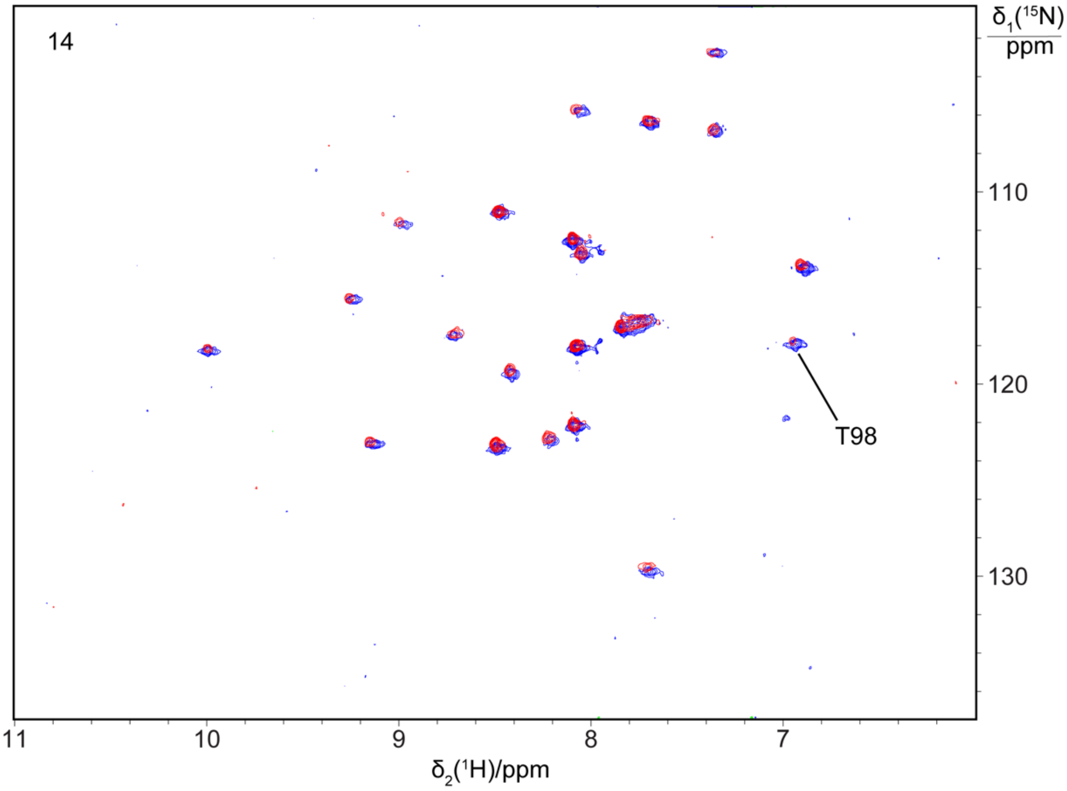
14
T98
δ1(15N)/ppm
110
120
130
11
10
9
8
7
δ2(1H)/ppm

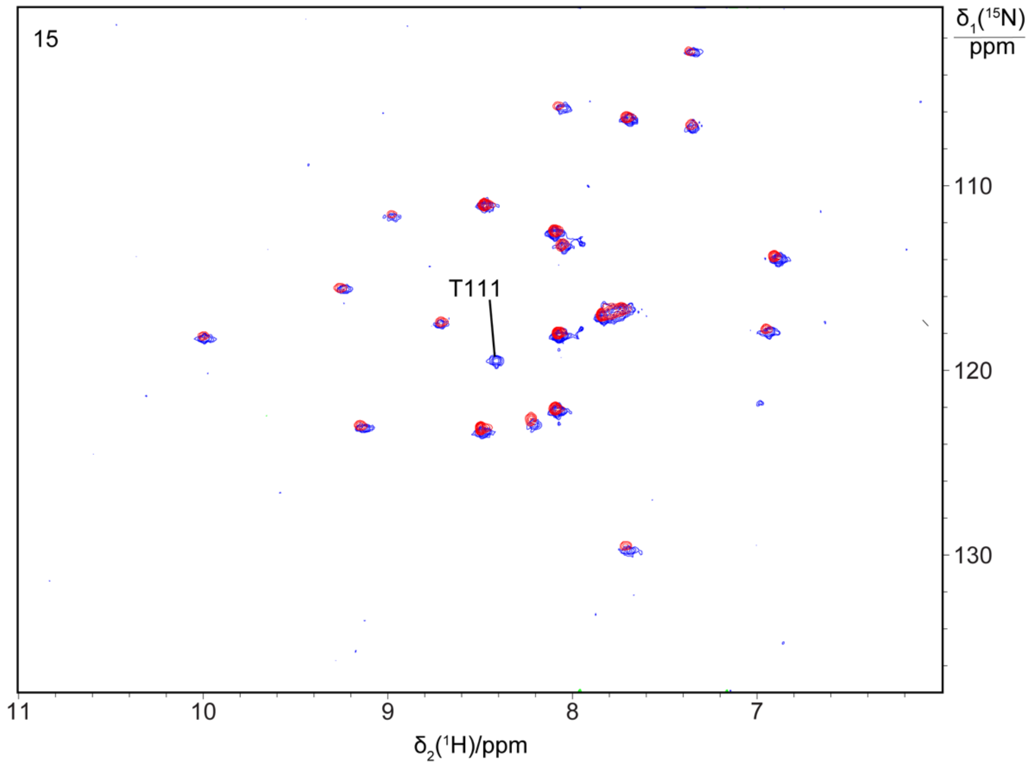
15
T111
δ1(15N)/ppm
110
120
130
11
10
9
8
7
δ2(1H)/ppm

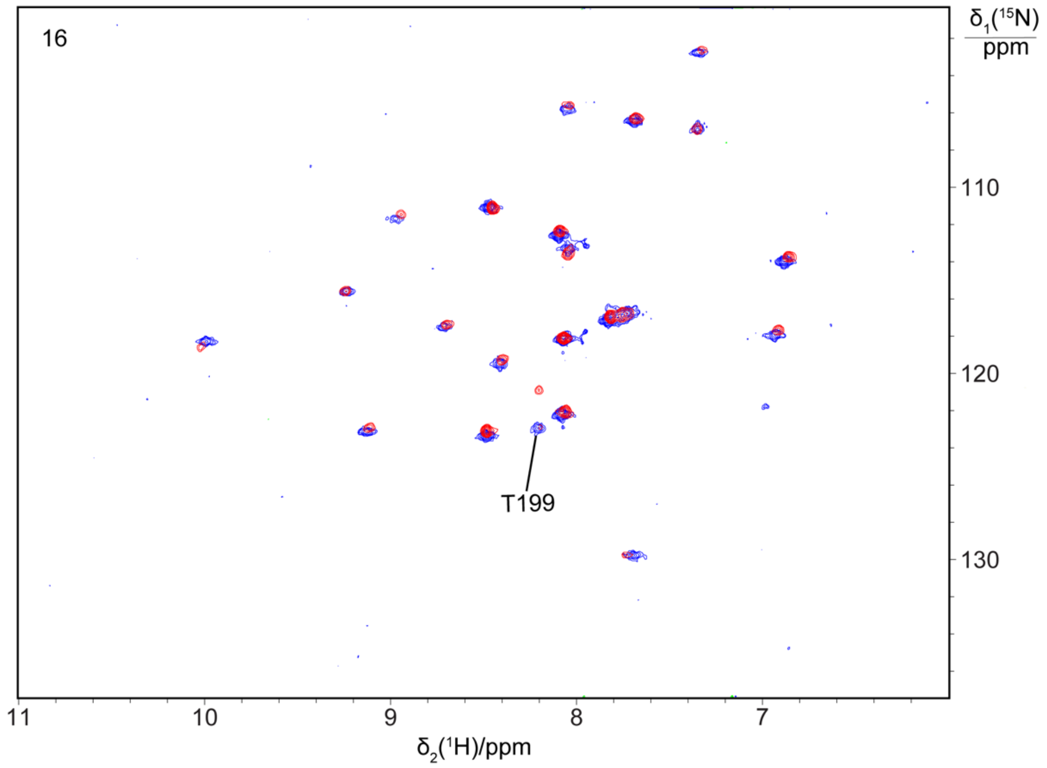
16
T199
δ1(15N)/ppm
110
120
130
11
10
9
8
7
δ2(1H)/ppm

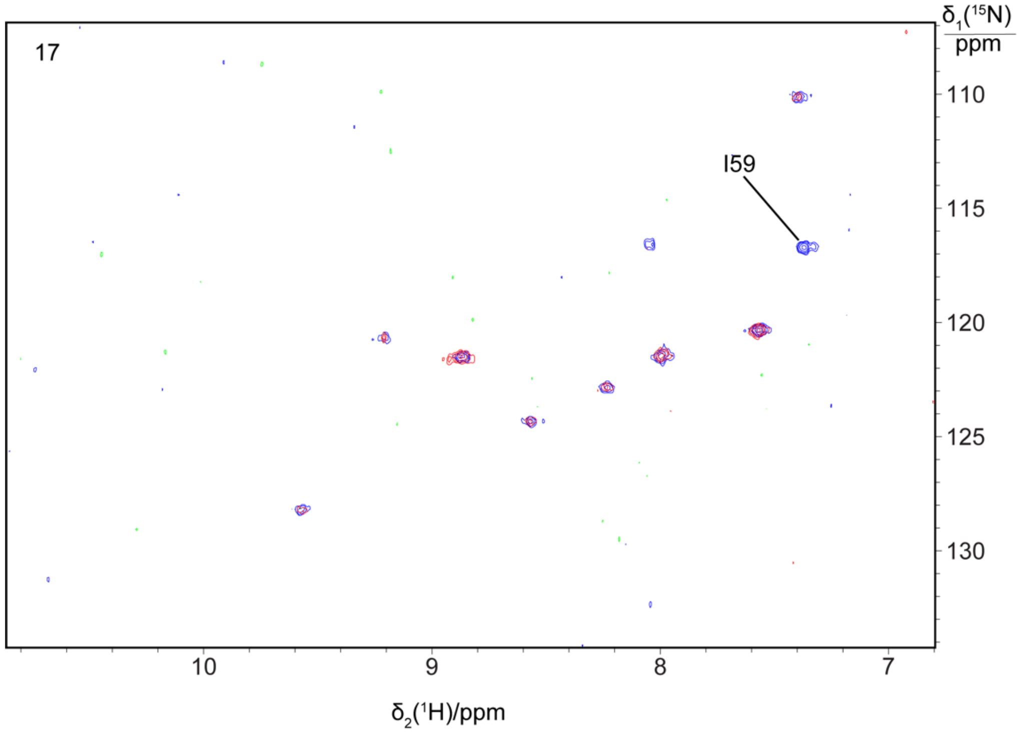
17
I59
δ1(15N)/ppm
110
115
120
125
130
10
9
8
7
δ2(1H)/ppm

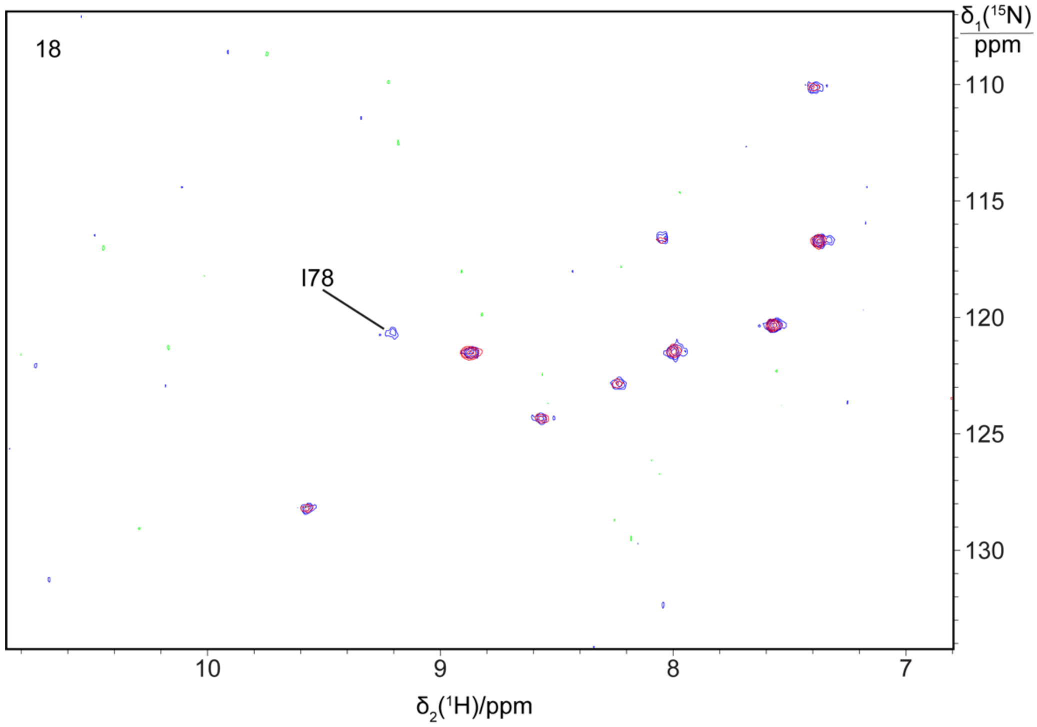
18
I78
δ1(15N)/ppm
110
115
120
125
130
10
9
8
7
δ2(1H)/ppm

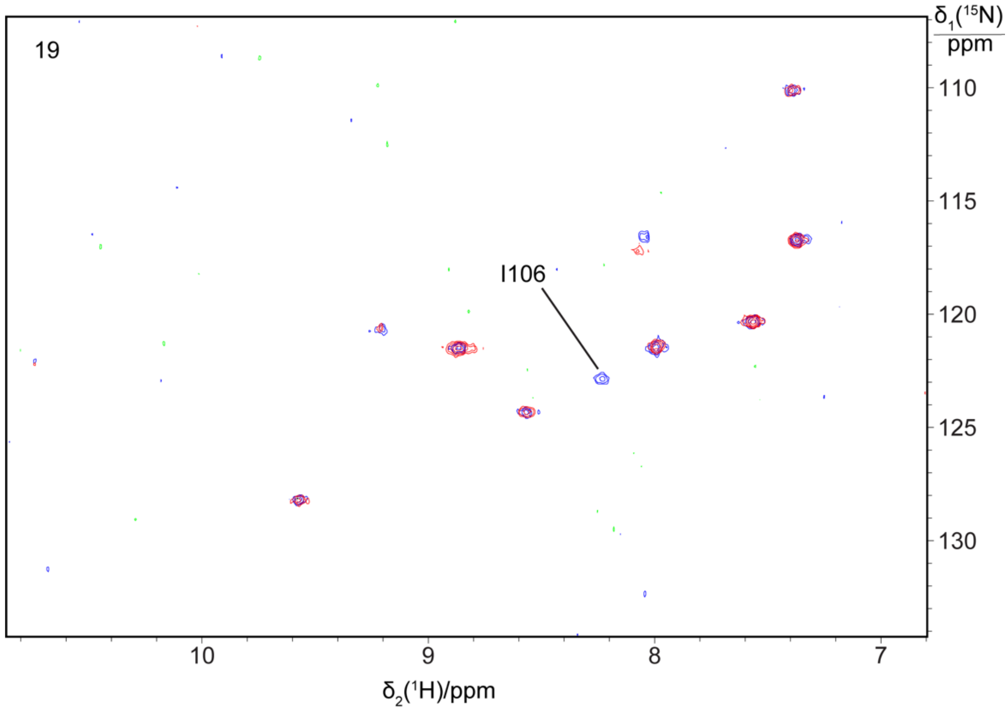
19
I106
δ1(15N) ppm
110
115
120
125
130
10
9
8
7
δ2(1H)/ppm

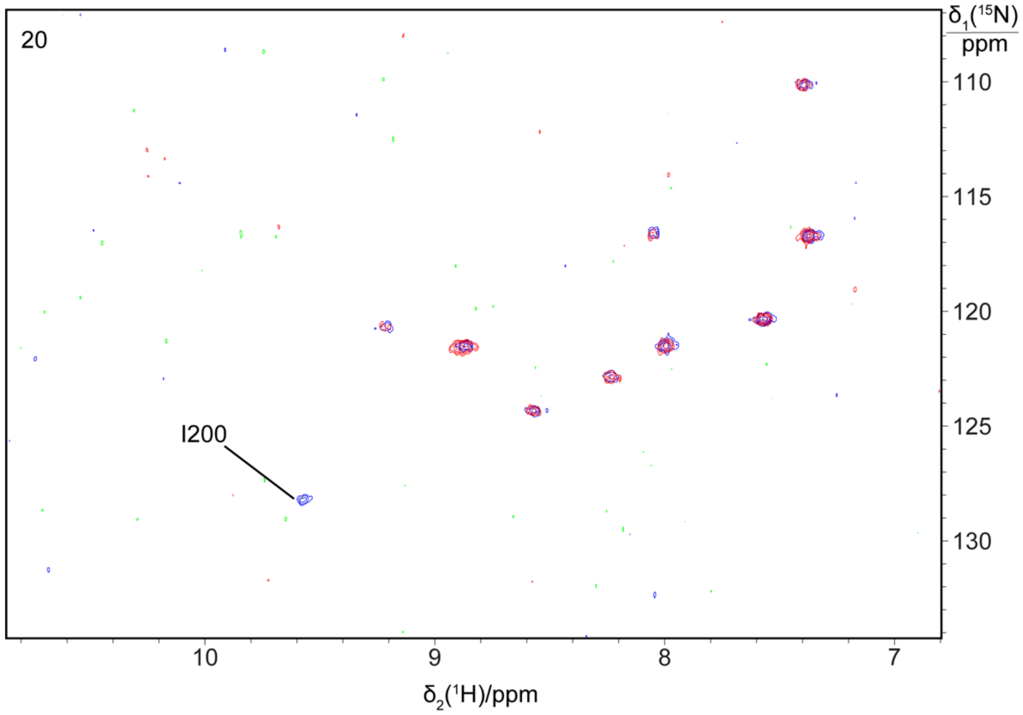
20
I200
δ1(15N) ppm
110
115
120
125
130
10
9
8
7
δ2(1H)/ppm

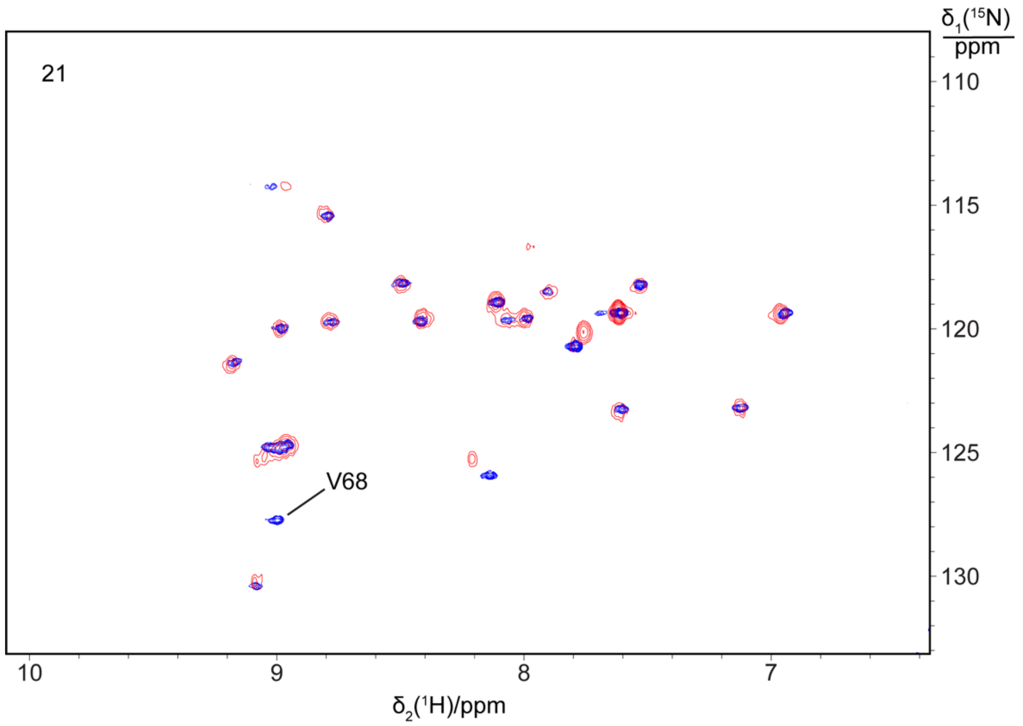

21
V68
δ1(15N)/ppm
110
115
120
125
130
10
9
8
7
δ2(1H)/ppm

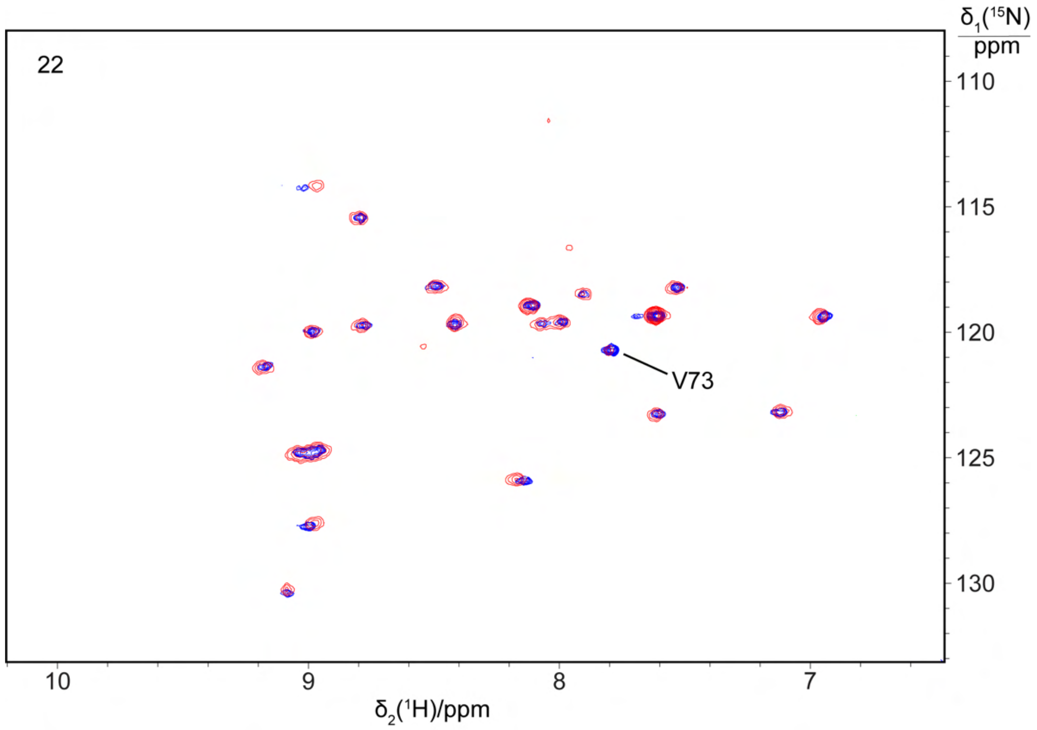

22
V73
δ1(15N)/ppm
110
115
120
125
130
10
9
8
7
δ2(1H)/ppm

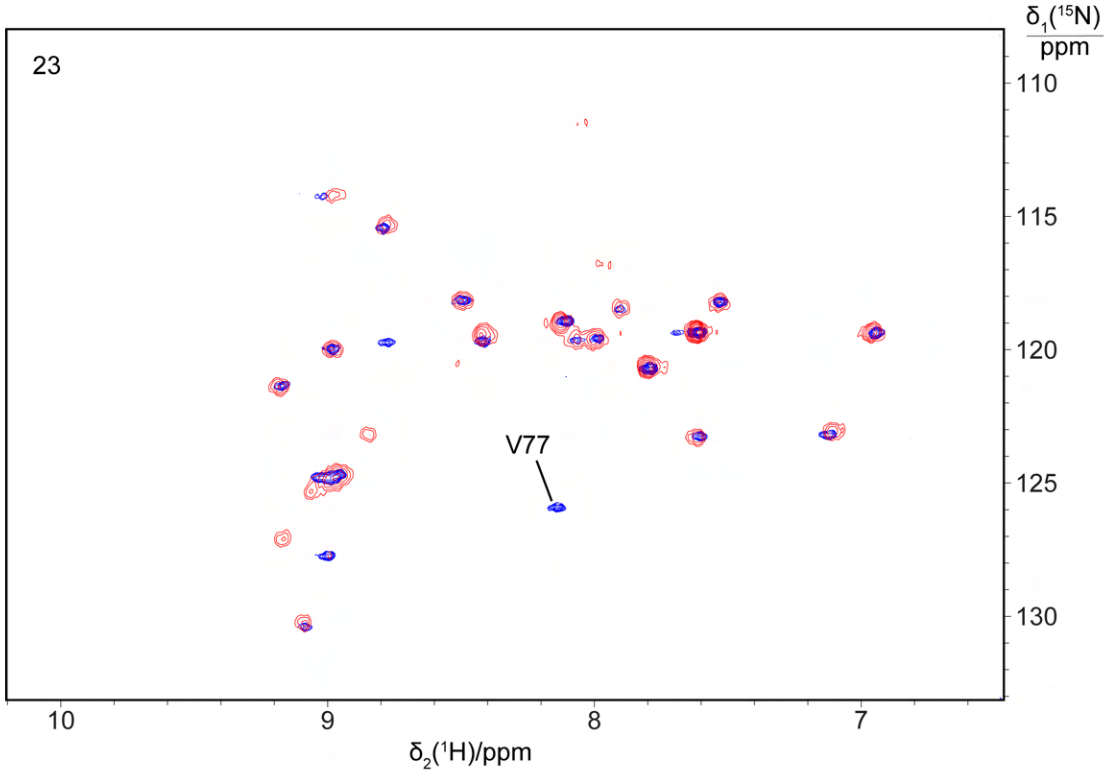
23
V77
δ1(15N)/ppm
110
115
120
125
130
10
9
8
7
δ2(1H)/ppm

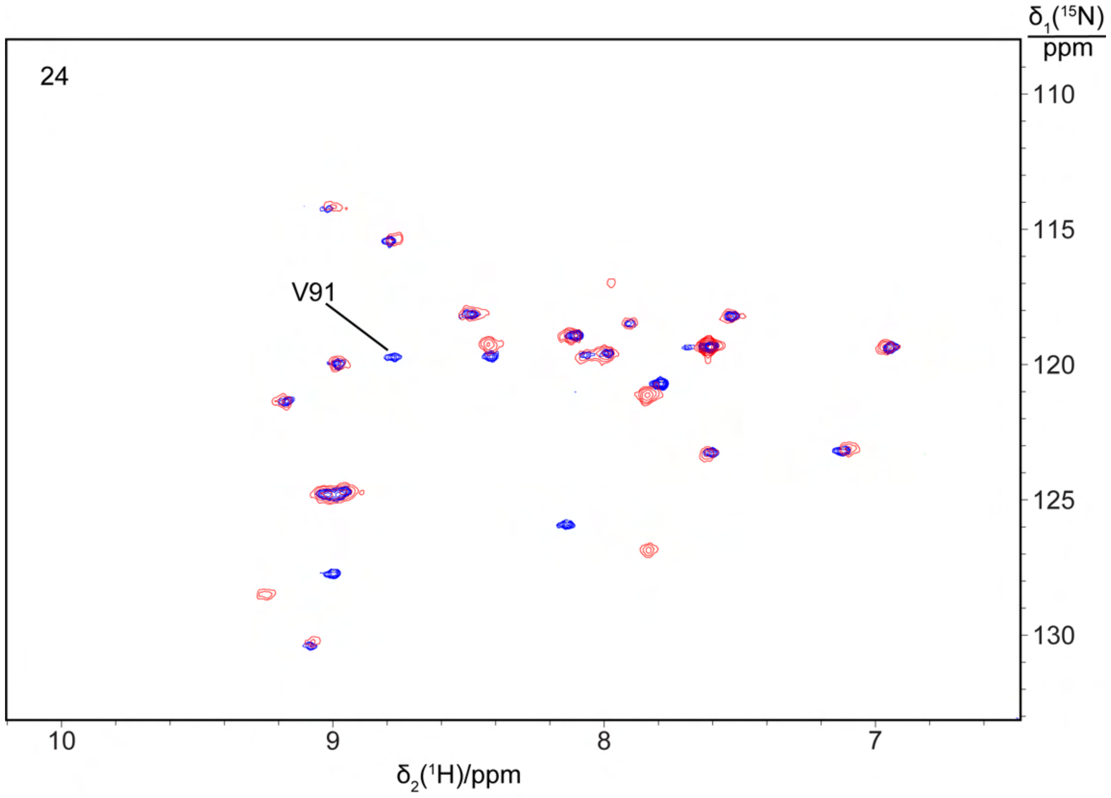
24
V91
δ1(15N)/ppm
110
115
120
125
130
10
9
8
7
δ2(1H)/ppm

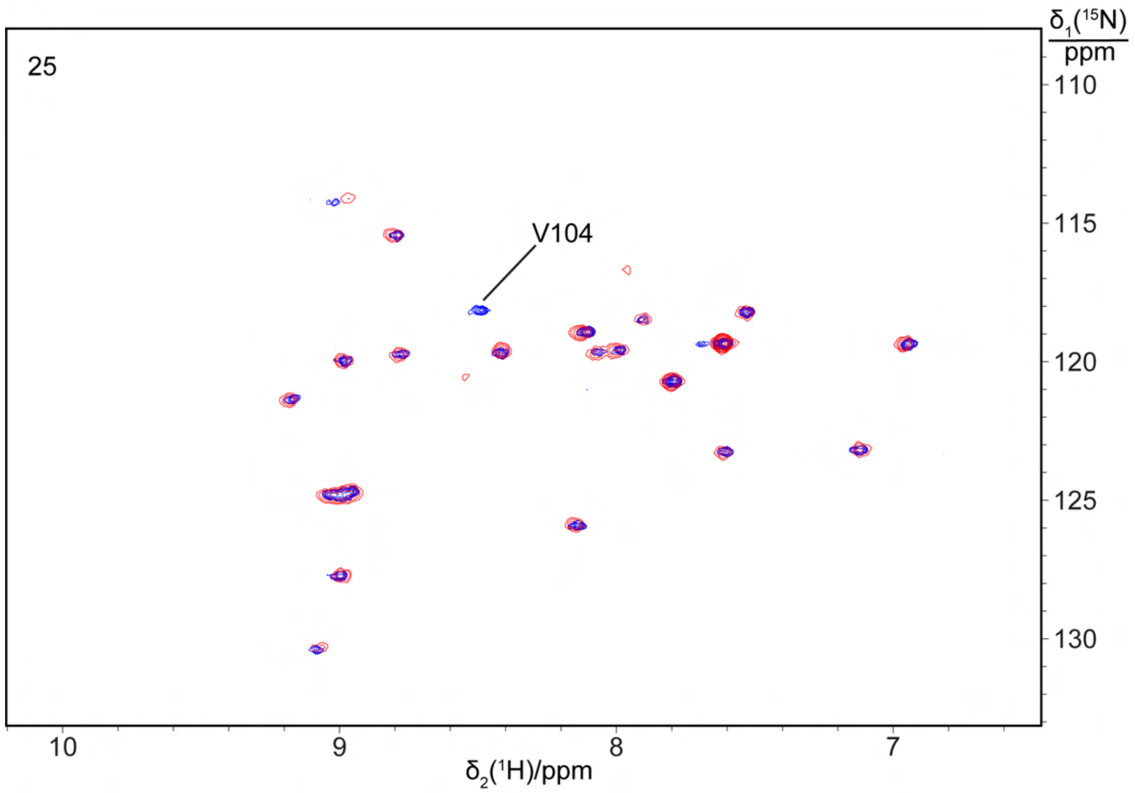
25
V104
δ1(15N)/ppm
110
115
120
125
130
10
9
8
7
δ2(1H)/ppm

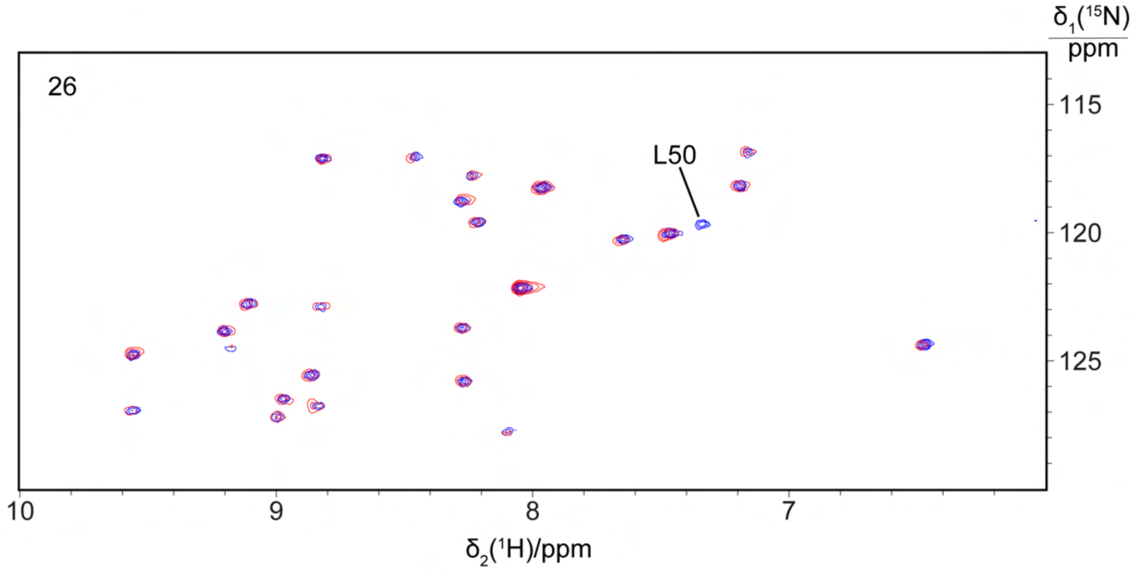
26
L50
δ1(15N)/ppm
115
120
125
10
9
8
7
δ2(1H)/ppm

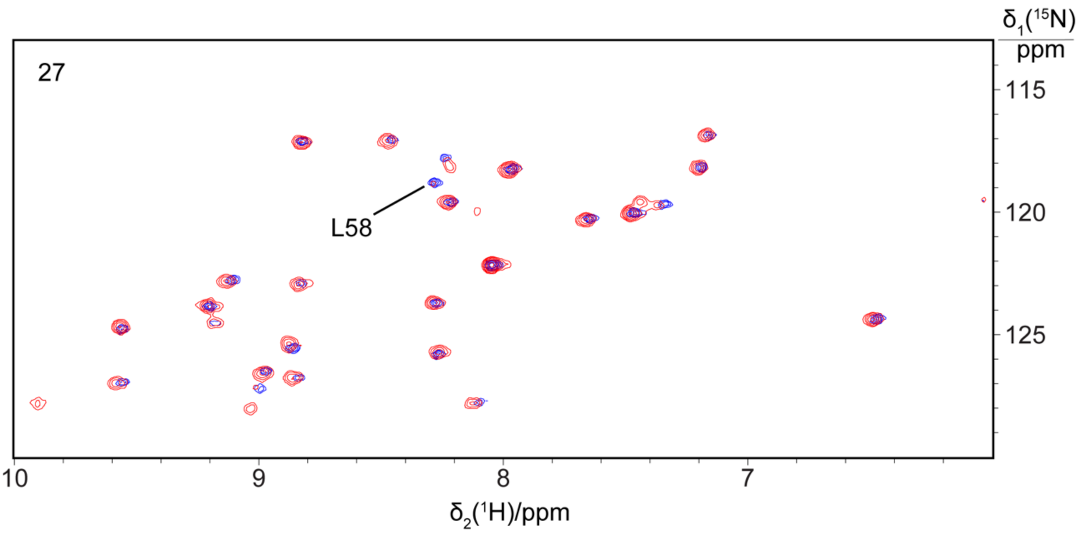
27
L58
δ1(15N)/ppm
115
120
125
10
9
8
7
δ2(1H)/ppm

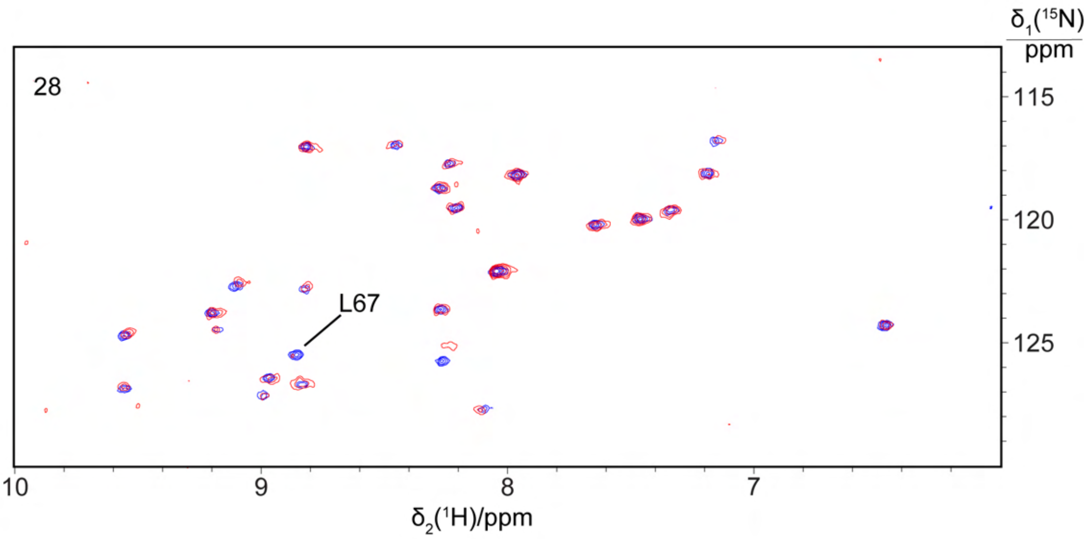
28
L67
δ1(15N)/ppm
115
120
125
10
9
8
7
δ2(1H)/ppm

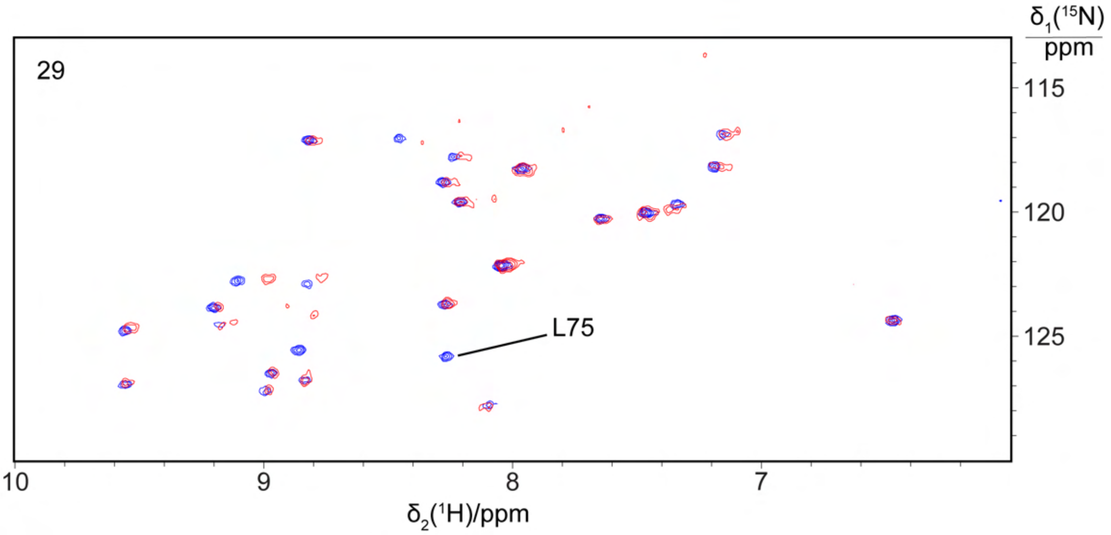
29
L75
δ1(15N)/ppm
115
120
125
10
9
8
7
δ2(1H)/ppm

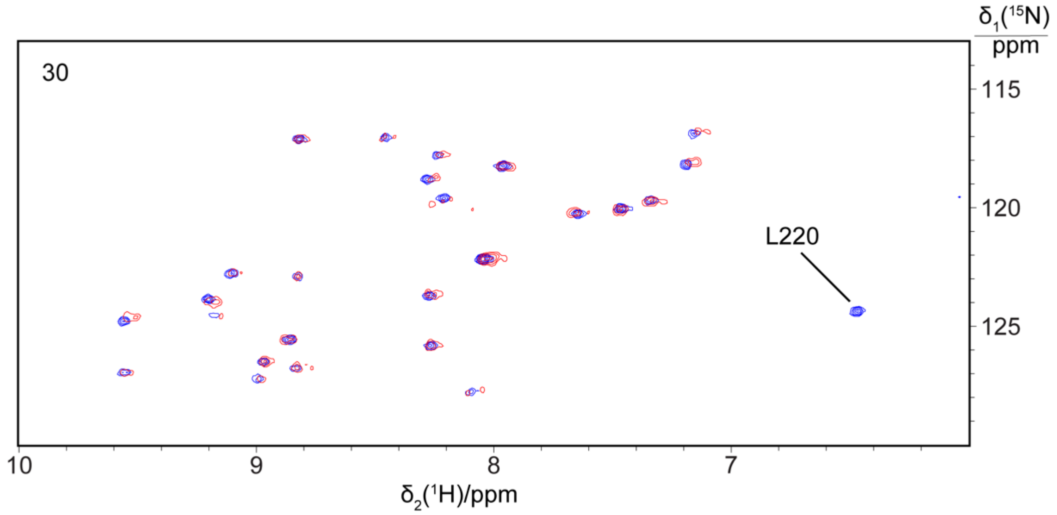
30
L220
δ1(15N)/ppm
115
120
125
10
9
8
7
δ2(1H)/ppm

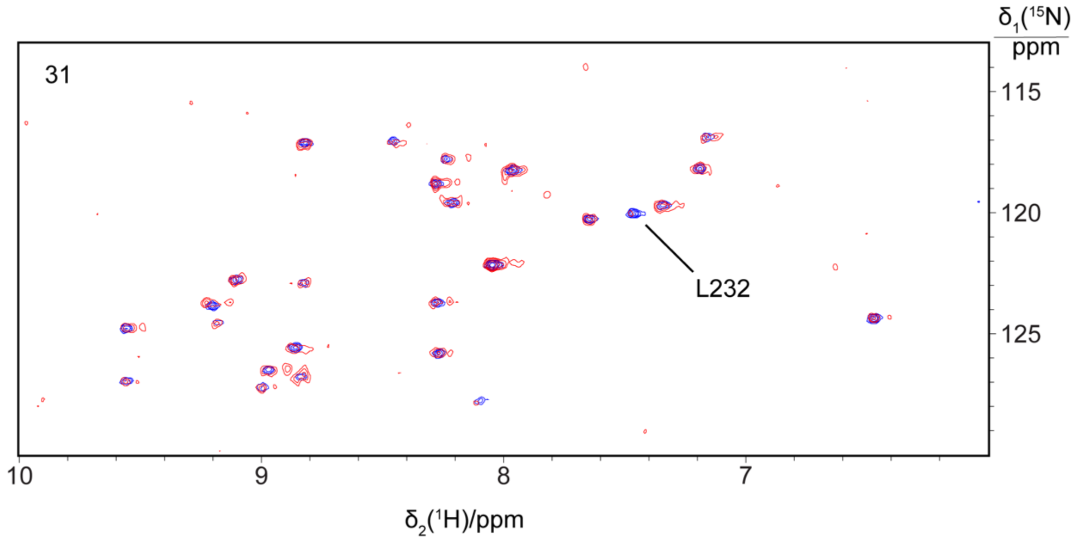
31
L232
δ1(15N)/ppm
115
120
125
10
9
8
7
δ2(1H)/ppm

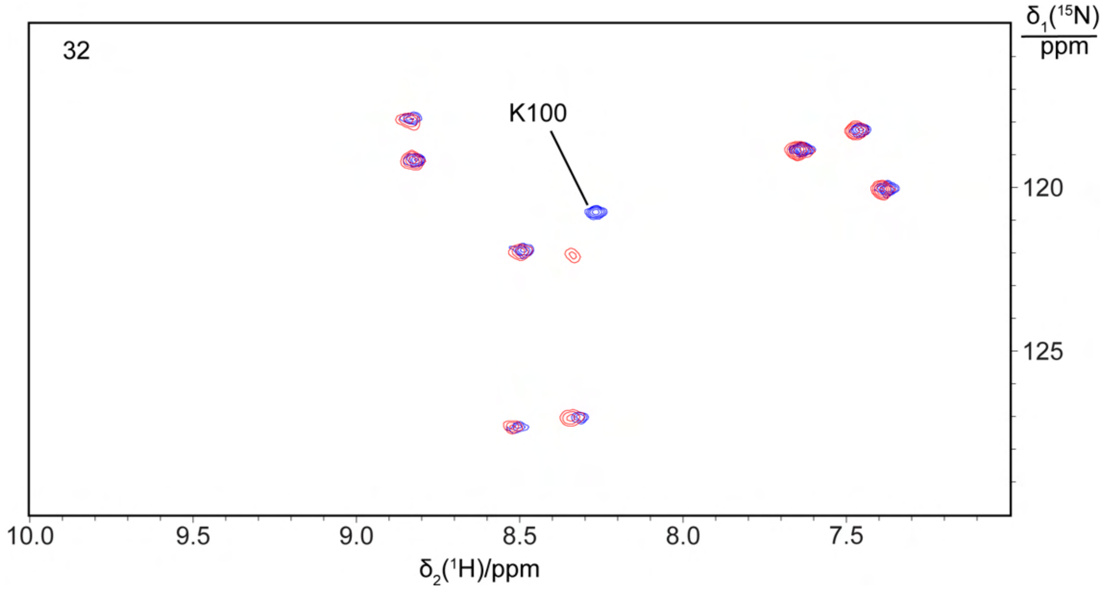
32
K100
δ1(15N)/ppm
120
125
10.0
9.5
9.0
8.5
8.0
7.5
δ2(1H)/ppm

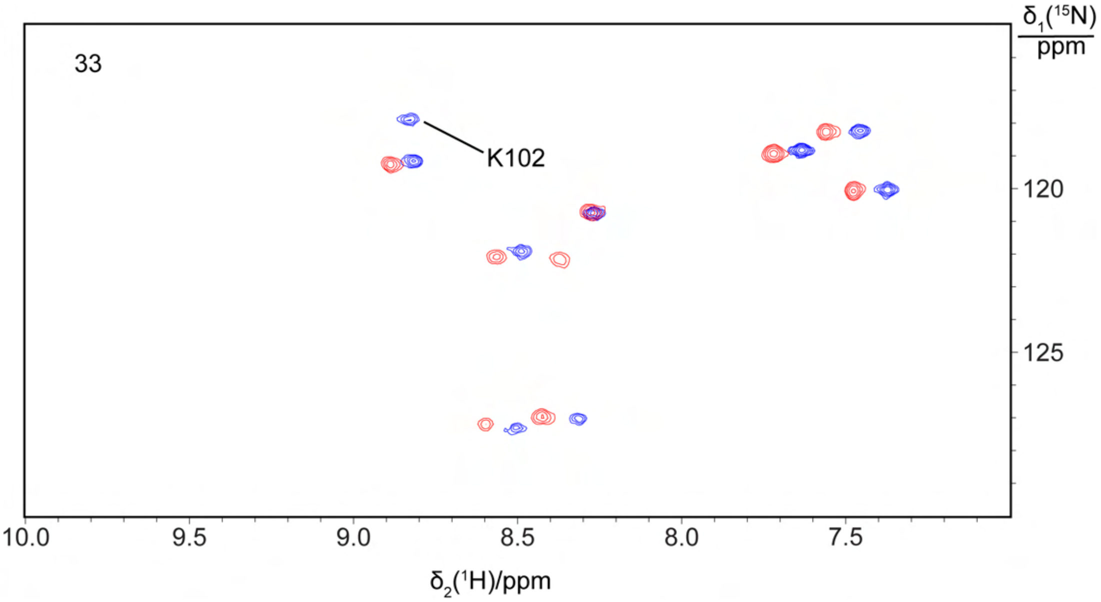
33
K102
δ1(15N)/ppm
120
125
10.0
9.5
9.0
8.5
8.0
7.5
δ2(1H)/ppm

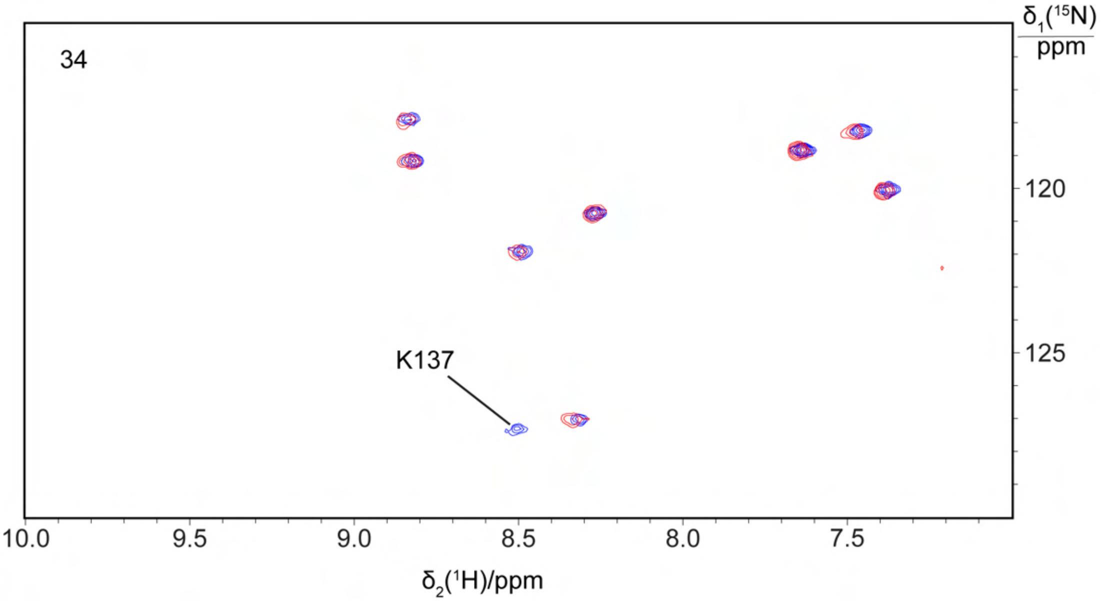
34
K137
δ1(15N)/ppm
120
125
10.0
9.5
9.0
8.5
8.0
7.5
δ2(1H)/ppm

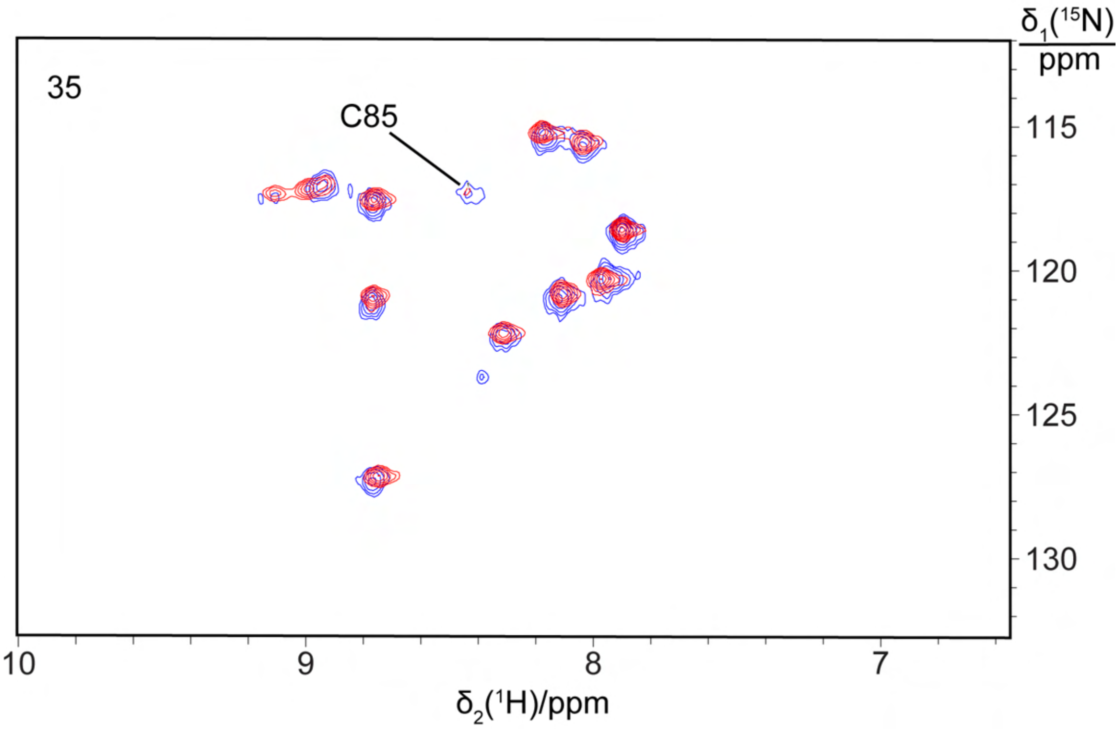

35
C85
δ1(15N)
ppm
115
120
125
130
10
9
8
7
δ2(1H)/ppm

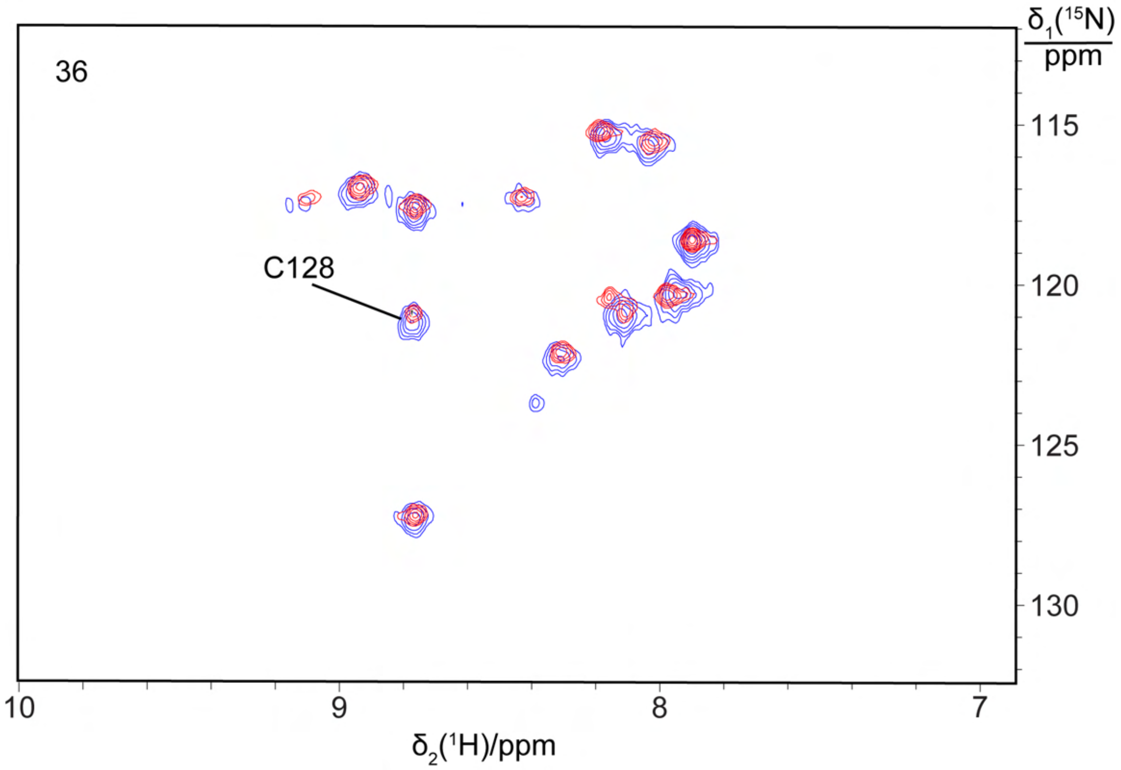

36
C128
δ1(15N)
ppm
115
120
125
130
10
9
8
7
δ2(1H)/ppm

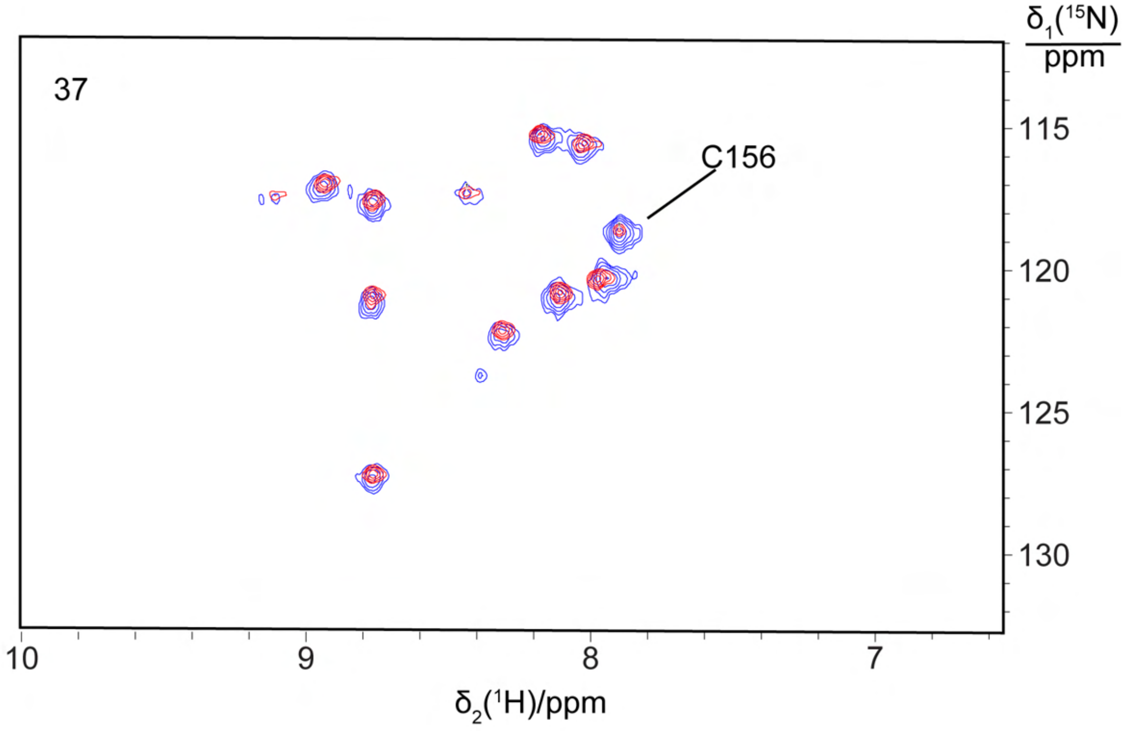
37
C156
δ1(15N)/ppm
115
120
125
130
10
9
8
7
δ2(1H)/ppm

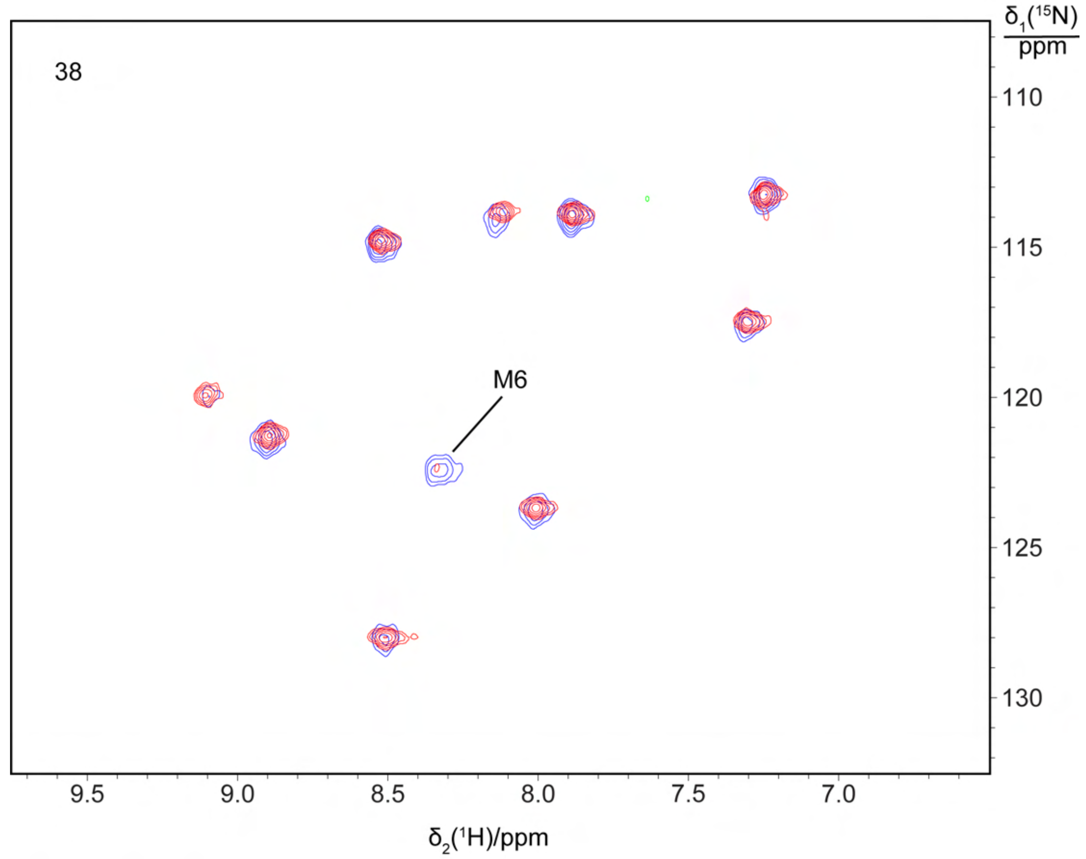
38
M6
δ1(15N)/ppm
110
115
120
125
130
9.5
9.0
8.5
8.0
7.5
7.0
δ2(1H)/ppm

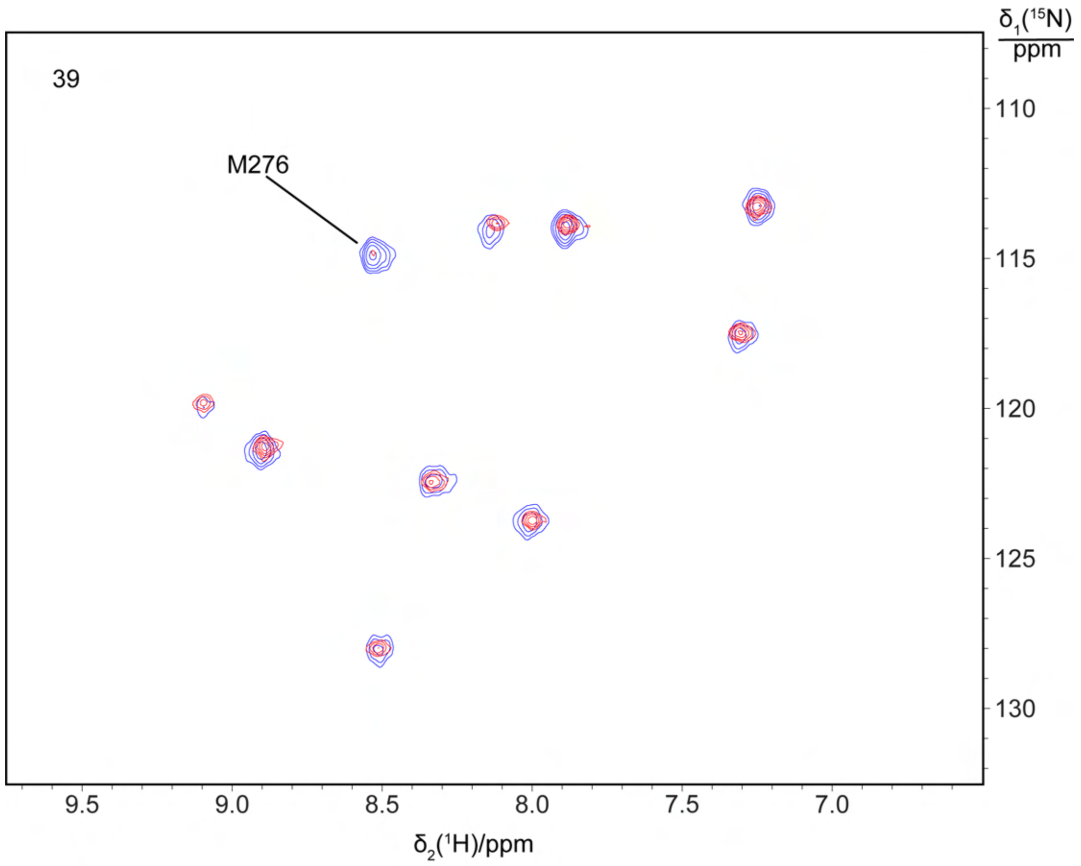

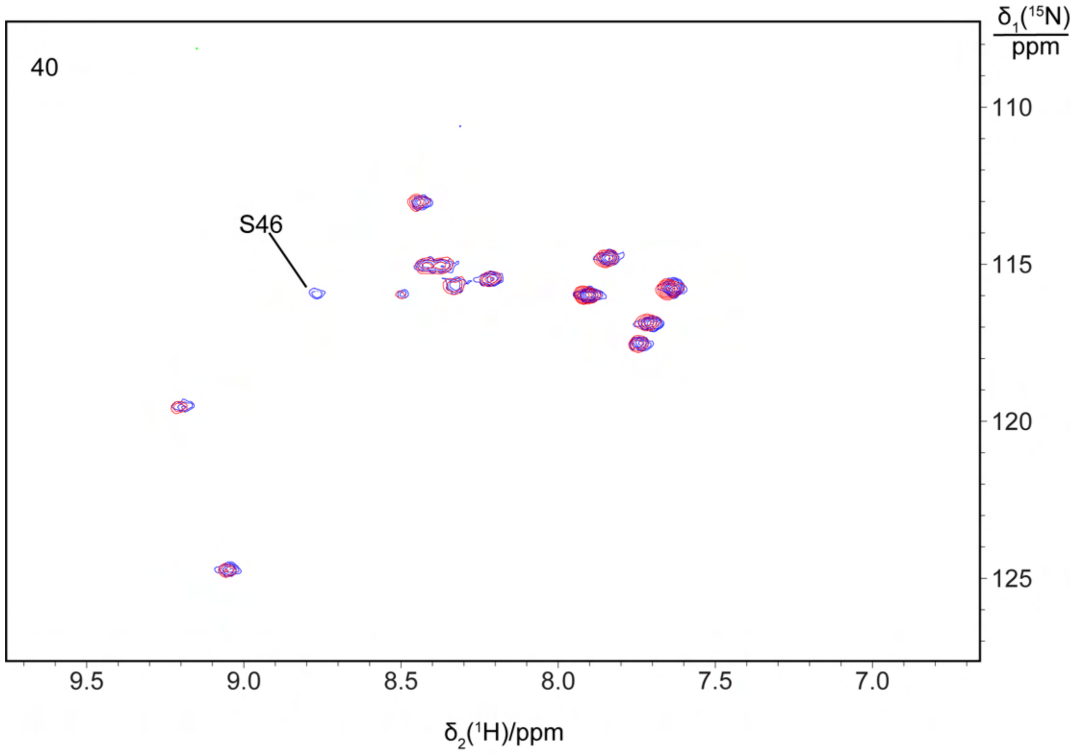

**Supplementary Figure 4.** [$^{15}N$,$^1H$]-HSQC spectra recorded of selectively $^{15}N$-labelled samples of M$^{pro}$ R298A and mutants enabling specific resonance assignments. The spectra show superimpositions of the selectively labelled samples (blue) and the corresponding samples with a site-specific mutation (red). The panel numbers refer to Table S2, which reports the amino acid type labelled with $^{15}N$ and the mutation site. The residue types installed by the mutations are specified in Table S1. The sequence-specific resonance assignment made with each mutation is indicated in the spectra.

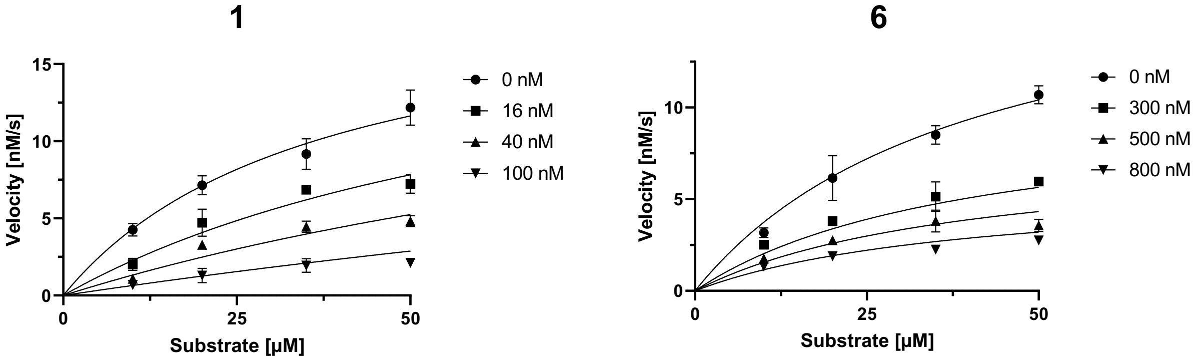

**Supplementary Figure 5. Michaelis-Menten saturation curves of SARS-CoV-2 $M^{pro}$ in the presence of range of concentrations of cyclic peptide inhibitors 1 and 6.**

**a**

**b**

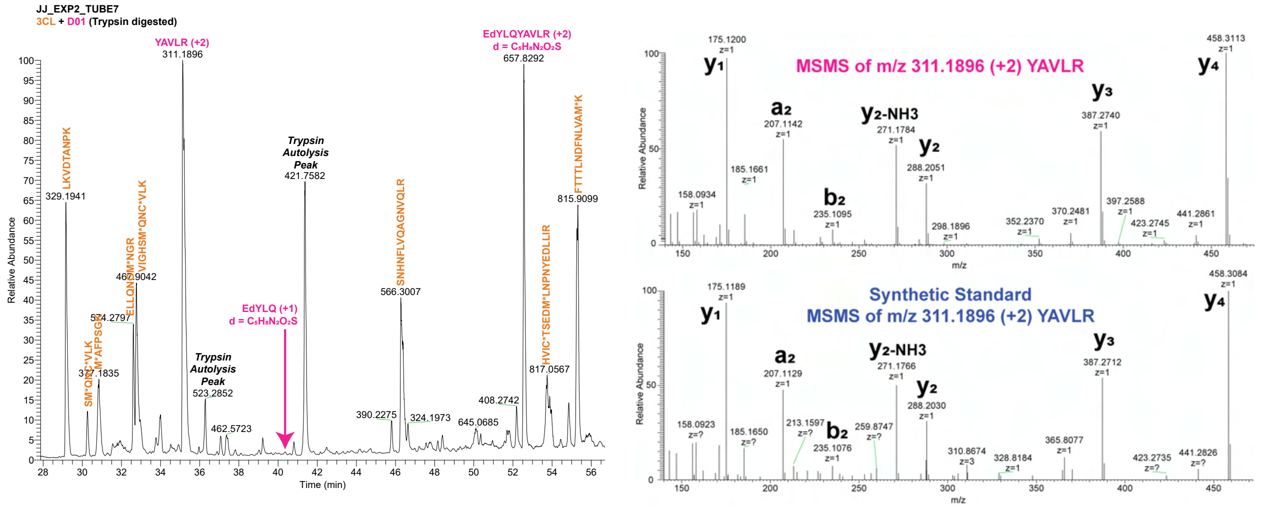

**c**

| Peptide | IC50 (μM) |
|---|---|
| 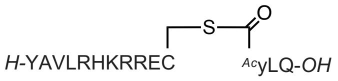 | 23.2 ± 5.0 |

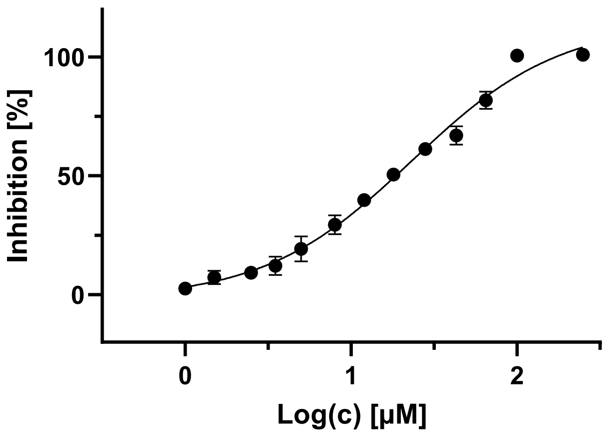

**Supplementary Figure 6. a LC-MS characterization of cleavage site of 1.** Peptide **1** (5 μM) was incubated with SARS-CoV-2 M$^{pro}$ (2.5 μM) in aqueous buffer (20 mM Tris, 100 mM NaCl, 1 mM DTT, 1mM EDTA, pH 7.5) for 1 h at 37 °C. After trypsin digestion the sample was analyzed by LC/MS-MS analysis. Major masses identified to trypsin fragments were consistent with cleavage by M$^{pro}$ between Q3 and Y4 (YAVLR and yLQ). **b MS/MS analysis of peak at m/z 311.1896**. The fragmentation pattern corresponded with the peptide sequence YAVLR, consistent with proteolytic cleavage by M$^{pro}$ C-terminal to Gln3 (upper panel). Validation of HCD MSMS fragmentation pattern using a synthetic standard of the linear peptide that would be generated upon cleavage of cyclic peptide **1** C-terminal to Gln3 (lower panel). **c Inhibitory activity of authentic standard of cleaved linear form of 1 against**

**SARS-CoV-2 M$^{pro}$.** Structure of linear peptide and associated $IC_{50}$ value against SARS-CoV-2 M$^{pro}$ shown in the table.

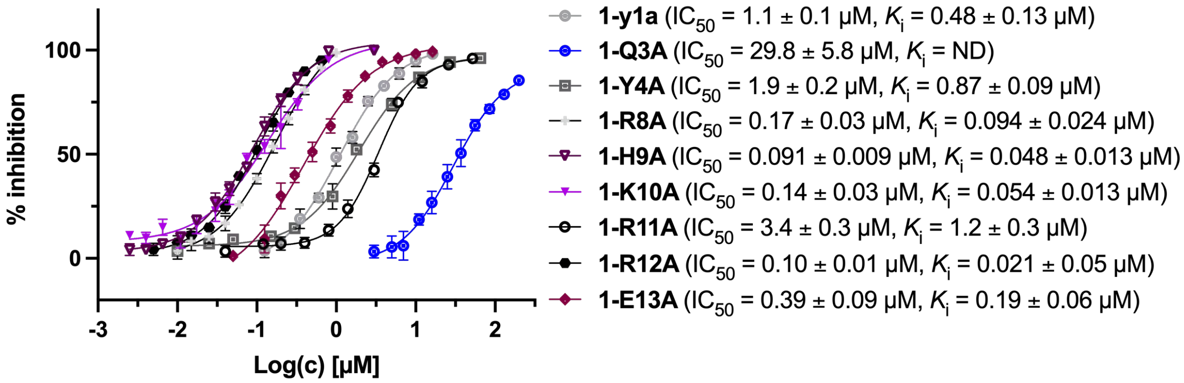

**Supplementary Figure 7. Dose response curves for single site alanine mutants of lead cyclic peptide 1 against SARS-CoV-2 M$^{pro}$.**

| Peptide | Structure | $IC_{50}$ (μM) |
|---|---|---|
| **1-Sec** | 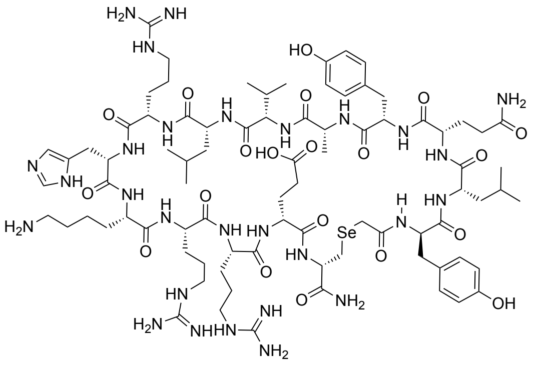 | 0.056 ± 0.005 |

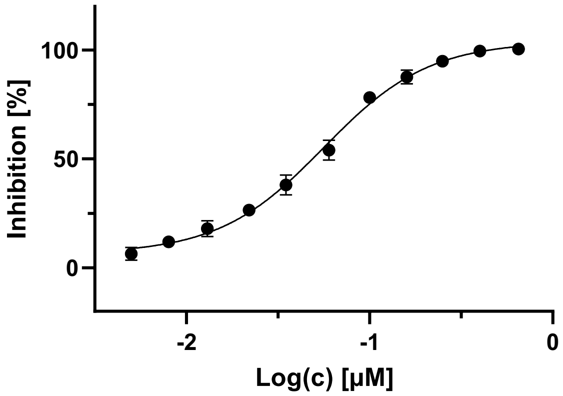

**Supplementary Figure 8. Structure and inhibitory activity of selenoether analogue of 1 (Sec-1) against SARS-CoV-2 M$^{pro}$.**

**Supplementary Table 1. Data collection and refinement statistics.** For the low resolution SARS-CoV-2 $M^{pro}$:**1** complex we provide data collection statistics to demonstrate the SARS-CoV-2 $M^{pro}$:**Se-1** crystal has similar space group/unit cell dimensions. The quality of the electron density in the SARS-CoV-2 $M^{pro}$:**1** complex was not sufficient to build a high-quality model or resolve the binding of the ligand.

| | SARS-CoV-2 $M^{pro}$-**Se-1** | SARS-CoV-2 $M^{pro}$ **-1** |
|---|---|---|
| **PDB ID** | 7RNW | N/A |
| **Data collection** | | |
| Space group | P 1 $2_1$ 1 | P1 $2_1$ 1 |
| Cell dimensions | | |
| *a, b, c (Å)* | 48.4 201.7 60.5 | 47.6, 190.1, 59.15 |
| a, b, g (°) | 90.0 113.5 90.0 | 90, 113.64, 90 |
| Resolution (*Å)* | 33.62-2.25 (2.32-2.25)* | 39.66-3.45 (3.78-3.45)* |
| $R_{merge}$ | 0.267 (2.587) | 0.658 (2.249) |
| $R_{pim}$ | 0.168 (1.663) | 0.370 (1.297) |
| $I/\sigma I$ | 5.7 (0.9) | 4.4 (1.4) |
| $CC_{1/2}$ | 0.99 (0.329) | 0.933 (0.315) |
| Completeness (%) | 99.8 (99.5) | 99.5 (99.3) |
| Redundancy | 6.9 (6.3) | 7.2 (7.0) |
| **Refinement** | | |
| Resolution (*Å)* | 33.62-2.25 (2.32-2.25) | |
| No. reflections | 50104 (5021) | |
| $R_{work}/R_{free}$ | 0.2182/0.2757 | |
| No. atoms | 9897 | |
| Protein | 9834 | |
| Ligand/ion | 16 | |
| Water | 47 | |
| B-factors (overall) | 38.75 | |
| Protein | 38.81 | |
| Ligand/ion | 42.08 | |
| Water | 24.09 | |
| R.m.s. deviations | | |
| Bond lengths (*Å)* | 0.014 | |
| Bond angles (*Å)* | 1.84 | |

*Statistics for the highest resolution shell are shown in parentheses

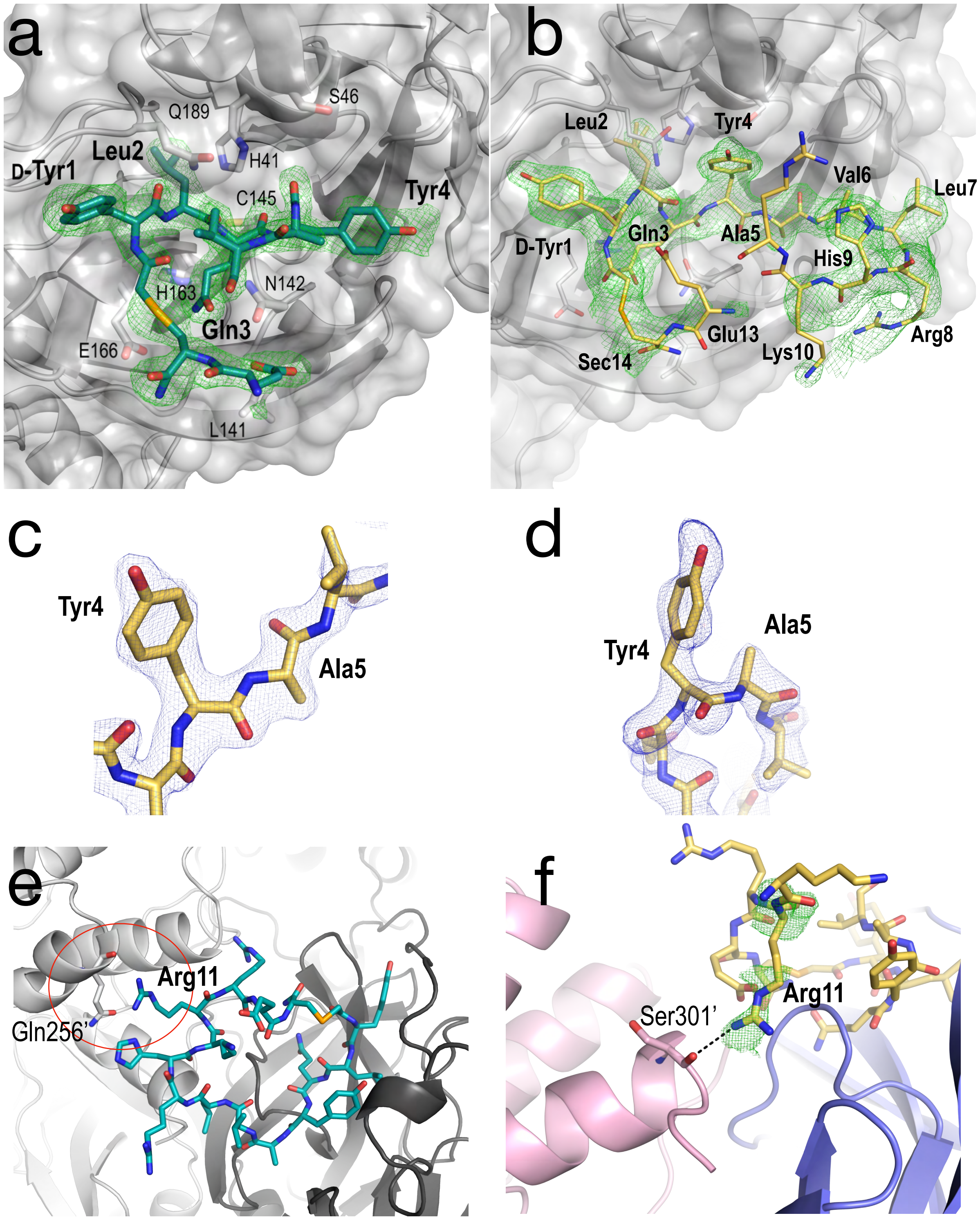

**Supplementary Figure 9. Structural analysis of the complex between selenoether analogue of 1 (Se-1) and SARS-CoV-2 M$^{pro}$. a** Comparison between the binding mode of the peptide in chains A, C, D and **b** chain B shown as sticks with 3.5 s Polder omit electron density map. While the Tyr4-Ala5 *cis*-peptide bond causes a tight turn in the peptide in chains A, C, D, the Tyr4-Ala5 *trans*-peptide bond in chains

B results in the main chain running through the site otherwise occupied by Tyr4 in the other chains. **c** and **d** show 3.5 s Polder omit electron density maps of the Tyr4-Ala5 *trans*-peptide bond in chain B and the Tyr4-Ala5 *cis*-peptide bond in chain C. **e** Molecular dynamics simulations reveal the peptide can transiently interact with the neighbouring chain in the dimer *via* Arg11. **f** Arg11 can be modelled into weak electron density consistent with a transient interaction with the carbonyl carbon of Ser301 in the neighbouring monomer.

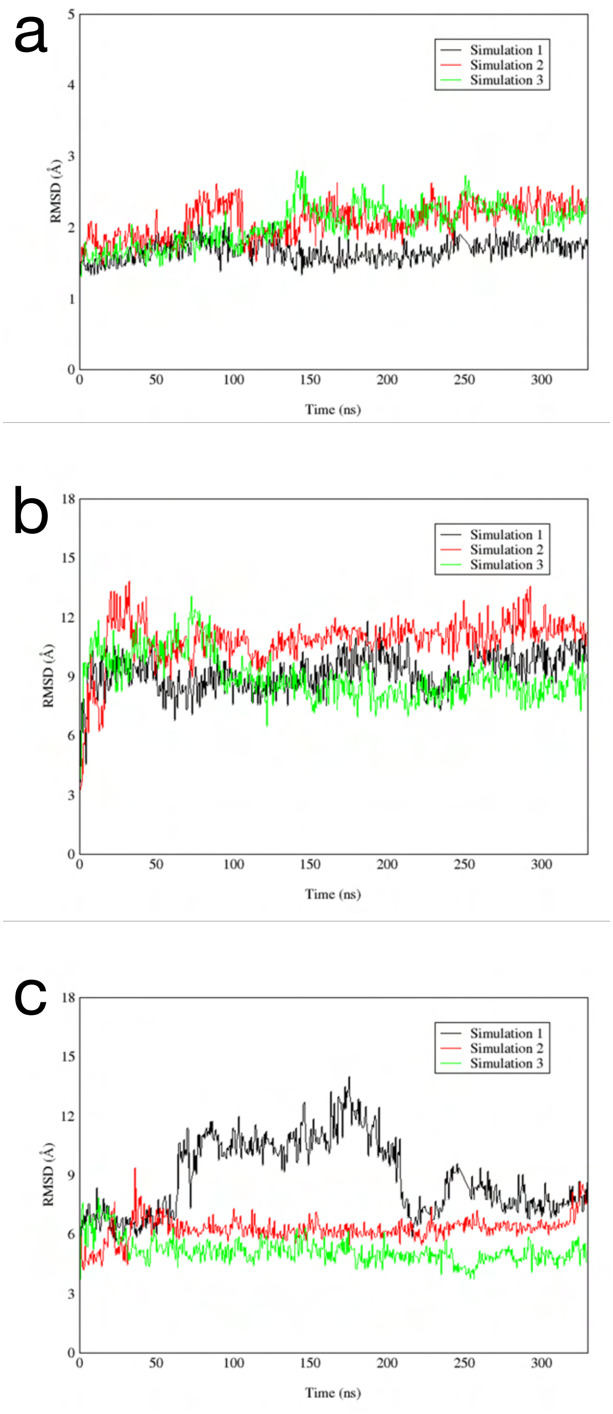

**Supplementary Figure 10. RMSD plots of MD simulations.** For chain A, the peptide Gln3 and Leu2 (P1 and P2 respectively) groups were oriented towards the S1 and S2 subcavities, respectively. As a negative control, for chain B, the peptide was modelled in the opposite orientation where the S2 subcavity was occupied by Tyr4 group. RMSD plots were obtained from the three independent 330 ns simulations. **a** Protein backbone RMSD; **b** RMSD of ligand (bound to chain A) fit to the protein; **c** RMSD of ligand (bound to chain B) fit to protein. The canonical model (Gln3-Leu2 = P1 & P2) was more stable than the alternative reverse model.

| Peptide | Structure | $IC_{50}$ (μM) |
|---|---|---|
| **pen-1** | 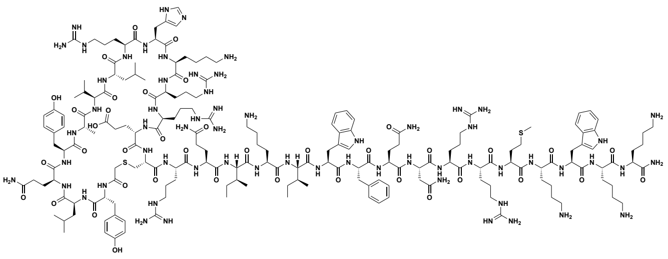 | 0.049 ± 0.009 |
| **pen-6** | 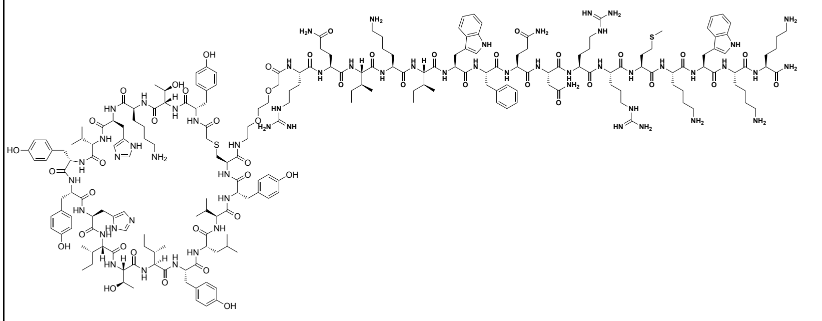 | 3.1 ± 0.2 |

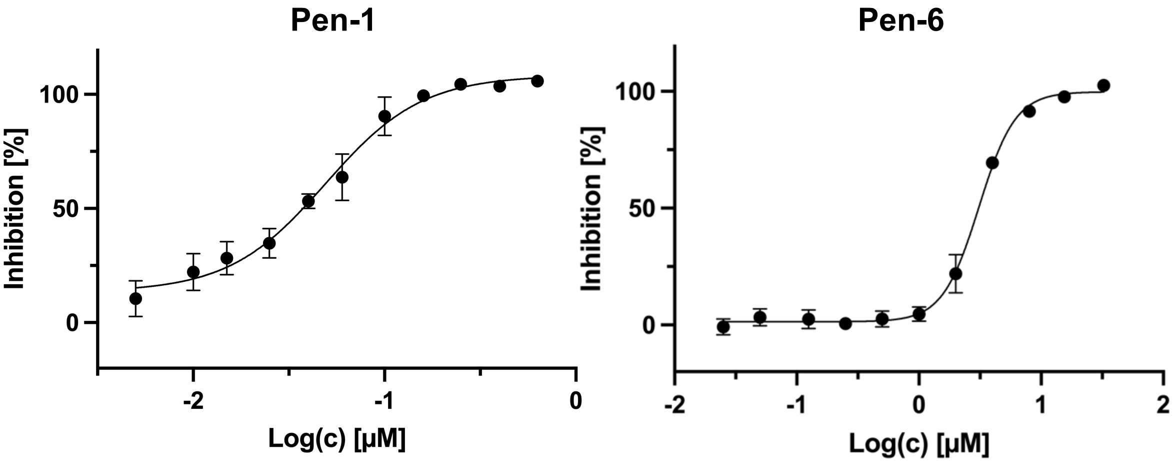

**Supplementary Figure 11. Structures and inhibitory activities of penetratin conjugates of 1 and 6 (pen-1 and pen-6) against SARS-CoV-2 M$^{pro}$.**

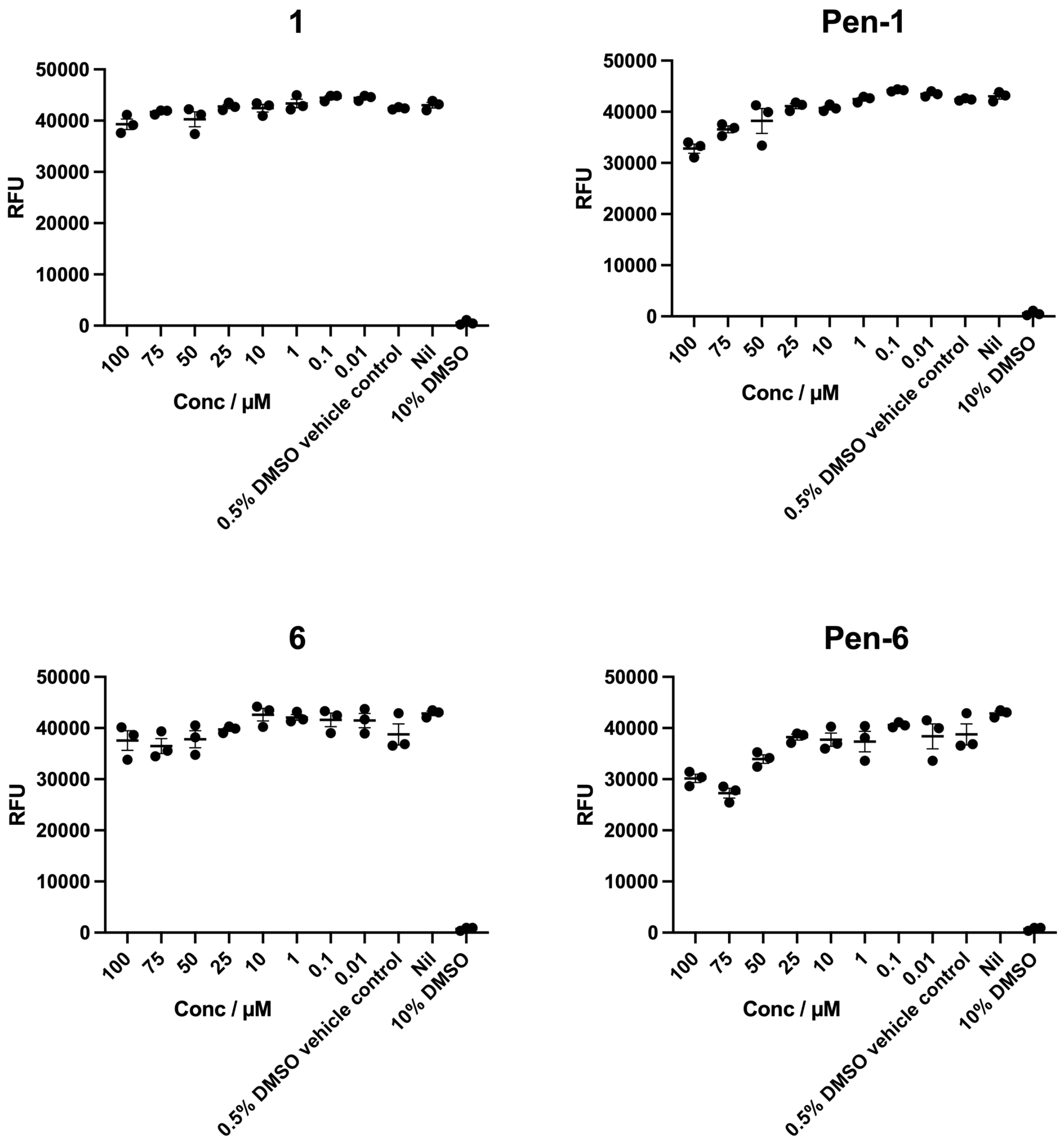

**Supplementary Figure 12. Cytotoxicity of 1, pen-1, 6 and pen-6 against HEK 293-ACE-2-TMPRSS2 cells.** No cytotoxicity was observed for **1** and **6** whilst **pen-1** and **pen-6** resulted in minimal cytotoxicity over 50 µM.

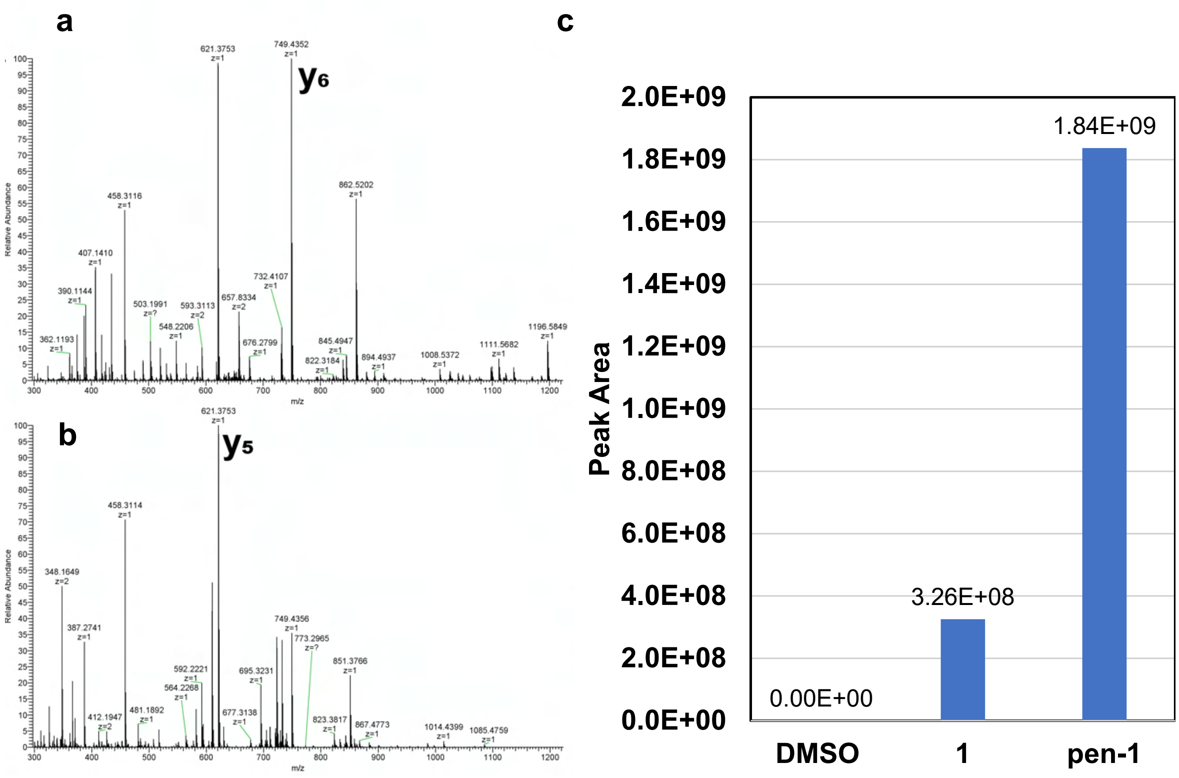

**Supplementary Figure 13.** Cyclic peptide M$^{pro}$ inhibitor **1** and the penetratin conjugate (**pen-1**) were incubated with HEK293-ACE2-TMPRSS2 cells. After 10 mins, cells were washed before lysis. Analysis of trypsin digested cell-lysates by data-independent acquisition (DIA) LC-MS/MS enabled the quantification by peak area of both **1** and **pen-1**. **a** Quantification of **1** used the y6 ion (749.4304, +1) in the 648.500 DIA window from precursor m/z 657.8292 (+2). **b** Quantification of **pen-1** used the y5 ion (621.3719, +1) in the 490.500 DIA window from precursor m/z 491.2503 (+3). **c pen-1** was observed to have >5.5-fold larger peak area compared to **1** under the same conditions. This shows that while both **pen-1** and **1** can enter cells to inhibit SARS-CoV-2 M$^{pro}$, **pen-1** does so more efficiently.

### Characterization of peptides

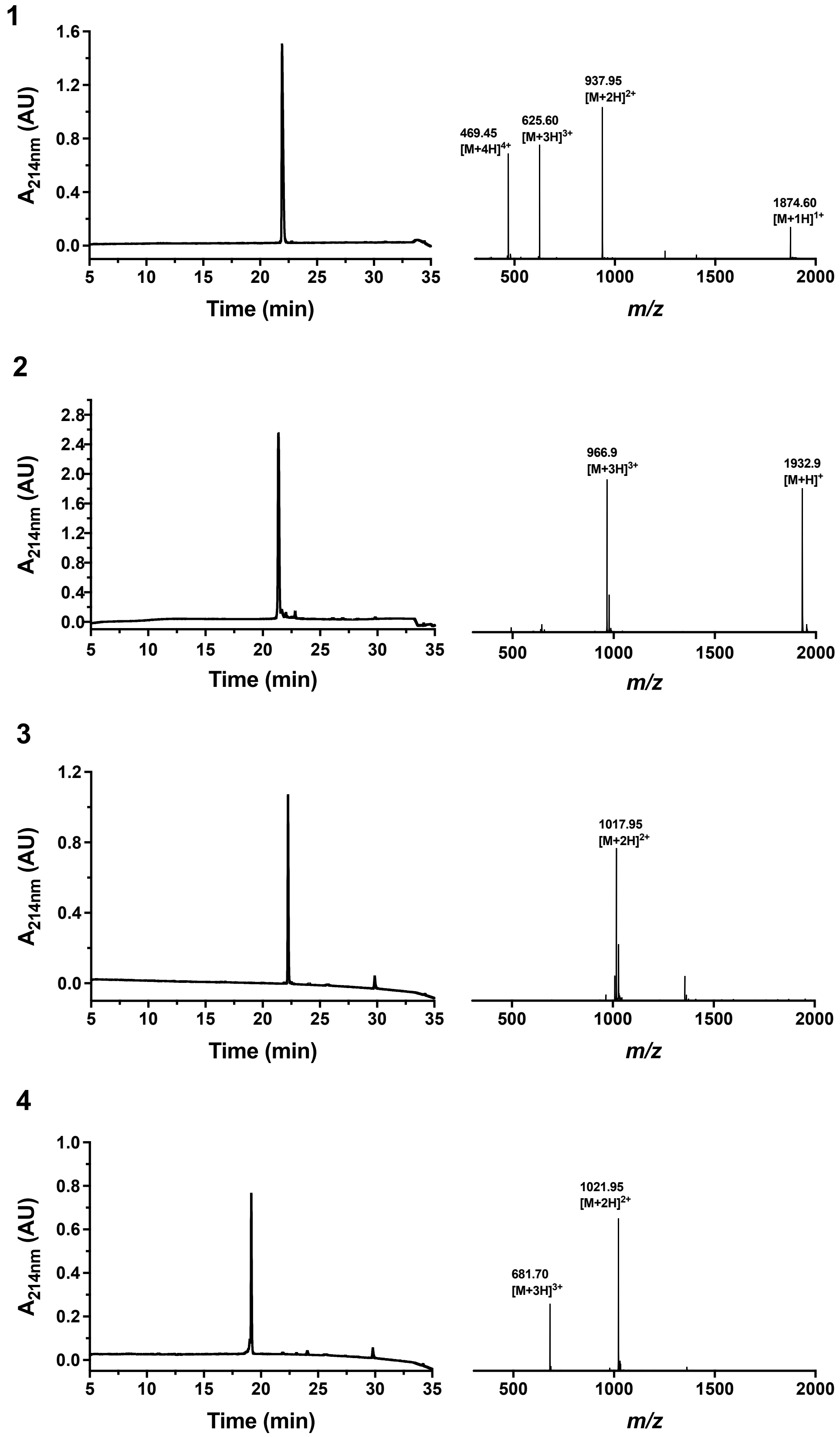

**Supplementary Figure 14. Analytical HPLC traces and ESI+ mass spectra of peptides 1-4.**

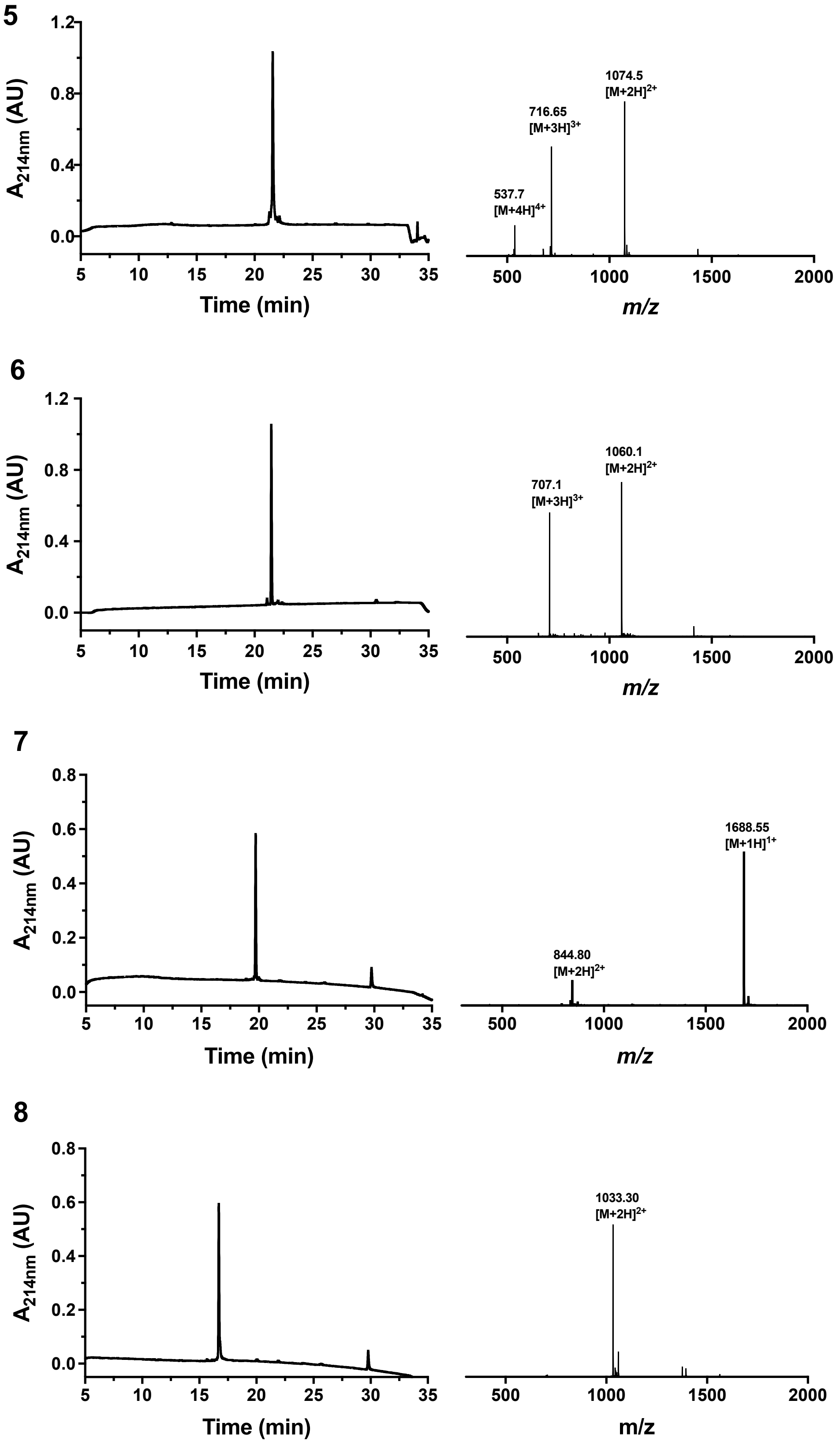

**Supplementary Figure 15. Analytical HPLC traces and ESI+ mass spectra of peptides 5-8.**

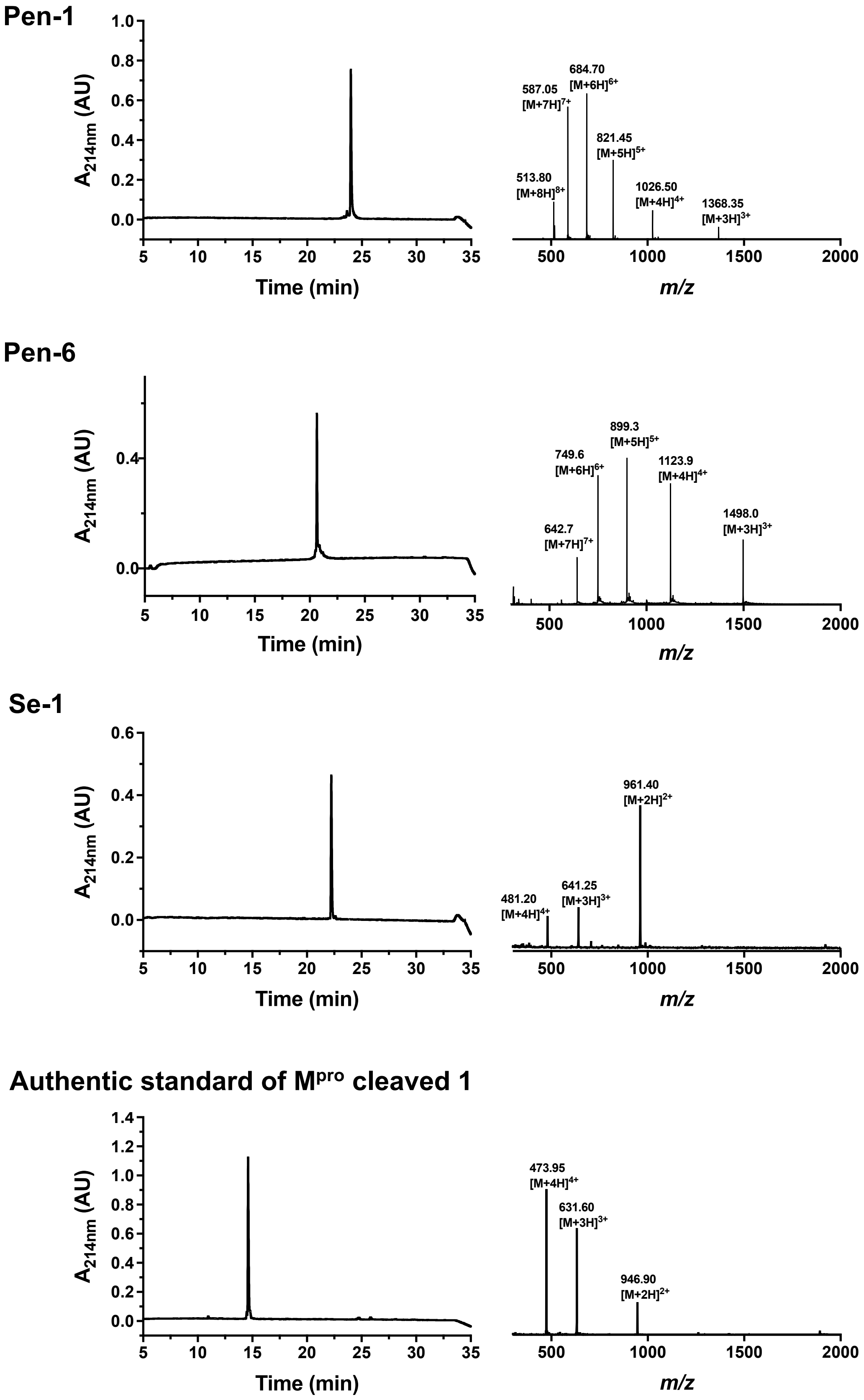

**Supplementary Figure 16. Analytical HPLC traces and ESI+ mass spectra of Pen-1, Pen-6, Se-1 and authentic standard of M$^{pro}$ cleaved 1.**

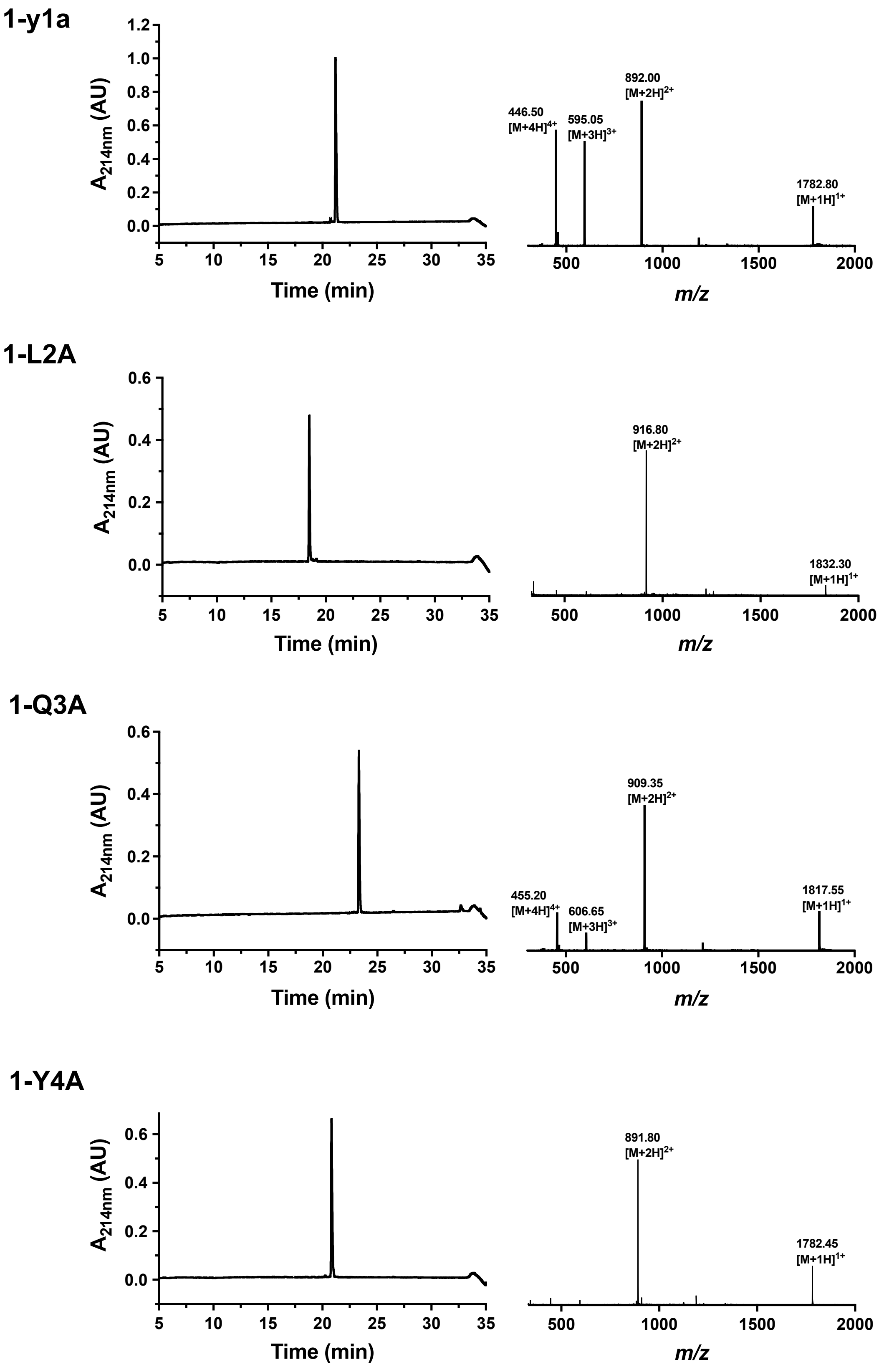

**Supplementary Figure 17. Analytical HPLC traces and ESI+ mass spectra of 1-y1a, 1-L2A, 1-Q3A and 1-Y4A.**

1-R8A

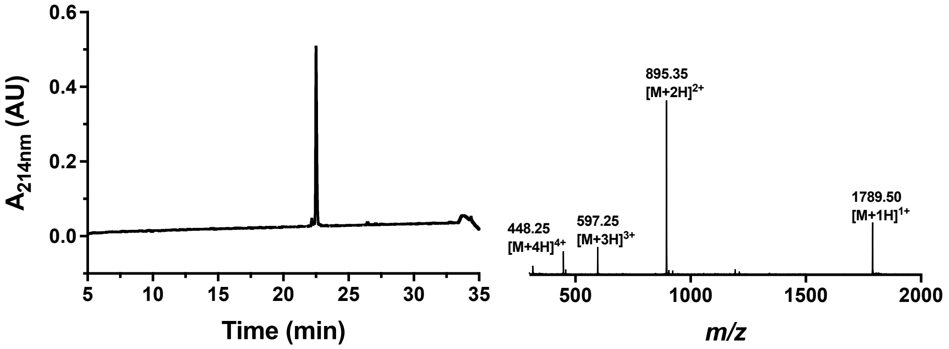

1-H9A

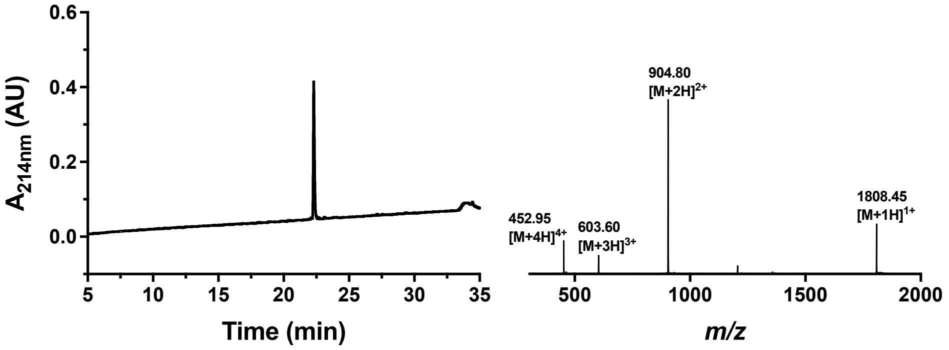

1-K10A

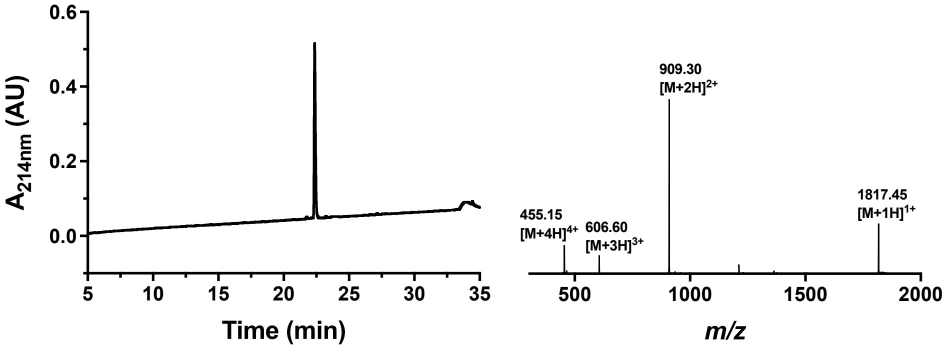

1-R11A

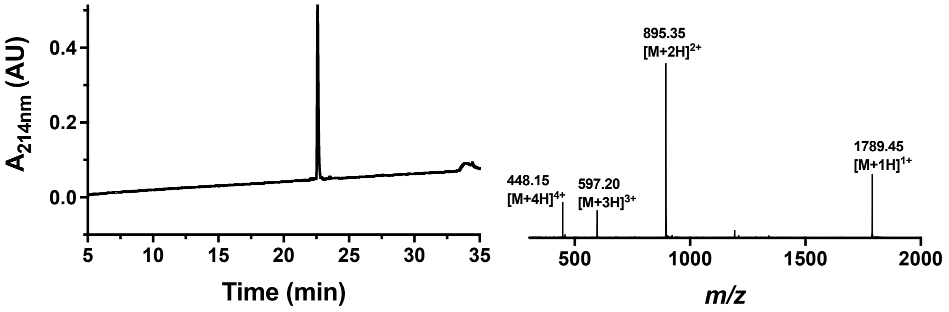

**Supplementary Figure 18. Analytical HPLC traces and ESI+ mass spectra of 1-R8A, 1-H9A, 1-K10A and 1-R11A.**

1-R12A

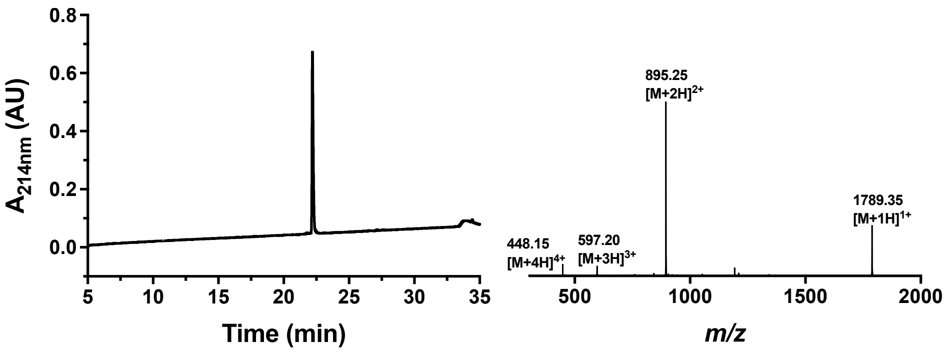

1-E13A

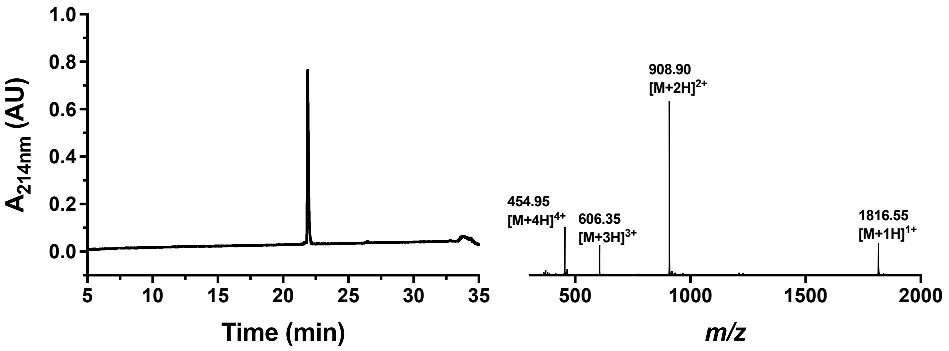

**Supplementary Figure 19. Analytical HPLC traces and ESI+ mass spectra of 1-R12A and 1-E13A.**

### Supplementary Methods

#### General Materials and Methods

Peptide grade *N,N*-dimethylformamide (DMF) and dichloromethane ($CH_2Cl_2$) for peptide synthesis were purchased from RCI Labscan and Merck, respectively. Gradient grade acetonitrile ($CH_3CN$) for chromatography was purchased from Sigma Aldrich and ultrapure water (Type 1) was obtained from a Merck Millipore Direct-Q 5 water purification system. Standard Fmoc-protected amino acids (Fmoc-Xaa-OH), coupling reagents and resins were purchased from Mimotopes and Novabiochem. PEG building blocks were purchased from Broadpharm. Fmoc-SPPS was performed manually with these reagents and solvents in polypropylene Teflon-fritted syringes purchased from Torviq or through automated synthesis on a SYRO I peptide synthesizer (Biotage). All other reagents were purchased from AK Scientific or Merck and used as received.

#### Preparative Chromatography

Preparative and semi-preparative reversed-phase high-performance liquid chromatography (RP-HPLC) was carried-out using a Waters 600E multisolvent delivery system with a Rheodyne 7725i injection valve fitted with a 5 mL loading loop, a Waters 500 pump and a Waters Fraction Collector III with detection using a Waters 490E programmable wavelength detector operating at 214 nm and 280 nm. Preparative RP-HPLC was carried-out using a Waters XBridge® C18 OBDTM Prep Column (5 µm, 30 x 150 mm) at a flow rate of 35 mL $min^{-1}$ using a mobile phase of $H_2O$ with 0.1vol.% TFA (solvent A) and $CH_3CN$ with 0.1vol.% TFA (solvent B) on linear gradients, unless otherwise specified. Semi-preparative RP-HPLC was performed using a Waters XBridge® BEH C18 OBDTM Prep Column (300 Å, 5 µm, 10 x 250 mm) at a flow rate of 5 mL $min^{-1}$ using a mobile phase of $H_2O$ with 0.1vol.% TFA (solvent A) and $CH_3CN$ with 0.1vol.% TFA (solvent B) on linear gradients, unless otherwise specified.

#### Liquid Chromatography-Mass Spectrometry

Liquid Chromatography-Mass Spectrometry (LC-MS) was performed on a Shimadzu 2020 UPLC-MS instrument with a Nexera X2 LC-30AD pump, Nexera X2 SPD-M30A

UV/Vis diode array detector and a Shimadzu 2020 (ESI) mass spectrometer operating in positive ion mode. Separations were performed on a Waters Acquity BEH300 1.7 µm, 2.1 × 50 mm (C18) column at a flow rate of 0.6 mL min$^{-1}$. All separations were performed using a mobile phase of 0.1 vol.% formic acid in water (solvent A) and 0.1 vol.% formic acid in $CH_3CN$ (solvent B) using linear gradients over 5 min.

**Analytical RP-HPLC**

Analytical RP-HPLC was performed on a Waters Alliance e2695 HPLC system equipped with a 2998 PDA detector ($\lambda$ = 210–400 nm). Separations were performed on a Waters XBridge® Peptide BEH300 5 µm, 4.6 × 250 mm (C18) column at 40 ºC with a flow rate of 1.0 mL min$^{-1}$. All separations were performed using a mobile phase of 0.1% TFA in water (Solvent A) and 0.1% TFA in $CH_3CN$ (Solvent B) using linear gradients, unless otherwise specified.

***In vitro* Protein Expression**

Selectively isotope-labelled samples of M$^{pro}$ R298A and site-directed mutants thereof were prepared by cell-free protein synthesis.[2,3] Selectively $^{15}$N-labelled samples were prepared with Ala, Cys, Gly, Ile, Leu, Lys, Met, Ser, Thr and Val. $^{15}$N-HSQC cross-peaks of selectively labelled samples were obtained by systematic site-directed mutagenesis, using cell-free protein synthesis from PCR-amplified DNA[4] to obtain 83 individual mutant protein samples (Supplementary Table 1). In each of these samples, the mutated residue type was labelled with $^{15}$N. Each cell-free reaction was conducted as two reactions, each with 1 mL inner and 10 mL outer buffer. The mutant proteins were purified using a 1 mL His GraviTrap™ TALON column (GE Healthcare, USA). Following the purification, the buffer was exchanged to NMR buffer and 10 % $D_2O$ added afterwards to provide a lock signal.

**NMR Experiments**

All NMR spectra of M$^{pro}$ R298A were recorded at 25 °C, using 3 mm NMR tubes on 800 MHz or 600 MHz Bruker Avance NMR spectrometers. 0.1–0.5 mM protein samples were used. Spectra recorded included 2D [$^{15}$N,$^{1}$H]-HSQC (typical parameters: $t_{1max}$ = 42 ms, $t_{2max}$ = 100 ms, total recording time 2.5 h) and [$^{15}$N,$^{1}$H]-TROSY ($t_{1max}$ = 82 ms, $t_{2max}$ = 142 ms, total recording time 2.2 h) and TROSY versions of 3D HNCA, HNCO, HNCACB and HNCOCA experiments. NOEs were recorded using a 3D NOESY-$^{15}$N-HSQC experiment with a mixing time of 150 ms.

**NMR Resonance Assignments of M$^{pro}$ R298A**

Selectively $^{15}$N-labelled samples were prepared by cell-free protein synthesis, which uses amino acids only sparingly (at a final concentration of 1 mM). Ten different amino acid types, one at a time, were targeted by using $^{15}$N-labelled amino acids. In addition, 83 samples were prepared with site-directed mutagenesis to remove single peaks from the [$^{15}$N,$^{1}$H]-HSQC spectrum of the selectively labelled samples. The side chains of all targeted residues were solvent exposed according to the crystal structure 6LU7.[5] 40 HSQC cross-peaks could be assigned in this way (Supplementary Tables 1 and 2). In the remaining cases, the assignment was compromised as the mutation perturbed the appearance of the [$^{15}$N,$^{1}$H]-HSQC spectrum too much or the cross-peak of the mutated amino acid was too weak to be observed in the first place. The resonance assignments made by site-directed mutation presented useful starting points for additional assignments made by 3D NMR spectra of $^{15}$N/$^{2}$H/$^{13}$C-labelled protein. Ultimately, we assigned 146 [$^{15}$N,$^{1}$H]-HSQC cross-peaks (Supplementary Figure 3b), which corresponds to almost half of the non-proline backbone amides. The assignments agree with previously published assignments made for the M$^{pro}$ dimer.[6]

**Molecular Dynamics Simulations**

The crystal structure of the cyclic peptide **Se-1** complexed with the SARS-CoV-2 M$^{pro}$ homodimer was used to model the full-length peptide. For chain A, the starting peptide conformation was manually built as accurately as possible using the available electron density, in the canonical orientation as seen in crystal structure. As a negative control, for chain B, the peptide was modelled in the opposite orientation, i.e. where the S2 subcavity was occupied by Tyr4 group (rather than Leu2).

The co-crystallized structure was processed using the protein preparation wizard in Maestro.[7] The sequence of steps involved in the preparation were the assignment of bond orders, addition of missing hydrogens, creating disulfide bonds, converting selenomethionines to methionines, and generation of het states using Epik program at a pH of 7.4 ± 0.02. The hydrogen bond networks were optimized using the default parameters and the PROPKA program was selected to assign the protonation states of the residues at a pH of 7.4. This was followed by restrained minimization, where the heavy atoms were converged to an RMSD of 0.30 Å, using the OPLS3e forcefield. Overlapping bonds were fixed using the 3-D builder tool in Maestro before an H-bond optimization step.

The structure generated after the restrained minimization step was selected for molecular dynamics (MD) studies using Desmond.[8] The Desmond System builder module was used to build the system, which involved enclosing the protein-ligand complex in an orthorhombic box with a buffer distance of (25x25x25) Å using the Transferable Intermolecular Potential 3P (TIP3P) water model. Counterions were added to neutralize the net charge, followed by the addition of 0.15 M NaCl to the system. The OPLS3e forcefield was selected for the study. The minimization step was carried out using the default parameters with a total simulation time of 100 ps.

The molecular dynamics module of Desmond was utilized to perform simulations of 330 ns duration in three replicates (0.99 μs in total) starting from different random seeds. The NPT ensemble was chosen with an initial temperature and pressure of 300 K and 1.01325 bar respectively. The time step was set to 2.0 fs, while the Nose-Hoover chain Langevin thermostat and the Martyna-Tobias-Klein barometer was selected to control the temperature and pressure, respectively, during the simulation. The Cutoff method was chosen to determine the short-range Coulombic interactions and the cut-off radius was set to 9 Å. The default NVT ensemble was used to run a short minimization to properly equilibrate the system. After completion, the trajectories for all the three simulations were analyzed using Simulation Event Analysis (SEA) module of Desmond to study the protein and ligand RMSD plots. The simulation interaction diagram (SID) program of Desmond was utilized to view the ligand-protein interactions occurring throughout the simulation run time.

The Trajectory Frame Clustering module of Desmond was used to generate representative structures from the combined trajectories. Clustering was performed based on the backbone of the complex, with the frequency set to 10 and the maximum number of the reported clusters set to 10. All the generated structures were analyzed and the representative model from each simulation was identified for detailed binding interaction analysis. The structural figures were produced from PyMOL.[9]

### Cyclic peptide synthesis

#### General procedure A; Automated Fmoc-Solid-Phase Peptide Synthesis (SPPS) – SYRO I automatic peptide synthesizer (Biotage)

Unless otherwise specified, peptides were synthesized on a 50 µmol scale. Rink amide resin (0.56 mmol $g^{-1}$, 1 eq.) was treated with a solution of piperidine (40 vol.%, 0.8 mL) in DMF for 3 min, drained, before repeat treatment with piperidine (20 vol.%, 0.8 mL) in DMF for 10 min. The resin was then drained and washed with DMF (4 x 1.2 mL) before addition of a solution of Fmoc-amino acid (200 µmol, 4 eq.) and Oxyma (4.4 eq.) in DMF (400 µL), followed by a solution of *N-N'*-diisopropylcarbodiimide (4 eq.) in DMF (400 µL). The resin was then agitated at 75 °C for 15 min or 50 °C for 30 min as specified (coupling of Fmoc-His(Trt)-OH and Fmoc-Cys(Trt)-OH were reacted at 50 °C for 30 min in all instances). The resin was then drained and a repeat treatment of the coupling conditions was conducted. The resin was then washed with DMF (4 x 1.2 mL) before being treated with a solution of 5 vol.% $Ac_2O$ and 10 vol.% *i*-$Pr_2NEt$ in DMF (1.6 mL) and agitated for 5 min to cap unreacted peptide N-termini. The resin was then drained and washed with DMF (4 x 1.6 mL). Iterative cycles of this process were repeated until complete peptide elongation was achieved after which the resin was washed with DMF (4 x 5 mL) and $CH_2Cl_2$ (5 x 5 mL).

#### General procedure B; Cyclisation ($CH_3CN$:$H_2O$)

The peptide was dissolved in 1:1 v/v $CH_3CN$:$H_2O$ (5 mM) in the presence of *i*-$Pr_2NEt$ (2.5 vol.%) and incubated for 1 h to facilitate thioether or selenoether cyclization.

#### General procedure C; Cyclisation (DMSO)

The peptide was dissolved in DMSO (5-10 mM) in the presence of *i*-$Pr_2NEt$ (2.5 vol.%) and incubated for 1 h to facilitate thioether cyclization.

**1:**

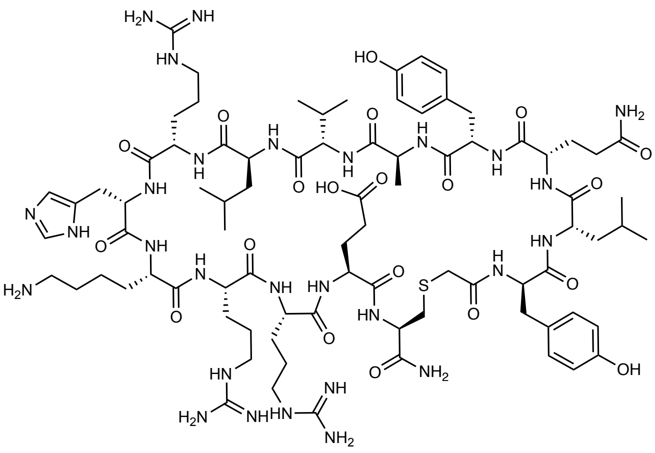

The peptide (50 µmol) was synthesized by general procedure **A** at 75 °C (Oxyma Pure was omitted from the terminal coupling of chloroacetic acid). The peptide was then cleaved from resin by treatment with 87.5:5:5:2.5 v/v/v/v trifluoroacetic acid/triisopropylsilane/$H_2O$/EDT for 3 h. The cleave solution was collected, dried to ~1 mL under $N_2$ flow and the peptide product precipitated from $Et_2O$ (30 mL) and collected by centrifugation. The crude linear peptide was then purified by preparative RP-HPLC (0 vol.% $CH_3CN$ + 0.1 vol.% TFA for 5 min, then 0-40 vol.% $CH_3CN$ + 0.1 vol.% TFA over 80 min, 35 mL/min, XBridge® C8, 300 Å, 30 x 150 mm). Fractions containing the linear peptide were combined and lyophilized. The lyophilized peptide was then cyclized by general procedure **B**. The reaction was concentrated under $N_2$ flow before being purified by preparative HPLC (0 vol.% $CH_3CN$ + 0.1 vol.% TFA for 5 min, then 0-30 vol.% $CH_3CN$ + 0.1 vol.% TFA over 80 min, 35 mL/min, XBridge® C8, 300 Å, 30 x 150 mm) to afford **1** as a white solid (12.6 mg, 10%). **$R_{t\ 214nm}$:** 21.93 min. (1 to 40 vol.% $CH_3CN$+ 0.1 vol.% TFA over 30 min). **LR-MS (+ESI):** m/z = 1874.60 $[M+H]^+$, 937.95 $[M+2H]^{2+}$, 625.60 $[M+3H]^{3+}$, 469.45 $[M+4H]^{4+}$.

**2:**

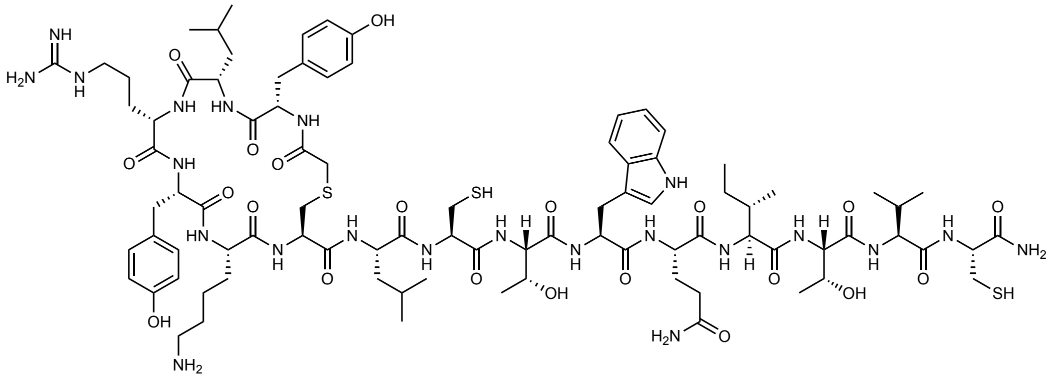

The peptide (50 µmol) was synthesized by general procedure **A** at 75 °C installing Cys8 and Cys15 as the acid stable Fmoc-Cys(Acm)-OH building block and coupling chloroacetic acid as the final residue. The peptide was then cleaved from resin by treatment with 90:5:5 v/v/v trifluoroacetic acid/triisopropylsilane/$H_2O$ for 2 h, after which the cleave solution was collected, dried to ~1 mL under $N_2$ flow and the peptide product precipitated from $Et_2O$ (30 mL) and collected by centrifugation. The crude peptide was then cyclized by general procedure **B** before addition of AgOAc (66 mg, 400 µmol). The reaction was shaken for 16 h before addition of dithiothreitol (90 mg, 600 µmol) causing generation of a precipitate that was removed by filtration and the filtrate lyophilized. The product was purified by semi-preparative HPLC (0 vol.% $CH_3CN$ + 0.1 vol.% TFA for 5 min, then 0-60 vol.% $CH_3CN$ + 0.1 vol.% TFA over 40 min) to afford the pure peptide as a white amorphous solid (1.2 mg, 1.1%). **$R_{t\ 214nm}$:** 21.35 min. (1 to 60 vol.% $CH_3CN$+ 0.1 %vol TFA over 30 min). **LR-MS (+ESI):** m/z = 1932.9 $[M+H]^+$, 966.9 $[M+2H]^2$.

**3:**

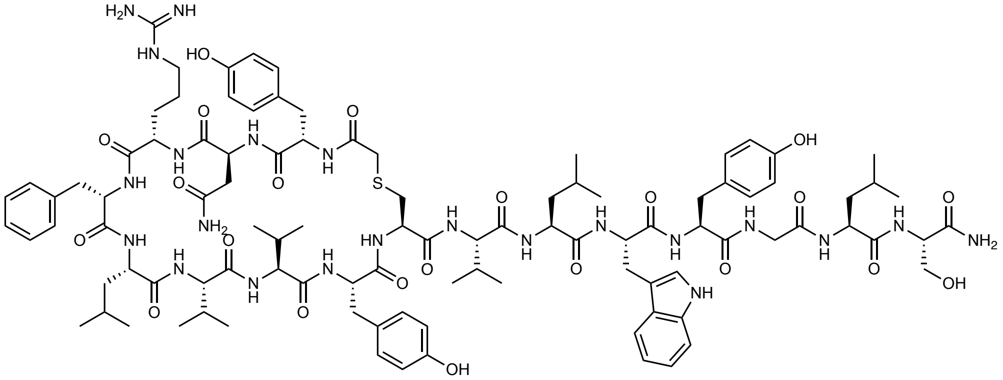

The peptide (50 µmol) was synthesized by general procedure **A** at 75 °C. The peptide was then cleaved from resin by treatment with 85:5:5:2.5:2.5 v/v/v/v/v trifluoroacetic acid/triisopropylsilane/$H_2O$/phenol/EDT for 2 h. The cleave solution was collected, dried to ~1 mL under $N_2$ flow and the peptide product precipitated from $Et_2O$ (2 x 40 mL) and collected by centrifugation. The peptide was then lyophilized from 1:1 v/v $CH_3CN$:$H_2O$, 0.1 vol.% TFA before being dissolved in 3:2 v/v $H_2O$/DMF (50 mL) and cyclized with 4 vol.% *i*-$Pr_2NEt$ for 2 h. The solution was then concentrated by $N_2$ flow and purified by semi-preparative RP-HPLC (0 vol.% $CH_3CN$ + 0.1 vol.% formic acid for 5 min, then 15-50 vol.% $CH_3CN$ + 0.1 vol.% formic acid over 80 min) to afford **3** as a white solid (1.6 mg, 1%). **$R_{t\ 214nm}$:** 22.22 min. (1 to 80 vol.% $CH_3CN$ + 0.1 %vol TFA over 30 min). **LR-MS (+ESI):** *m/z* = 1017.95 $[M+2H]^{2+}$.

**4.**

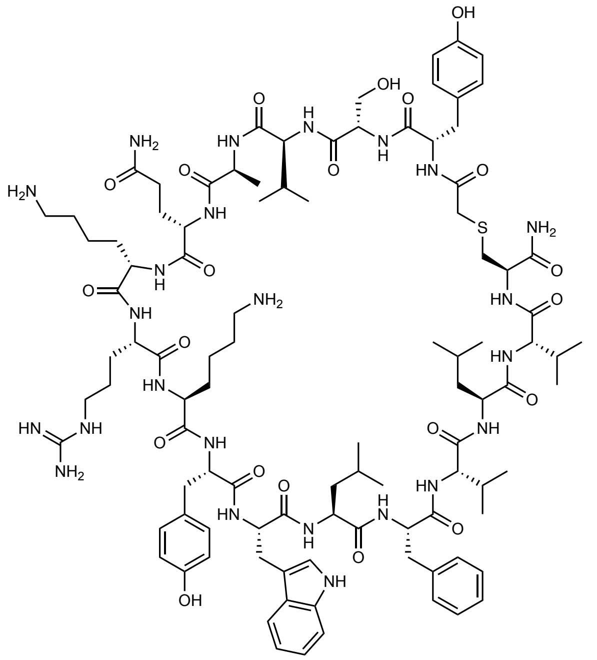

The peptide (50 µmol) was synthesized by general procedure **A** at 75 °C, with the exception that K6, R7 and K9 were subjected to a third treatment at the amino acid coupling step. The peptide was then cleaved from resin with 85:5:5:2.5:2.5 v/v/v/v/v TFA/TIS/$H_2O$/phenol/EDT for 2 h. The cleave solution was collected, dried to ~1 mL under $N_2$ flow and the peptide product precipitated from $Et_2O$ over dry ice (2 x 40 mL) and collected by centrifugation. The crude peptide was then lyophilized and purified by preparative RP-HPLC (0 vol.% $CH_3CN$ + 0.1 vol.% formic acid for 5 min, then 0-50 vol.% $CH_3CN$ + 0.1 vol.% formic acid over 60 min). The linear peptide was then cyclized by general procedure **B**. The cyclic peptide solution was then dried under $N_2$ flow and purified by semi-preparative RP-HPLC (0 vol.% $CH_3CN$ + 0.1 vol.% TFA acid for 5 min, then 0-50 vol.% $CH_3CN$ + 0.1 vol.% TFA over 80 min) to afford **4** as a white amorphous solid (1.0 mg, 0.8%). **$R_{t\,214nm}$:** 19.15 min. (1 to 80 vol.% $CH_3CN$ + 0.1 %vol TFA over 30 min). **LR-MS (+ESI):** *m/z* = 1021.95 $[M+2H]^{2+}$, 681.70 $[M+3H]^{3+}$.

**5.**

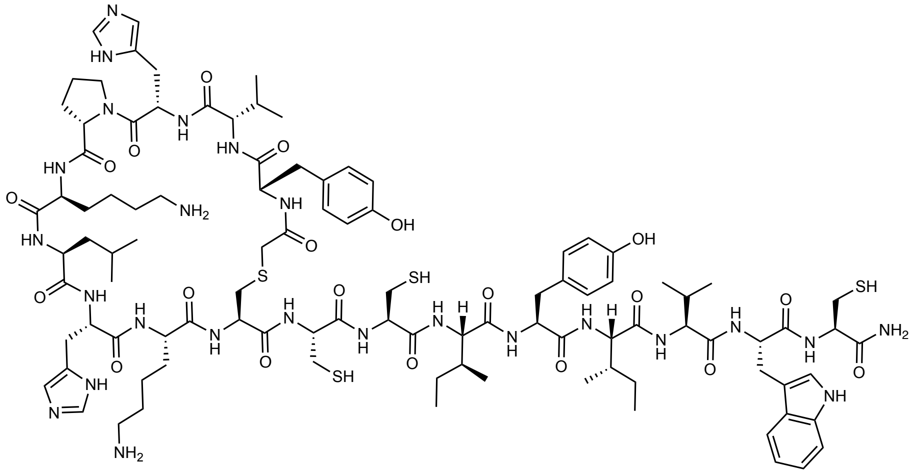

The peptide (50 µmol) was synthesized by general procedure **A** installing Cys10, Cys11 and Cys17 as the acid stable Fmoc-Cys(Acm)-OH building block and coupling chloroacetic acid as the final residue. The peptide was then cleaved from half of the resin (25 µmol) by treatment with 90:5:5 v/v/v trifluoroacetic acid/triisopropylsilane/$H_2O$ for 2 h. The crude peptide was then cyclized by general procedure **B** before being purified by HPLC (0 vol.% $CH_3CN$ + 0.1 vol.% TFA acid for 5 min, then 0-60 vol.% $CH_3CN$ + 0.1 vol.% TFA over 30 min) to afford the Acm-protected peptide, which was subsequently dissolved in $H_2O$ with 0.1 vol.% TFA and AgOAc (20.8 mg, 125 µmol) added. The reaction was shaken for 1 h before addition of dithiothreitol (38.6 mg, 250 µmol) causing generation of a precipitate that was removed by filtration and the filtrate lyophilized. The product was purified by semi-preparative HPLC (0 vol.% $CH_3CN$ + 0.1 vol.% TFA acid for 5 min, then 0-60 vol.% $CH_3CN$ + 0.1 vol.% TFA over 30 min) to afford **5** as a white amorphous solid (1.9 mg, 2.9%). **$R_{t\,214nm}$:** 21.56 min. (1 to 60 vol.% $CH_3CN$ + 0.1 %vol TFA over 30 min). **LR-MS (+ESI):** *m/z* = 1074.5 $[M+2H]^{2+}$, 716.65 $[M+3H]^{3+}$, 537.7 $[M+4H]^{4+}$.

**6.**

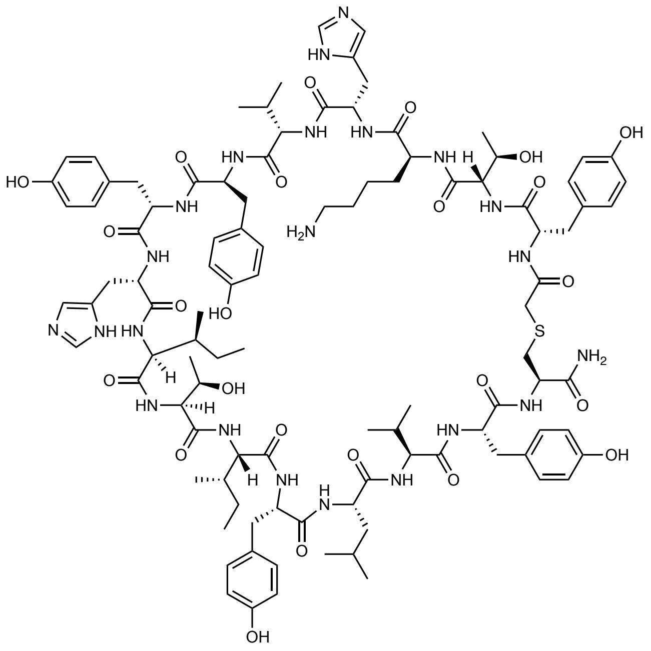

The peptide (50 μmol) was synthesized by general procedure **A** at 75°C. The peptide was then cleaved from resin by treatment with 90:5:5 v/v/v trifluoroacetic acid/triisopropylsilane/$H_2O$ for 1.5 h, after which the cleave solution was collected, dried to ~1 mL under $N_2$ flow and the peptide product precipitated from $Et_2O$ (30 mL) and collected by centrifugation. The crude peptide was then purified by RP-HPLC (0 vol.% $CH_3CN$ + 0.1 vol.% TFA acid for 5 min, then 20-60 vol.% $CH_3CN$ + 0.1 vol.% TFA over 40 min, 15 mL/min, Waters Symmetry C4, 300 Å, 5 μm, 4.6 mm x 250 mm). Fractions containing the linear peptide were combined and lyophilized. The crude peptide was then cyclized by general procedure **C**. The product was then purified by RP-HPLC (0 vol.% $CH_3CN$ + 0.1 vol.% formic acid for 5 min, then 20-60 vol.% $CH_3CN$ + 0.1 vol.% formic acid over 40 min, 38 mL/min, XBridge® C8, 300 Å, 30 x 150 mm) to afford the pure peptide as a white solid (3.8 mg, 3.1%). **$R_{t\ 214nm}$:** 21.44 min. (1 to 60 vol.% $CH_3CN$ + 0.1 %vol TFA over 30 min). **LR-MS (+ESI):** *m/z* = 1060.1 $[M+2H]^{2+}$, 707.1 $[M+3H]^{3}$.

**7.**

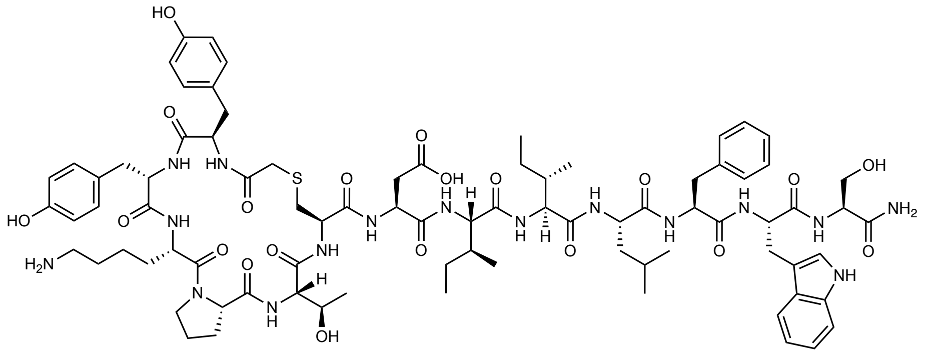

The peptide (50 µmol) was synthesized by general procedure **A** at 75 °C. The peptide was then cleaved from resin by treatment with 87.5:5:5:2.5 v/v/v/v trifluoroacetic acid/triisopropylsilane/$H_2O$/EDT for 2 h. The cleave solution was collected, dried to ~1 mL under $N_2$ flow and the peptide product precipitated from $Et_2O$ (2 x 20 mL) and collected by centrifugation. The peptide was then cyclized by general procedure **B** before being concentrated by $N_2$ flow and purified by preparative RP-HPLC (0 vol.% $CH_3CN$ + 0.1 vol.% TFA for 5 min, then 5-45 vol.% $CH_3CN$ + 0.1 vol.% TFA over 60 min) to afford **7** as a white solid (14.4 mg, 16%). $\mathbf{R_{t\ 214nm}}$**:** 19.75 min. (1 to 80 vol.% $CH_3CN$ + 0.1 %vol TFA over 30 min). **LR-MS (+ESI):** *m/z* = 1688.55 $[M+H]^+$, 844.80 $[M+2H]^{2+}$.

**8.**

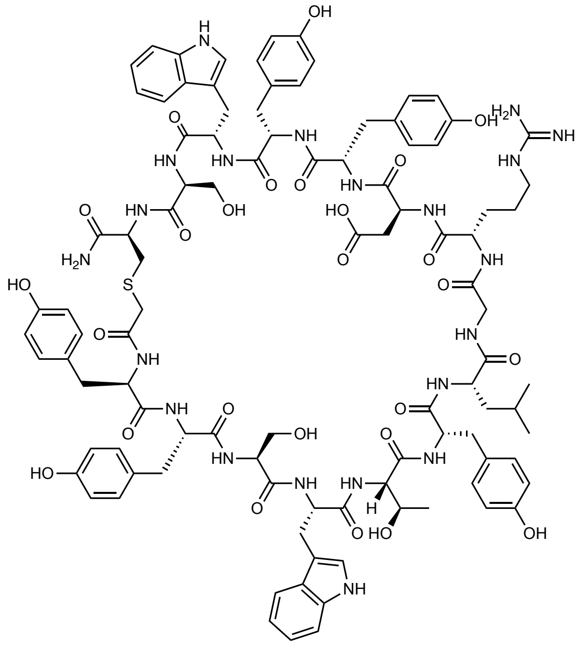

The peptide (50 µmol) was synthesized by general procedure **A** at 75 °C (Oxyma Pure was omitted from the terminal coupling of chloroacetic acid). The peptide was then cleaved from resin by treatment with 87.5:5:5:2.5 v/v/v/v trifluoroacetic acid/triisopropylsilane/$H_2O$/EDT for 2 h. The cleave solution was collected, dried to ~1 mL under $N_2$ flow and the peptide product precipitated from $Et_2O$ (2 x 20 mL) and collected by centrifugation. The peptide was then cyclized by general procedure **B** before being concentrated by $N_2$ flow and purified by preparative RP-HPLC (0 vol.% $CH_3CN$ + 0.1 vol.% TFA for 5 min, then 5-45 vol.% $CH_3CN$ + 0.1 vol.% TFA over 60 min) to afford **8** as a white solid (26.9 mg, 25%). **$R_{t\ 214nm}$:** 16.71 min. (1 to 80 vol.% $CH_3CN$ + 0.1 %vol TFA over 30 min). **LR-MS (+ESI):** *m/z* = 1033.30 $[M+2H]^{2+}$.

**Pen-1.**

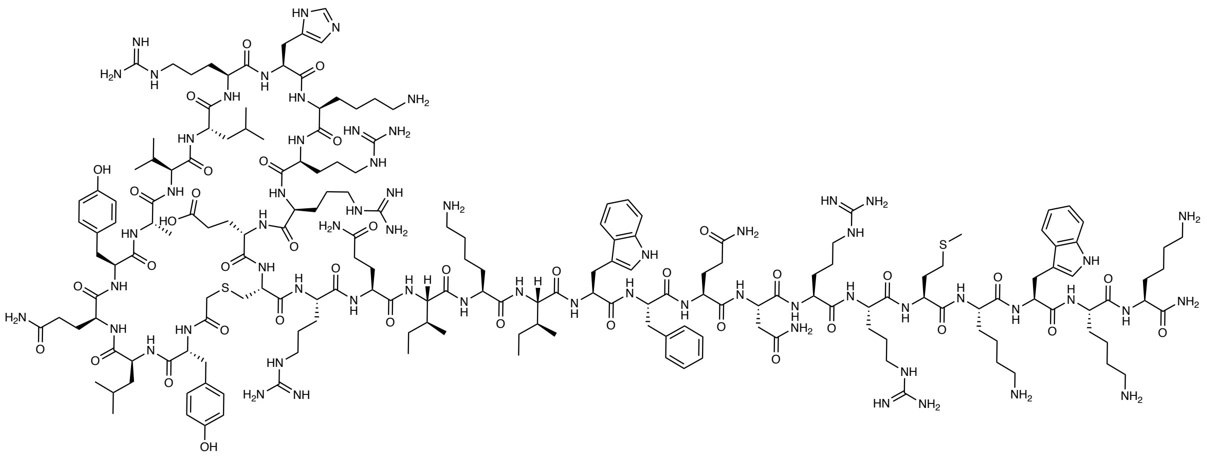

The peptide (25 µmol) was synthesized by general procedure **A** (Oxyma Pure was omitted from the terminal coupling of chloroacetic acid). The peptide was then cleaved from resin by treatment with 87.5:5:5:2.5 v/v/v/v trifluoroacetic acid/triisopropylsilane/$H_2O$/EDT for 3 h. The cleave solution was collected, dried to ~1 mL under $N_2$ flow and the peptide product precipitated from $Et_2O$ (2 x 40 mL) and collected by centrifugation. The crude peptide was then purified by preparative RP-HPLC (0 vol.% $CH_3CN$ + 0.1 vol.% TFA for 5 min, then 0-40 vol.% $CH_3CN$ + 0.1 vol.% TFA over 80 min, 35 mL/min, XBridge® C8, 300 Å, 30 x 150 mm). The peptide was then cyclized by general procedure **B** with the addition of TCEP (2.5 mg, 10 µmol) and concentrated under $N_2$ flow before purification by preparative RP-HPLC (0 vol.% $CH_3CN$ + 0.1 vol.% TFA for 5 min, then 5-30 vol.% $CH_3CN$ + 0.1 vol.% TFA over 80 min, 15 mL/min, XBridge® C18, 300 Å, 19 x 150 mm) to afford **Pen-1** as a white solid (5.1 mg, 3%). **$R_{t\ 214nm}$:** 24.16 min. (1 to 40 vol.% $CH_3CN$+ 0.1 %vol TFA over 30 min). **LR-MS (+ESI):** m/z = 1368.35 $[M+3H]^{3+}$, 1026.50 $[M+4H]^{4+}$, 821.45 $[M+5H]^{5+}$, 684.70 $[M+6H]^{6+}$, 587.05 $[M+7H]^{7+}$, 513.80 $[M+8H]^{8+}$.

**Pen-6**

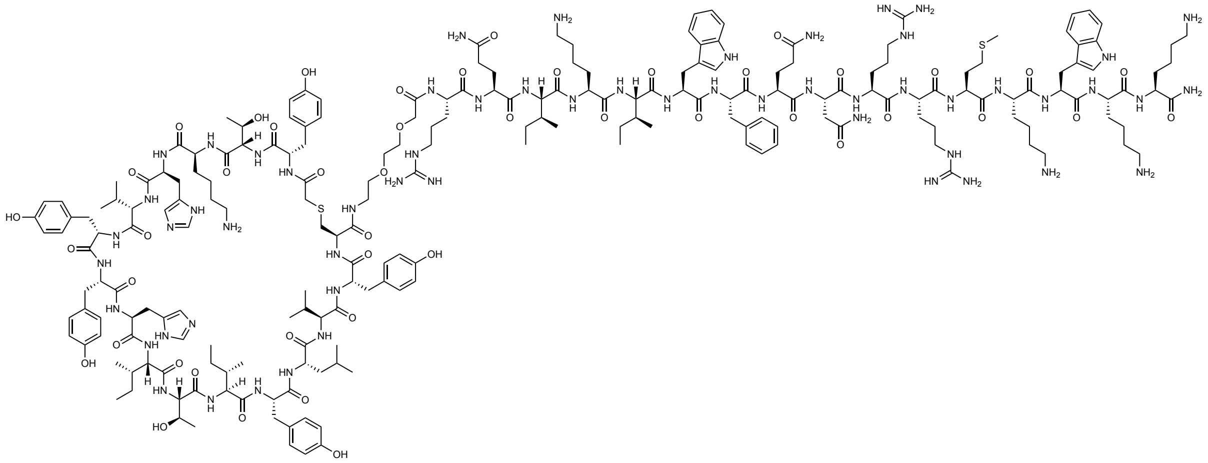

The peptide (12.5 μmol) was synthesized by general procedure **A**. The peptide was then cleaved from resin by treatment with 90:5:5 v/v/v trifluoroacetic acid/triisopropylsilane/$H_2O$ for 1.5 h, after which the cleave solution was collected, dried to ~1 mL under $N_2$ flow and the peptide product precipitated from $Et_2O$ (30 mL) and collected by centrifugation. The crude peptide was then cyclized by general procedure **C** before being purified by preparative RP-HPLC (0 vol.% $CH_3CN$ + 0.1 vol.% TFA for 5 min, then 0-40 vol.% $CH_3CN$ + 0.1 vol.% TFA over 40 min, 38 mL/min, XBridge® C8, 300 Å, 30 x 150 mm) to afford **Pen-6** as a white amorphous solid (2.31 mg, 3.3%). **$R_{t\ 214nm}$:** 20.64 min. (1 to 60 vol.% $CH_3CN$+ 0.1 %vol TFA over 30 min). **LR-MS (+ESI):** m/z = 1498.0 $[M+3H]^{3+}$, 1123.9 $[M+4H]^{4+}$, 899.3 $[M+5H]^{5+}$, 749.6 $[M+6H]^{6+}$, 642.7 $[M+7H]^{7+}$.

**1-y1a**

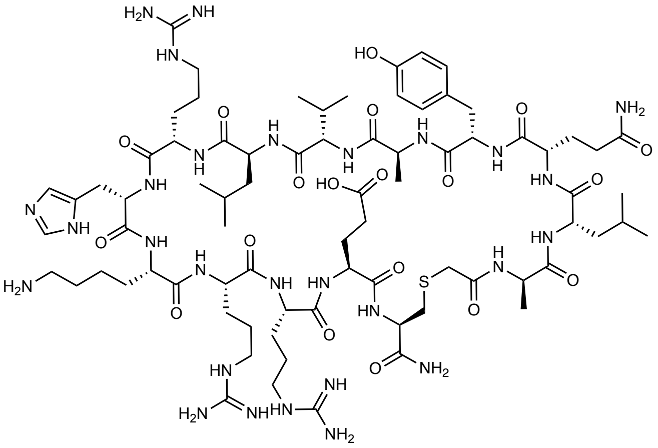

The peptide (50 µmol) was synthesized by general procedure **A** at 50 °C (Oxyma Pure was omitted from the terminal coupling of chloroacetic acid). The peptide was then cleaved from resin by treatment with 87.5:5:5:2.5 v/v/v/v trifluoroacetic acid/triisopropylsilane/$H_2O$/EDT for 2 h. The cleave solution was collected, dried to ~1 mL under $N_2$ flow and the peptide product precipitated from $Et_2O$ (2 x 20 mL) and collected by centrifugation. The crude peptide was then purified by RP-HPLC (100% $H_2O$ with 0.1 vol.% TFA over 5 min then 0 to 30 vol.% $CH_3CN$ in $H_2O$ with 0.1 vol.% TFA over 40 min XBridge® C8, 300 Å, 30 x 150 mm). The peptide was then cyclized by general procedure **B** before being lyophilized and purified by RP-HPLC (0 vol.% $CH_3CN$ + 0.1 vol.% formic acid for 5 min, then 0-25 vol.% $CH_3CN$ + 0.1 vol.% formic acid over 60 min, 15 mL/min, XBridge® C18, 300 Å, 19 x 150 mm) to afford **1-y1a** as a white solid (12.9 mg, 12%). **$R_{t\ 214nm}$:** 21.19 min. (1 to 40 vol.% $CH_3CN$+ 0.1 vol.% TFA over 30 min). **LR-MS (+ESI):** m/z = 1782.80 $[M+H]^{+}$, 892.00 $[M+2H]^{2+}$, 595.05 $[M+3H]^{3+}$, 446.50 $[M+4H]^{4+}$.

**1-L2A**

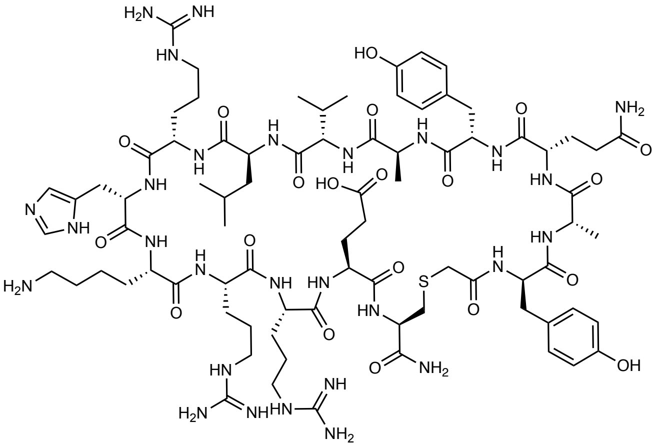

The peptide (50 µmol) was synthesized by general procedure **A** at 50 °C. The final coupling was performed with chloroacetic acid (37.8 mg, 448 µmol, 8 equiv.) and *N-N'*-diisopropylcarbodiimide (70 µL, 448 µmol, 8 equiv.) for 12 h at rt. The peptide was then cleaved from resin by treatment with 87.5:5:5:2.5 v/v/v/v trifluoroacetic acid/triisopropylsilane/$H_2O$/EDT for 2 h. The cleave solution was collected, dried to ~1 mL under $N_2$ flow and the peptide product precipitated from $Et_2O$ (2 x 20 mL) and collected by centrifugation. The crude peptide was then purified by RP-HPLC (0 vol.% $CH_3CN$ + 0.1 vol.% TFA for 5 min, then 0-40 vol.% $CH_3CN$ + 0.1 vol.% TFA over 60 min). The peptide was then cyclized by general procedure **B** before being lyophilized and purified by RP-HPLC (0 vol.% $CH_3CN$ + 0.1 vol.% TFA for 5 min, then 0-35 vol.% $CH_3CN$ + 0.1 vol.% TFA over 60 min) to afford **1** as a white solid (11.0 mg, 8%). **$R_t$ $_{214nm}$:** 18.48 min. (1 to 40 vol.% $CH_3CN$+ 0.1 %vol TFA over 30 min, 1 mL/min, 50 ºC). **LR-MS (+ESI):** m/z = 1832.30 $[M+1H]^{1+}$, 916.80 $[M+2H]^{2+}$.

**1-Q3A**

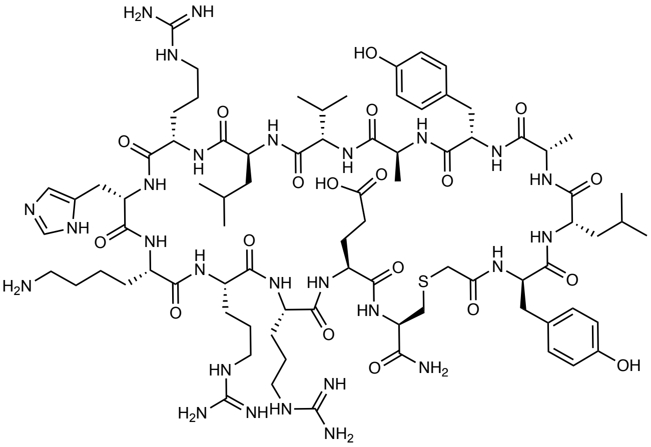

The peptide (50 µmol) was synthesized by general procedure **A** at 50 °C (Oxyma Pure omitted from the terminal coupling of chloroacetic acid). The peptide was then cleaved from resin by treatment with 87.5:5:5:2.5 v/v/v/v trifluoroacetic acid/triisopropylsilane/$H_2O$/EDT for 2 h. The cleave solution was collected, dried to ~1 mL under $N_2$ flow and the peptide product precipitated from $Et_2O$ (2 x 30 mL) and collected by centrifugation. The crude peptide was then cyclized by general procedure **B** before being concentrated under $N_2$ flow and purified by preparative RP-HPLC (0 vol.% $CH_3CN$ + 0.1 vol.% formic acid for 5 min, then 0-30 vol.% $CH_3CN$ + 0.1 vol.% formic acid over 80 min, 15 mL/min, XBridge® C18, 300 Å, 19 x 150 mm) to afford **1** as a white solid (5.2 mg, 5%). **$R_{t\ 214nm}$:** 23.41 min, (1 to 40 vol.% $CH_3CN$+ 0.1 vol.% TFA over 30 min). **LR-MS (+ESI):** m/z = 1817.55 $[M+H]^{+}$, 909.35 $[M+2H]^{2+}$, 606.65 $[M+3H]^{3+}$, 455.20 $[M+4H]^{4+}$.

**1-Y4A**

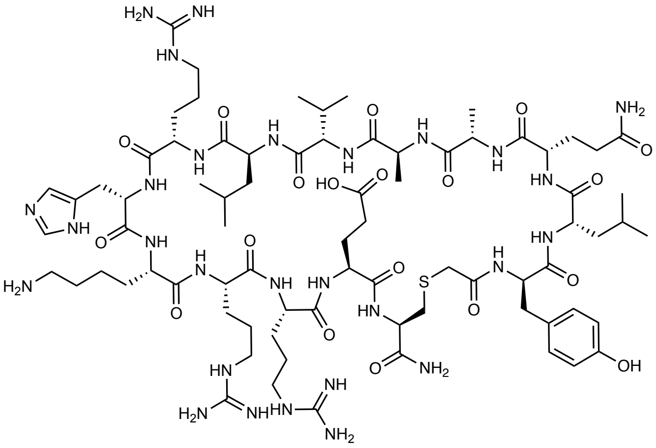

The peptide (50 µmol) was synthesized by general procedure **A** at 50 °C (Oxyma Pure omitted from the terminal coupling of chloroacetic acid). The peptide was then cleaved from resin by treatment with 87.5:5:5:2.5 v/v/v/v trifluoroacetic acid/triisopropylsilane/$H_2O$/EDT for 2 h. The cleave solution was collected, dried to ~1 mL under $N_2$ flow and the peptide product precipitated from $Et_2O$ (2 x 20 mL) and collected by centrifugation. The crude peptide was then purified by preparative RP-HPLC (0 vol.% $CH_3CN$ + 0.1 vol.% TFA for 5 min, then 0-40 vol.% $CH_3CN$ + 0.1 vol.% TFA over 60 min). The peptide was then cyclized by general procedure **B** before being lyophilized and purified by preparative RP-HPLC (0 vol.% $CH_3CN$ + 0.1 vol.% TFA for 5 min, then 5-35 vol.% $CH_3CN$ + 0.1 vol.% TFA over 60 min) to afford **1** as a white solid (6.7 mg, 5%). **$R_{t\ 214nm}$:** 20.84 min. (1 to 40 vol.% $CH_3CN$+ 0.1 %vol TFA over 30 min). **LR-MS (+ESI):** m/z = 1782.45 $[M+H]^+$, 891.80 $[M+2H]^{2+}$.

**1-R8A**

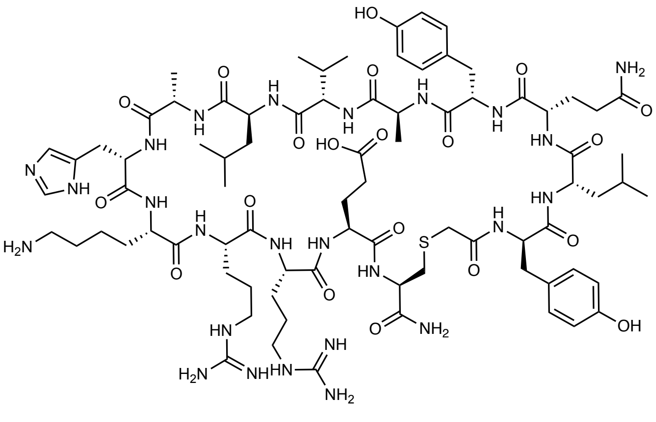

The peptide (50 µmol) was synthesized by general procedure **A** (Oxyma Pure was omitted from the terminal coupling of chloroacetic acid). The peptide was then cleaved from resin by treatment with 87.5:5:5:2.5 v/v/v/v trifluoroacetic acid/triisopropylsilane/$H_2O$/EDT for 2 h. The cleave solution was collected, dried to ~1 mL under $N_2$ flow and the peptide product precipitated from $Et_2O$ (2 x 40 mL) and collected by centrifugation. The crude peptide was then purified by preparative RP-HPLC (0 vol.% $CH_3CN$ + 0.1 vol.% TFA for 5 min, then 5-30 vol.% $CH_3CN$ + 0.1 vol.% TFA over 60 min). Fractions containing the linear peptide were combined and lyophilized. The peptide was then cyclized by general procedure **B**. The reaction was concentrated under $N_2$ flow before being purified by preparative RP-HPLC (0 vol.% $CH_3CN$ + 0.1 vol.% TFA for 5 min, then 8-30 vol.% $CH_3CN$ + 0.1 vol.% TFA over 60 min, 15 mL/min, XBridge® C18, 300 Å, 19 x 150 mm) to afford **1** as a white solid (12.0 mg, 10%). **$R_{t\ 214nm}$:** 22.51 min. (1 to 40 vol.% $CH_3CN$+ 0.1 vol.% TFA over 30 min). **LR-MS (+ESI):** m/z = 1789.50 $[M+H]^+$, 895.35 $[M+2H]^{2+}$, 597.25 $[M+3H]^{3+}$, 448.25 $[M+4H]^{4+}$.

**1-H9A**

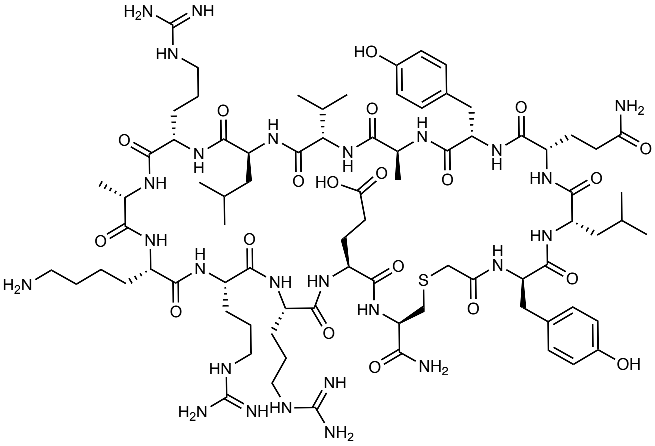

The peptide (50 μmol) was synthesized by general procedure **A** (Oxyma Pure was omitted from the terminal coupling of chloroacetic acid). The peptide was then cleaved from resin by treatment with 87.5:5:5:2.5 v/v/v/v trifluoroacetic acid/triisopropylsilane/$H_2O$/EDT for 2 h. The cleave solution was collected, dried to ~1 mL under $N_2$ flow and the peptide product precipitated from $Et_2O$ (2 x 40 mL) and collected by centrifugation. The crude peptide was then purified by preparative RP-HPLC (0 vol.% $CH_3CN$ + 0.1 vol.% TFA for 5 min, then 5-30 vol.% $CH_3CN$ + 0.1 vol.% TFA over 60 min, 35 mL/min, XBridge® C8, 300 Å, 30 x 150 mm). Fractions containing the linear peptide were combined and lyophilized. The peptide was then cyclized by general procedure **B**. The reaction was concentrated under $N_2$ flow before being purified by preparative RP-HPLC (0 vol.% $CH_3CN$ + 0.1 vol.% TFA for 5 min, then 5-25 vol.% $CH_3CN$ + 0.1 vol.% TFA over 60 min) to afford **1** as a white solid (11.5 mg, 10%). **$R_{t\ 214nm}$:** 22.30 min. (1 to 40 vol.% $CH_3CN$+ 0.1 vol.% TFA over 30 min, 1 mL/min). **LR-MS (+ESI):** m/z = 1808.45 $[M+H]^+$, 904.80 $[M+2H]^{2+}$, 603.60 $[M+3H]^{3+}$, 452.95 $[M+4H]^{4+}$.

**1-K10A**

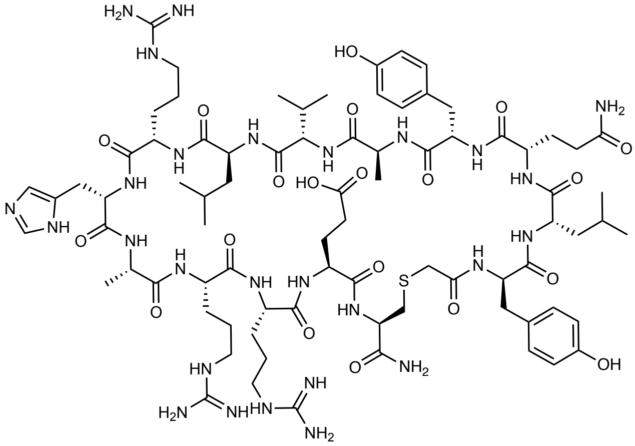

The peptide (50 µmol) was synthesized by general procedure **A** (Oxyma Pure was omitted from the terminal coupling of chloroacetic acid). The peptide was then cleaved from resin by treatment with 87.5:5:5:2.5 v/v/v/v trifluoroacetic acid/triisopropylsilane/$H_2O$/EDT for 2 h. The cleave solution was collected, dried to ~1 mL under $N_2$ flow and the peptide product precipitated from $Et_2O$ (2 x 40 mL) and collected by centrifugation. The crude peptide was then purified by preparative RP-HPLC (0 vol.% $CH_3CN$ + 0.1 vol.% TFA for 5 min, then 5-30 vol.% $CH_3CN$ + 0.1 vol.% TFA over 40 min). Fractions containing the linear peptide were combined and lyophilized. The peptide was then cyclized by general procedure **B**. The reaction was concentrated under $N_2$ flow before being purified by preparative RP-HPLC (0 vol.% $CH_3CN$ + 0.1 vol.% TFA for 5 min, then 5-25 vol.% $CH_3CN$ + 0.1 vol.% TFA over 60 min) to afford **1** as a white solid (14.2 mg, 13%). **$R_{t\ 214nm}$:** 22.37 min. (1 to 40 vol.% $CH_3CN$+ 0.1 %vol TFA over 30 min). **LR-MS (+ESI):** m/z = 1817.45 $[M+H]^+$, 909.30 $[M+2H]^{2+}$, 606.60 $[M+3H]^{3+}$, 455.15 $[M+4H]^{4+}$.

**1-R11A**

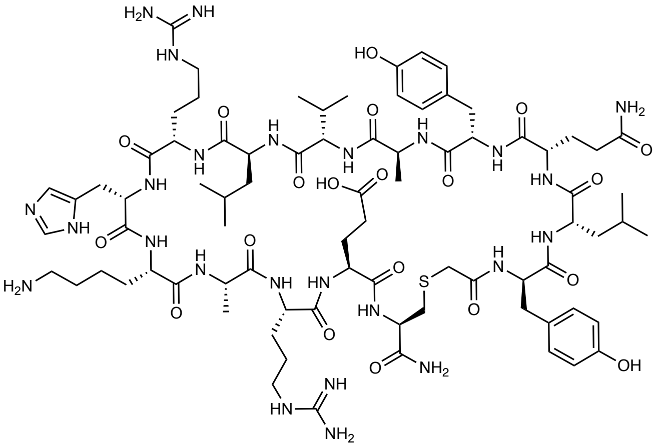

The peptide (50 µmol) was synthesized by general procedure **A** (Oxyma Pure was omitted from the terminal coupling of chloroacetic acid). The peptide was then cleaved from resin by treatment with 87.5:5:5:2.5 v/v/v/v trifluoroacetic acid/triisopropylsilane/$H_2O$/EDT for 2 h. The cleave solution was collected, dried to ~1 mL under $N_2$ flow and the peptide product precipitated from $Et_2O$ (2 x 40 mL) and collected by centrifugation. The crude peptide was then purified by preparative RP-HPLC (0 vol.% $CH_3CN$ + 0.1 vol.% TFA for 5 min, then 5-25 vol.% $CH_3CN$ + 0.1 vol.% TFA over 60 min). Fractions containing the linear peptide were combined and lyophilized. The peptide was then cyclized by general procedure **B**. The reaction was concentrated under $N_2$ flow before being purified by preparative RP-HPLC (0 vol.% $CH_3CN$ + 0.1 vol.% TFA for 5 min, then 5-25 vol.% $CH_3CN$ + 0.1 vol.% TFA over 60 min) to afford **1** as a white solid (3.8 mg, 3%). **$R_{t\ 214nm}$:** 22.60 min. (1 to 40 vol.% $CH_3CN$+ 0.1 %vol TFA over 30 min). **LR-MS (+ESI):** m/z = 1789.45 $[M+H]^+$, 895.35 $[M+2H]^{2+}$, 597.20 $[M+3H]^{3+}$, 448.15 $[M+4H]^{4+}$.

**1-R12A**

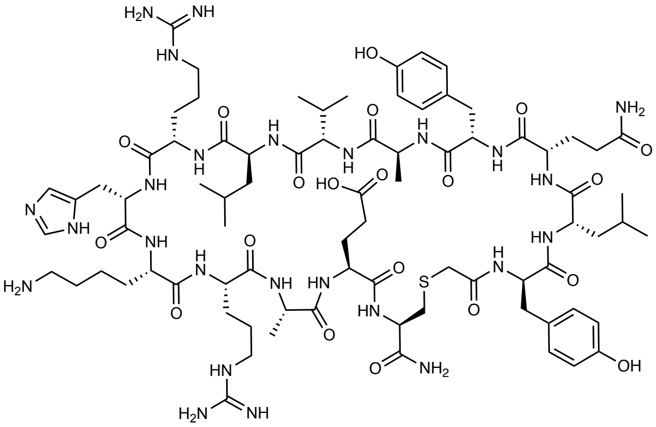

The peptide (50 µmol) was synthesized by general procedure **A** (Oxyma Pure was omitted from the terminal coupling of chloroacetic acid). The peptide was then cleaved from resin by treatment with 87.5:5:5:2.5 v/v/v/v trifluoroacetic acid/triisopropylsilane/$H_2O$/EDT for 2 h. The cleave solution was collected, dried to ~1 mL under $N_2$ flow and the peptide product precipitated from $Et_2O$ (2 x 40 mL) and collected by centrifugation. The crude peptide was then purified by preparative RP-HPLC (0 vol.% $CH_3CN$ + 0.1 vol.% TFA for 5 min, then 5-30 vol.% $CH_3CN$ + 0.1 vol.% TFA over 60 min). Fractions containing the linear peptide were combined and lyophilized. The peptide was then cyclized by general procedure **B**. The reaction was concentrated under $N_2$ flow before being purified by preparative RP-HPLC (0 vol.% $CH_3CN$ + 0.1 vol.% TFA for 5 min, then 5-25 vol.% $CH_3CN$ + 0.1 vol.% TFA over 60 min) to afford **1** as a white solid (16.3 mg, 14%). **$R_{t\ 214nm}$:** 22.20 min. (1 to 40 vol.% $CH_3CN$+ 0.1 %vol TFA over 30 min). **LR-MS (+ESI):** m/z = 1789.35 $[M+H]^+$, 895.25 $[M+2H]^{2+}$, 597.20 $[M+3H]^{3+}$, 448.15 $[M+4H]^{4+}$.

**1-E13A**

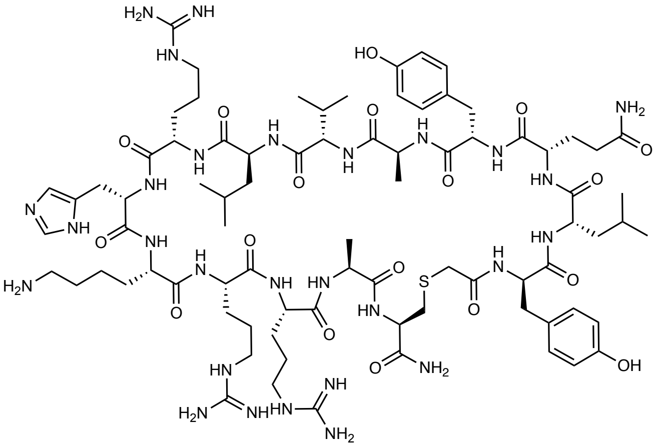

The peptide (50 µmol) was synthesized by general procedure **A** at 75 °C (Oxyma Pure omitted from the terminal coupling of chloroacetic acid). The peptide was then cleaved from resin with 87.5:5:5:2.5 v/v/v TFA/triisopropylsilane/$H_2O$/EDT (5 mL, 2 h). The cleave solution was collected, dried to ~1 mL under $N_2$ flow and the peptide product precipitated from $Et_2O$ over dry ice (2 x 40 mL) and collected by centrifugation. The peptide was then purified by preparative RP-HPLC (0 vol% $CH_3CN$ + 0.1 vol% TFA for 5 min, then 5-25 vol% $CH_3CN$ + 0.1 vol% TFA over 60 min). Fractions containing the linear peptide were combined and lyophilized. The lyophilized peptide was then cyclized by general procedure **B** before being dried under $N_2$ flow and purified by preparative RP-HPLC (0 vol% $CH_3CN$ + 0.1 vol% formic acid for 5 min, then 5-30 vol% $CH_3CN$ + 0.1 vol% formic acid over 60 min, 15 mL/min, XBridge® C18, 300 Å, 19 x 150 mm). The appropriate fractions were combined and lyophilized to afford **1-E13A** as a white amorphous solid (7.0 mg, 6%). **$R_{t\ 214nm}$:** 21.88 min. (1 to 40 vol% $CH_3CN$+ 0.1 %vol TFA over 30 min, 1 mL/min, 40 ºC, XBridge® BEH C18, 4.6 x 250 mm, 300 Å, 5 µM). **LR-MS (+ESI):** m/z = 1816.55 $[M+H]^+$, 908.90 $[M+2H]^{2+}$, 606.35 $[M+3H]^{3+}$, 454.95 $[M+4H]^{4+}$.

**Se-1**

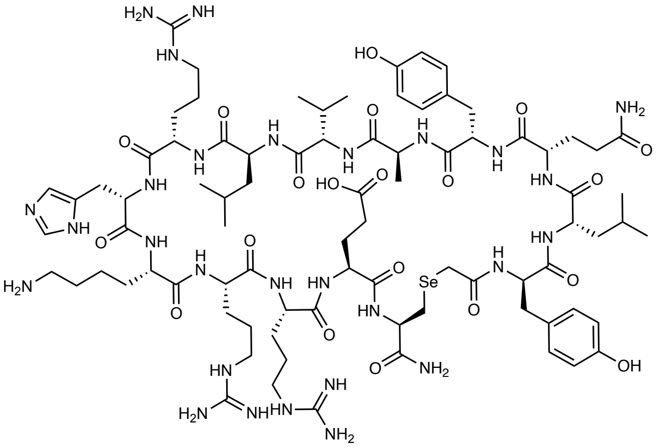

Rink Amide resin (40 μmol, 0.488 mmol.g$^{-1}$) was treated with a solution of 20 vol.% piperidine in DMF (3 mL) for 2 x 5 min, then washed with DMF (5 x 4 mL), $CH_2Cl_2$ (5 x 4 mL) and DMF (5 x 4 mL) before Fmoc-Sec(PMB)-OH (51 mg, 100 μmol) was loaded in the presence of *N-N'*-diisopropylcarbodiimide (15.6 μL, 100 μmol) and Oxyma Pure (14.2 mg, 100 μmol). The resin was then washed with DMF (5 x 4 mL) $CH_2Cl_2$ (5 x 4 mL) and DMF (5 x 4 mL) before treatment with 5 vol.% $Ac_2O$ with 10 vol.% *i*-$Pr_2NEt$ in DMF for 5 min followed by washing with DMF (5 x 4 mL), $CH_2Cl_2$ (5 x 4 mL) and DMF (5 x 4 mL). The remainder of the peptide was then synthesized by general procedure **A** at 50 °C (Oxyma Pure was omitted from the terminal coupling of chloroacetic acid). The peptide was then cleaved from resin by treatment with 87.5:5:5:2.5 v/v/v/v trifluoroacetic acid/triisopropylsilane/$H_2O$/EDT for 2 h. The cleave solution was collected, dried to ~1 mL under $N_2$ flow and the peptide product precipitated from $Et_2O$ (2 x 40 mL) and collected by centrifugation. The crude peptide was then purified by preparative RP-HPLC (0 vol.% $CH_3CN$ + 0.1 vol.% TFA for 5 min, then 0-30 vol.% $CH_3CN$ + 0.1 vol.% TFA over 60 min). Fractions containing the linear peptide were combined and lyophilized. The peptide was then cyclized by general procedure **B** in the presence of TCEP (10 mM). The reaction was lyophilized before being purified by RP-HPLC (0 vol.% $CH_3CN$ + 0.1 vol.% TFA for 5 min, then 0-40 vol.% $CH_3CN$ + 0.1 vol.% TFA over 60 min) to afford **1** as a white solid (6.2 mg, 6%). **$R_t$ $_{214nm}$:** 22.23 min. (1 to 40 vol.% $CH_3CN$+ 0.1 vol.% TFA over 30 min). **LR-MS (+ESI):** m/z = 961.40 $[M+2H]^{2+}$, 641.25 $[M+3H]^{3+}$, 481.20 $[M+4H]^{4+}$.

**Authentic standard of cleaved 1**

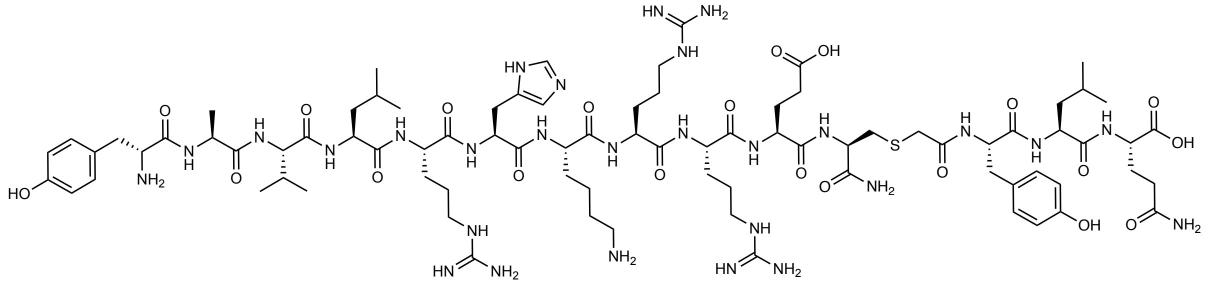

The peptide was synthesized in two fragments:

**Fragment 1:** The sequence YAVLRHKRREC was synthesized by general procedure **A** at 50 °C. The resin-bound peptide was washed with DMF (3 x 4 mL), DCM (3 x 4 mL) and DMF (3 x 4 mL), prior to Fmoc-deprotection of the N-terminus with 20 vol.% piperidine in DMF (4 mL, 2 x 10 min, rt.). The resin-bound peptide was washed with DMF (3 x 4 mL) and DCM (5 x 4 mL), then cleaved from bead with TFA/TIS/$H_2O$/EDT (87.5/5/5/2.5 vol.%, 5 mL, 2 h, rt). The cleave solution was collected, dried to ~1 mL under $N_2$ flow and the peptide product precipitated from $Et_2O$ over dry ice (2 x 40 mL) and collected by centrifugation. The peptide was then purified by preparative RP-HPLC (0 vol.% $CH_3CN$ + 0.1 vol.% TFA for 5 min, then 0-40 vol.% $CH_3CN$ + 0.1 vol.% TFA over 60 min). Fractions containing the linear peptide were combined and lyophilized to afford **Fragment 1** as a white solid (45.0 mg, 40%).

**Fragment 2:** 2-chlorotritylchloride resin (Mimotopes, 50 µmol, 1.12 mmol $g^{-1}$) was loaded with Fmoc-Gln(Trt)-OH (8 equiv.) and *i*-$Pr_2$NEt (8 equiv) for 16 h before being drained and treated with 17:2:1 v/v/v $CH_2Cl_2$:MeOH:*i*-$Pr_2$NEt (5 mL) for 30 mins. The resin was then washed with DMF (4 x 5 mL), $CH_2Cl_2$ (4 x 5 mL) and DMF (4 x 5 mL). The resin was then treated with a solution of piperidine (20 vol.%) in DMF (2 x 5 mL, 5 min each). The resin was then washed with DMF (4 x 5 mL), $CH_2Cl_2$ (4 x 5 mL) and DMF (4 x 5 mL). The sequence IAc-yLQ-OH was then elongated by coupling Fmoc-amino acid (200 µmol), *N*-*N'*-diisopropylcarbodiimide (31.2 µL, 200 µmol) and Oxyma Pure (28.4 mg) in DMF (4 mL, 2 h, rt.). Following each coupling, the resin-bound peptide treated with 10 vol.% $Ac_2O$/pyridine (2 x 5 min, rt.), washed, and then treated with 20 vol.% piperidine in DMF (2 x 10 min). Iodoacetic acid (37 mg, 200 µmol) was then coupled *N*-*N'*-diisopropylcarbodiimide (31.2 µL, 200 µmol) and Oxyma Pure (28.4 mg) in DMF (4 mL, 2 h, rt.). The resin-bound peptide was washed with DMF (3 x 4 mL) and DCM (5 x 4 mL), before resin cleavage with 90:5:5 v/v/v TFA/TIS/$H_2O$ (5 mL, 2 h, rt). The cleave solution was collected, dried to ~1 mL under $N_2$ flow and the peptide product precipitated from $Et_2O$ over dry ice (2 x 20 mL) and collected by centrifugation. The crude peptide pellet was then air dried and used without further purification.

**Authentic standard of cleaved 1: Fragment 1** (1.0 eq., 11.4 mg, 5.39 μmol) and **Fragment 2** (1.2 eq., 3.9 mg, 6.61 μmol) were dissolved in 1:1 v/v $CH_3CN/H_2O$ (1 mL) and *i*-$Pr_2NEt$ (5 vol.%) added. The reaction was stirred for 1 h at rt. The product was then isolated by preparative RP-HPLC (0 vol.% $CH_3CN$ + 0.1 vol.% TFA for 5 min, then 0-40 vol.% $CH_3CN$ + 0.1 vol.% TFA over 80 min, 15 mL/min, XBridge® C18, 300 Å, 19 x 150 mm). The appropriate fractions were combined and lyophilized to afford the product as a white solid (4.65 mg, 33%). **$R_{t\ 214nm}$:** 14.61 min. (1 to 50 vol.% $CH_3CN/H_2O$ + 0.1 vol.% formic acid over 30 min, 60 ºC) **LR-MS (+ESI):** m/z = 946.90 $[M+2H]^{2+}$, 631.60 $[M+3H]^{3+}$, 473.95 $[M+4H]^{4+}$.

9 The PyMOL Molecular Graphics System, Version 2.0 Schrödinger, LLC.
